# An antiviral self-replicating molecular heterotroph

**DOI:** 10.1101/2020.08.12.248997

**Authors:** Anastasia Shapiro, Alexander Rosenberg, Adva Levy-Zamir, Liron Bassali, Shmulik Ittah, Almogit Abu-Horowitz, Ido Bachelet

## Abstract

We report the synthesis of a molecular machine, fabricated from nucleic acids, which is capable of digesting viral RNA and utilizing it to assemble additional copies of itself inside living cells. The machine’s body plan combines several parts that build upon the target RNA, assembling an immobile, DNA:RNA 4-way junction, which contains a single gene encoding a hammerhead ribozyme (HHR). Full assembly of the machine’s body from its parts enables the subsequent elongation of the gene and transcription of HHR molecules, followed by HHR-mediated digestion of the target molecule. This digestion converts the target to a building block suitable for participation in the assembly of more copies of the machine, mimicking biological heterotrophy. In this work we describe the general design of a prototypical machine, characterize its activity cycle and kinetics, and show that it can be efficiently and safely delivered into live cells. As a proof of principle, we constructed a machine that targets the *Autographa californica* multicapsid nucleopolyhedrovirus (AcMNPV) GP64 gene, and show that it effectively suppresses viral propagation in a cell population, exhibiting predator/prey-like dynamics with the infecting virus. In addition, the machine significantly reduced viral infection, stress signaling, and innate immune activation inside virus-infected animals. This preliminary design could control the behavior of antisense therapies for a range of applications, particularly against dynamic targets such as viruses and cancer.

## Introduction

The discovery and synthesis of the immobile DNA junction by Seeman, nearly 4 decades ago^1^, emerged as a cornerstone of DNA nanotechnology, an expanding field with unique technological potential. The immobile junction enabled, for the first time, programming and control of the spatial positioning of matter at a single DNA base resolution, or approximately 3.5 Å^2–4^. This discovery was followed by a range of works which explored various geometries, scales, construction methods, application, and behaviors^5–18^.

The term DNA nanotechnology could be used to define a more loose group of phenomena and applications, based on nucleic acid functions beyond the context of carrying and manipulating genetic information, both artificial and naturally occurring. Some of these functions are molecular computation^19–21^, ribo- and deoxyribozymes^22–26;^ aptamers^27, 28^; and toehold-mediated strand displacement reactions^29–32^. Such systems have been recently implemented in combination in several of the works referenced above in order to obtain complex functions and high level of logical control. Of particular impact in this progress has been the existence of high-quality computer-aided design (CAD) tools to program, simulate, and reliably analyze these systems^33–37^.

The ability to integrate various aspects of DNA nanotechnology in simple and relatively-tractable designs has highlighted a fascinating and long-standing aim - the design and fabrication of nanoscale robots, an achievement envisioned by prominent 20th century thinkers like Richard Feynman, John Von Neumann, and Isaac Asimov^38–41^. Indeed, several diverse designs of true robots - defined here as **automata** which are capable of **sensing** environmental cues and **responding** by actuation, and are guided by a non-random set of rules or **logic** - based on DNA, have been recently described^42–50^. DNA-based molecular machines offer a unique approach to challenges in domains such as manufacturing and medicine, which would be extremely hard or even impossible to tackle by conventional macroscopic or chemical means.

Perhaps the most remarkable property of such machines could be the ability to self-replicate. A hallmark of living organisms, self-replication has been synthetically emulated in several systems and scales^51, 52^, with some successful designs stemming directly from DNA nanotechnology^53–55^. The construction of self-replicating agents with direct application in a realistic setting is highly desired and has been explored in the past^56–58^. A particular field in which self replication could provide a crucial advantage over existing solutions is therapeutics, especially against replicating and evolving targets such as viruses, bacteria, and tumor cells. Conventional drugs are typically administered constantly until the pathogen is either eliminated or becomes resistant. On the other hand, self-replicating therapeutics could be administered at extremely low starting doses, or high doses of raw, inactive building blocks, and then replicate. Thus, they could track the target population until its elimination, and eventually be cleared from the system. Existing examples of self-replicating therapeutics include oncolytic viruses^59^ and RNA vaccines^60^, although the latter is essentially a delivery system for vaccines rather than a self-replicating therapeutic agent. Both examples are modified versions of natural viruses. The ability to design an entirely synthetic, self-replicating therapeutic machine remains a significant technological challenge.

In this work, a preliminary working prototype of such a system is reported. We describe a machine, or automaton, assembled from DNA and RNA, which incorporates structural and functional principles of DNA nanotechnology. The automaton is designed as an immobile, DNA:RNA 4-way junction assembled from 3 parts, and contains a single gene which encodes a hammerhead ribozyme (HHR). Full assembly of the automaton’s body from its parts enables the subsequent transcription of HHR molecules and HHR-mediated digestion of the target molecule. This digestion renders the target suitable for participation in the assembly of more copies of the automaton, essentially mimicking biological heterotrophy. The starting material for self-replication, and for the subsequent “lifecycle” of the automaton, is a small seed population of fully-assembled machines, and a non-limiting amount of the separate DNA parts required for the assembly of new copies of the machine.

The automaton we describe here was inspired by pathogenic viruses and viroids, and is designed to be delivered into and operate inside living, virus-infected cells. Inside the cells, the automaton’s goal is to counter viral infection by utilizing viral RNA as “food”. As a proof of principle towards therapeutic application, we designed an automaton that targets *Autographa californica* multicapsid nucleopolyhedrovirus (AcMNPV) RNA, characterized its assembly chemistry and activity *in-vitro*, and calibrated the precise stoichiometric packaging of its parts, along with the enzymatic machinery required for HHR transcription from this particular machine design, into therapeutically-compatible formulations. This machine effectively suppressed virus ability to create infective virions in AcMNPV-infected cells, exhibiting predator/prey dynamics with the infecting virus. Moreover, it significantly reduced viral infection in terms of potent virions titer, as well as rate of infection spreading, stress signaling, and innate immune activation inside AcMNPV-infected adult insects.

The design reported here is modular, and can be adapted rapidly to counter new targets, for example viruses once their genome has been sequenced. In addition, it could lead to new types of biologically-inspired, programmable agents for a range of therapeutic applications. Interestingly, this machine highlights complex anatomy and a hypothetical, rudimentary mechanism for predation, plausible features of molecular life forms from the RNA world.

## Results

**Fig. 1A** describes a schematic design of the machine (or automaton). The automaton’s body plan consists of 3 parts: a long DNA oligonucleotide (L), a short DNA oligonucleotide (S), and an RNA base unit (R), that together build a 4-way junction (the design of this junction and simulations of its assembly are detailed in **Supplementary notes 1 & 2, Fig. S1-S2, & Table S1**). In this schematic, R is depicted as consisting of two subunits, but in reality R is a single part formed from a variety of possible target molecules, as shown and discussed below. Parts L and S are designed such that their hybridization is supported only in the presence of R **(Fig. 1B)**. Fully assembled automata were visualized by atomic force microscopy (AFM) using DNA origami rectangles as reference objects **(Fig. 1C, Supplementary note 3, Fig. S7)**. Complete automaton assembly occurs on the order of 1 h **(Fig. 1D)**. The assembly reaction correlated with stoichiometry of parts L, S, and R, and at a 1:1:1 stoichiometry was measured to be 63±10% **(Fig. 1E)**. Automata were stable on the shelf for at least 29 days **(Supplementary note 1, Fig. S3)**.

**Figure 1.**
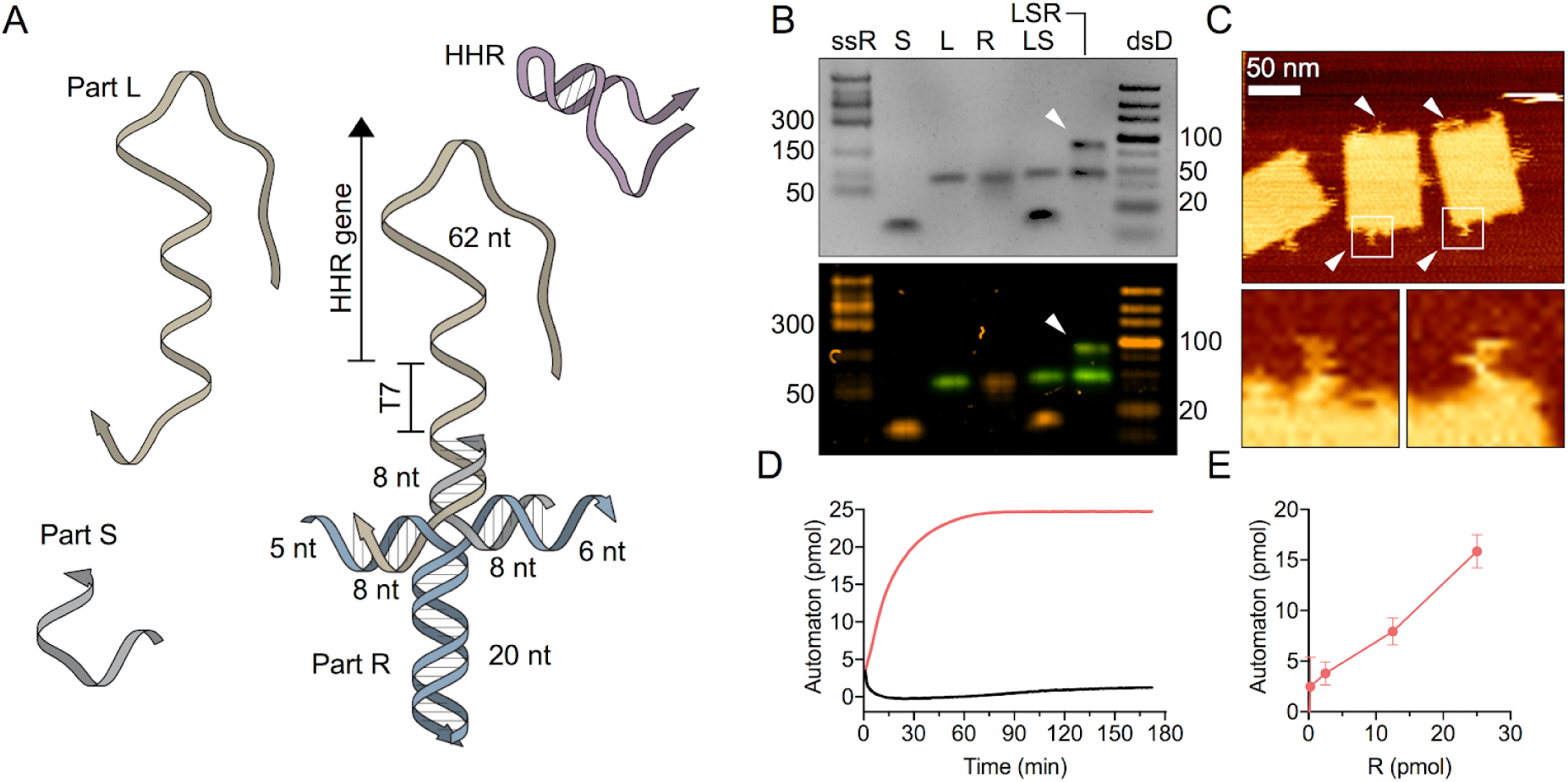
Design and assembly of the molecular machine. **A,** Design schematic. The machine’s body plan combines three parts: L (beige), S (grey), and R (blue). Region lengths are specified in nucleotides (nt). Locations of the T7 promoter and the hammerhead ribozyme (HHR) genes are shown. The transcribed HHR molecule itself is shown in purple. **B,** assembly of the machine from its parts, analyzed by gel electrophoresis. Bottom panel shows fluorescence scanning of the gel, gel is stained with ethidium bromide (orange), part L is fluorescently labeled (green). S, L, R denote the individual parts. LS and LSR and mixtures of the respective parts. White arrowhead points at assembled machines. Lower band at LSR represents excess parts L and S (ssR, single-stranded RNA ladder; dsD, double-stranded DNA ladder; selected ladder reference points are shown on either side, numbers in base/base pairs). **C,** AFM scans of assembled machines mounted on DNA origami rectangles. White arrowheads point at machines. White frames are magnified in the bottom panels. Scale bar = 50 nm. **D,** machine assembly kinetics (red) measured by fluorometry. Black curve represents LS. **E,** machine assembly efficiency, showed as a function of the quantity of R; quantities of L and S were maintained constant at 25 pmol.

Upon assembly of the automaton, part S undergoes elongation by DNA polymerase, using L as template **(Fig. 2A, 2B)**, creating a dsDNA region which functions as a gene controlled by the T7 promoter and encodes a hammerhead ribozyme (HHR). T7 RNA polymerase next transcribes the gene to synthesize HHR molecules **(Fig. S8)** which digests a specific RNA target. Digestion of this target molecule is the key driver of automaton self-replication. It is essentially a metabolic reaction that converts the target molecule into a building block capable of participating in the assembly of additional copies of the automaton from its parts L and S. This mechanism is analogous to biological heterotrophy, which is defined as the nutritional type that utilizes an exogenous source of carbon as food^61^. However, in our design it was imperative to validate that the automaton is able to utilize its target only after metabolizing it. For this reason, we designed a specific RNA target composed of a ubiquitous structural stem-loop motif, such that digestion converts the molecule from a stem-loop structure into a stem with two open ends, which comprises the part designated as R **(Fig. 2C)**. We used ribosoft^62^ in the design of HHR molecules specific to target RNA **(Fig. 2D)**. Binding of HHR to its target is mediated by Watson-Crick pairing via two separate regions of the HHR as previously described^62^. Various loop sizes within the target RNA motif were tested to highlight an optimal structure that enables efficient HHR activity without supporting an L-S interaction without digestion. A length allowing 8 nt in addition to the regions complementary to L and S was found to represent this optimum **(Fig. 2E)**. In the present design, a single automaton synthesized HHR molecules at a rate of approximately 30 molecules/h **(Fig. 2F, 2G)**. The transcribed HHR molecules efficiently digested the target RNA, converting it to part R **(Fig. 2H)**. We calculated a target digestion rate constant of approximately *k* ∼ 0.14 min^-1^ at 37 °C, which is lower than the *k* ∼ 1 min^-1^ observed for linear RNA^63^, and an EC_50_ of approximately 1 mol HHR per 5 mol target RNA **(Fig. 2I, 2J, Supplementary note 4, Fig. S9-S10)**. The entire “life-cycle” of the machine, starting from the low machine/high target state and ending with the high machine/low target one was visualized by gel electrophoresis **(Fig. 2K)**.

**Figure 2.**
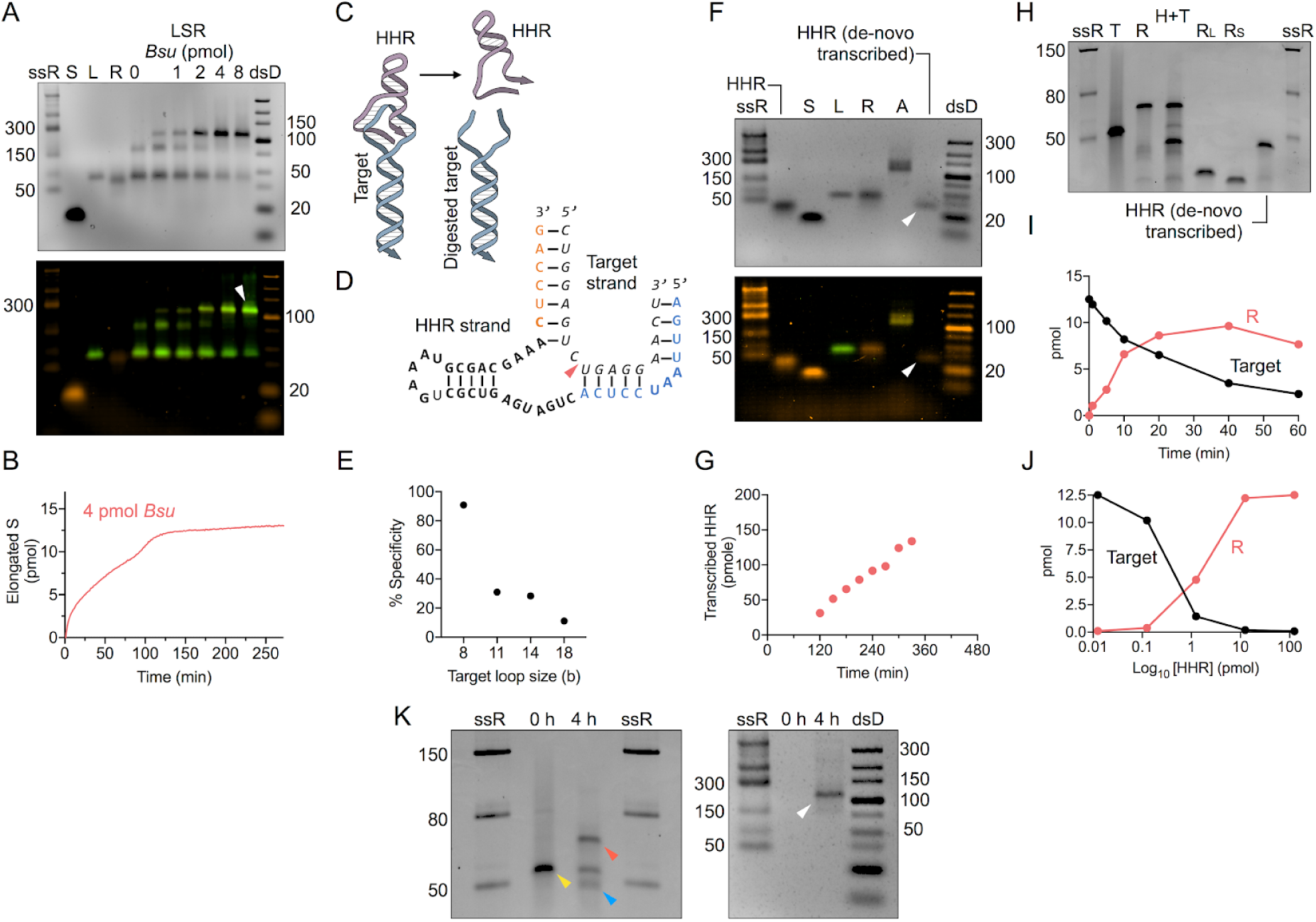
Characterization of machine activity and self-replication. **A,** elongation of part S by varying quantities of *Bsu* polymerase. Bottom panel shows fluorescence scanning of the gel, gel is stained with ethidium bromide (orange), part L is fluorescently labeled (green). S, L, R denote the individual parts. LSR denotes mixture of all parts. (ssR, single-stranded RNA ladder; dsD, double-stranded DNA ladder; selected ladder reference points are shown on either side, numbers in bases/base pairs; unlabeled lane denotes 0.5 pmol *Bsu*). **B,** kinetics of the elongation of part S by *Bsu* polymerase (representative quantity of 4 pmol), measured by fluorescence. **C,** schematic diagram of target digestion by HHR. **D,** schematic of the HHR-target complex. HHR regions complementing the target are shown in orange and blue. Red arrowhead pointing to the site of digestion in the target strand. **E,** Specificity of automaton formation, expressed as the ratio of automata (pmol) formed on digested vs. undigested targets with varying loop sizes. **F,** transcription of HHR from assembled automata, analyzed by gel electrophoresis. Bottom panel shows fluorescence scanning of the gel, gel is stained with ethidium bromide (orange), part L is fluorescently labeled (green). S, L, R denote the individual parts, A denotes assembled automata. White arrowhead points at *de-novo* transcribed HHR. **G,** HHR transcription kinetics from ∼ 50 pmol automata, measured by HPLC. **H,** target digestion by *de-novo* transcribed HHR. T, R denote target and R part, respectively. H+T is the target-HHR mixture. R(L) and R(S) denote the parts of R that hybridize with parts L and S, respectively. **I,** kinetics of target digestion by HHR, measured by gel electrophoresis. **J,** dose response of target digestion by HHR, measured by gel electrophoresis. **K,** automaton predatory action and self-replication. Representative gel electrophoresis of the reaction sampled at t=0 and 4 h. Fractions containing T, R, and automata were isolated by HPLC and run on separate gels for visualization clarity. Arrowheads denote as follows. Left panel: yellow, target. Red, digested target (part R). Blue, HHR. Right panel: white, assembled automata.

In addition to this automaton, we designed and tested several alternative automata to explore design flexibility and additional features **(Supplementary note 5, Fig. S11-S17, Table S2 & Table S3)**. For example, we incorporated a “sensory organ” based on toehold-mediated strand displacement. This part is designed to respond to the presence of the target by structural remodeling of the HHR gene region and enabling its transcription, which is otherwise disabled. Such a feature could provide an additional layer of control and specificity to the automaton’s activity, triggering its operation only in the presence of its target.

We next validated the automaton’s compatibility for delivery into live cells and its stability therein. DNA and RNA nanostructures have been stably generated and studied in live cells^64, 65^, and recent works have reported the successful delivery of DNA nanostructures into cells^66, 67^ as well as packing DNA nanostructures in liposomes^68, 69^, which improves their *in-vivo* stability. Our experimental design aimed to reinforce these observations. We first studied several variants of the automaton, in which parts L and S were modified with either locked nucleic acids (LNA) or 2-O-methoxyethyl RNA (2-MOE) at various lengths along the regions complementary to part R (3, 5, and 7 nt). We found that both modifications were effective in protecting the assembled automaton from RNase H, with stability correlating with modification length **(Fig. 3A, 3B, Fig. S4)**. The modified automata were also resistant to the junction resolvases RuvAB and RuvABC **(Fig. 3C)**. These experiments also confirmed that the modifications did not alter oligonucleotide assembly behavior and specificity **(Fig. 3A-C**, “LS” traces); rather, they significantly increased junction formation efficiency **(Fig. S3)**.

**Figure 3.**
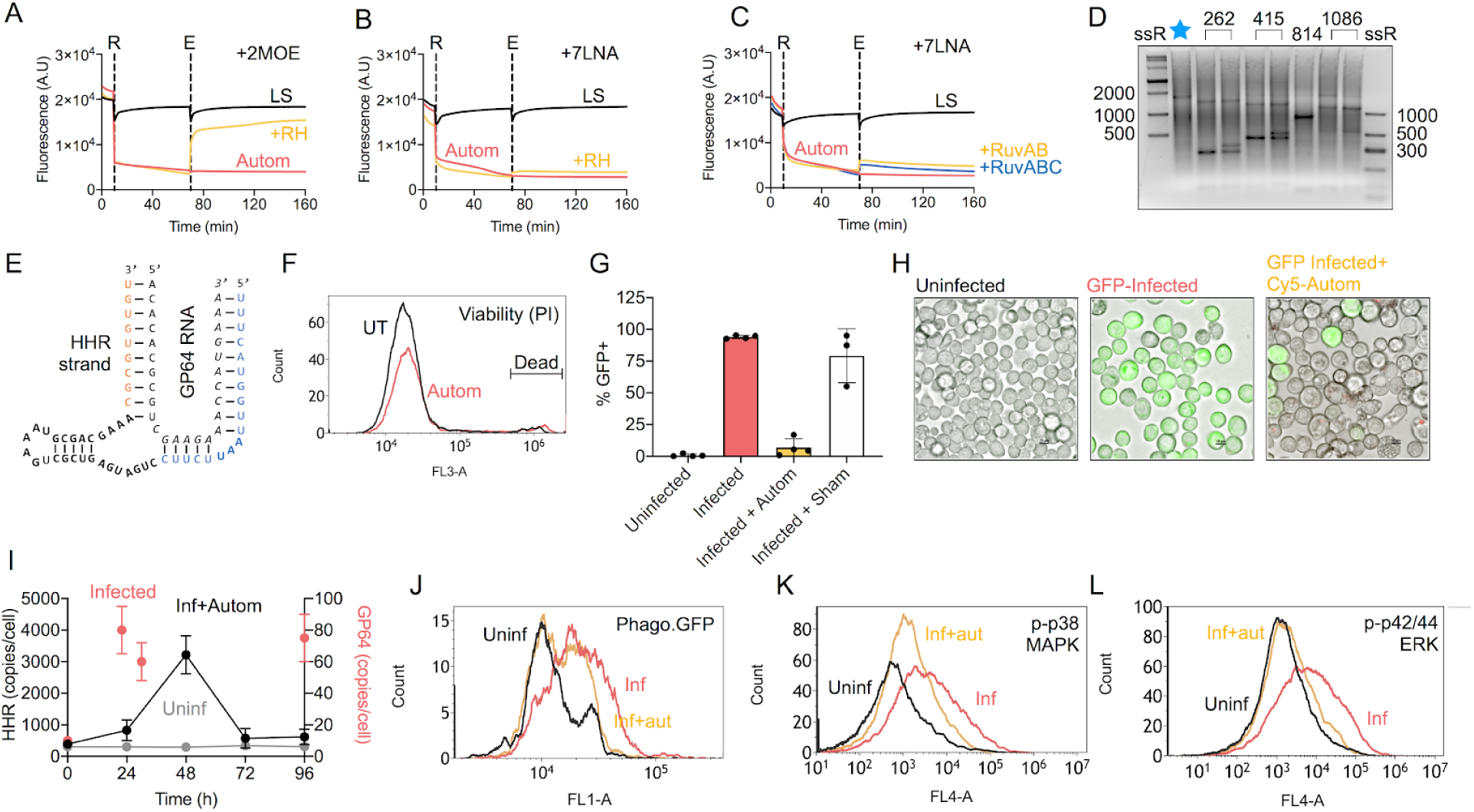
Automaton performance in infected cells and animals. **A-C**, stability measurements of automata in the presence of RNase H and the junction resolvases RuvAB/RuvABC. Parts S and L were labeled with fluorophore and quencher, respectively, such that automaton dissociation was observed as an increase in fluorescence. A mix of parts L and S (LS), which does not assemble without part R, was used as positive control for dissociation. Automata (Autom) were used as negative control. Dashed lines, R = addition of part R (in all but LS samples), E = addition of enzyme(s). 2MOE & 7LNA, tested modifications (see **Supplementary note 1)**. **D**, *in-vitro* activity of HHR on AcMNPV GP64 RNA. Numbers above image denote target site position (in bases from 5’). Blue star marks untreated GP64 RNA. **E**, chosen GP64-targeting HHR. **F**, viability analysis by propidium iodide (PI) of Sf9 cells treated with 250 pmol automata package for 4 days. UT, untreated cells; Autom, automata-treated. Marker includes PI^+^ (dead) cells. **G**, efficacy of automata in Sf9 cells infected with GFP-AcMNPV. Autom, automata. Analysis was performed on 4 biological repeats. **H**, representative microscope images of Sf9 cells infected with GFP-AcMNPV, and treated with Cy5-labeled automata (autom). Bar = 10 um. **I**, HHR transcription dynamics along the course of Sf9 infection with GFP-AcMNPV, relative to GP64 RNA quantity per cell. Cells at specific time points were fixed and HHR molecules were counted by quantitative flow cytometry. GP64 RNA was counted by qPCR. Uninf, uninfected and treated with automata. Inf, infected and treated. **3J-L**, Flow cytometric analysis of specific parameters in cells isolated from animals infected with AcMNPV and treated with automata. Cells were fixed and stained with Alexa Fluor 647-labeled phospho-specific antibodies, with a labeled anti-Akt (protein kinase B) antibody used as protein content/negative control. Histograms are from representative animals. Uninf, uninfected. Inf, infected. Inf+aut, infected and treated with automata. **J**, plasmatocyte-phagocytosed GFP (Phago.GFP). **K**, p38 MAPK phosphorylation. **L**, p42/44 MAPK (ERK1/2) phosphorylation.

In order to demonstrate automaton activity inside living cells, we designed variants that specifically target the genes of an infecting virus. As a model virus, we used AcMNPV to infect Sf9 cells in culture **(Supplementary notes 6, Fig. S18-S19)**. As the target gene we chose GP64, the critical envelope glycoprotein mediating host cell recognition and entry^70^. Several HHR molecules were designed against specific target sites in GP64, and were tested on GP64 RNA *in-vitro* **(Fig. 3D)**. The most effective molecule **(Fig. 3E)** was used as a design constraint for LNA-modified, GP64-targeting automata that exhibited efficient assembly **(Supplementary note 7, Fig. S24-S26, Table S4).** These automata were not toxic to cells during 4 days of exposure at a high dose (250 pmol) **(Fig. 3F)**.

AcMNPV infection was next set up in cells by mixing a population of Sf9 cells infected at an MOI of 1, with uninfected cells at a ratio of 1:99, simulating a viral infection at it’s initial stage **(Supplementary note 6)**. The entire population was treated with a single dose of automata packaged in either of several tested liposomal formulations **(Supplementary note 6, Fig. S21-S22)**. Cell populations were analyzed 96 h post infection by flow cytometry, looking at GFP expression. Compared with the infected untreated population, which contained 95±2% GFP^+^ cells, the population treated once with GP64-targeting automata showed a striking reduction of infection, to between 10±7% GFP^+^ cells (*P*<10^-4^) **(Fig. 3G, 3H, Supplementary note 7)**. The sham-treated group showed no difference from the infected group. In order to observe the temporal dynamics of automata activity relative to the target virus, cell populations were sampled at intervals of 24 h from 0 to 96 h, and the *de-novo* transcribed HHR molecules per cell were counted using *in-situ* hybridization quantitative flow cytometry. Interestingly, HHR transcription produced an approximate average of 800 copies per cell at 24 h (*P*=0.002), and to approximately 3000 copies per cell at 48 h (*P*<10^-4^). No HHR molecules were significantly observed above the background at 72 and 96 h, and HHR counts did not significantly change with time in uninfected cells treated with equal amounts of automata. Interestingly, the dynamics of HHR appeared to track that of GP64 RNA as measured by qPCR with an approximate lag of 24 h **(Fig. 3I)**. This tracking behavior suggests that the *de-novo* transcription of HHR responds to the cellular viral load.

Lastly, we investigated the potential effect of the GP64-targeting automata on viral load and consequently on the innate immune response to AcMNPV infection in adult insects. To this end, we infected adult *Blaberus discoidalis* with AcMNPV by a systemic injection of 4×10^5^ cells, out of which 40,000 were infected and expected to release approximately 100 virions per cell over the course of the next 48 h. Hemolymph was extracted from the insects at 2.5 d post infection, and flow cytometry was used to measure the fluorescence of GFP molecules phagocytosed by resident plasmatocytes, as well as the phosphorylation states of p38 MAPK and ERK, which are upregulated in stress and innate immune activation in insects in response to viral infection^71^. One group of animals received GP64-targeting automata by injection at t = 0. Animals injected with infected cells exhibited increased plasmatocyte GFP compared to animals injected with healthy cells (*P*<10^-4^); however, treatment with GP64-targeting automata reduced GFP levels significantly (*P*<10^-4^) **(Fig. 3J)**, indicating a reduction in viral load. Automata-treated animals also showed near-complete reversal of stress and innate immunity signals **(Fig. 3K)**. Particularly, p38 MAPK and ERK phosphorylation were significantly reversed (phospho-p38 MAPK, *P*<10^-4^; phospho-ERK, *P*<10^-4^) **(Fig. 3L)**, indicating that both the innate immune response and the systemic stress were significantly reduced as a consequence of viral load reduction by the GP64-targeting automata.

## Discussion

In this work we describe a working prototype of a synthetic molecular automaton, which digests viral RNA and utilizes it as “food” for assembling additional copies of itself inside living cells. This combination of heterotrophy and self-replication, both inspired by living organisms, with the automaton’s nucleic acid-based architecture, is unique among therapeutic agents, and provides several benefits.

Bacteria, viruses, and cancer represent some of the most difficult and devastating health challenges humanity has ever faced. As an example, the global cost of the current COVID-19 pandemic^72^ has been estimated by the Asian Development Bank at up to 4 Trillion USD^73^, which in modern USD values is comparable to the cost of World War II. Excluding COVID-19, there were at least 5 major viral pandemics during the decade 2009-2019, including influenza, SARS, and Ebola virus, each causing damage estimated in multi-billion USD. The efforts to discover and repurpose therapies for COVID-19 include more than 120 examples of vaccines alone^74^, and are still ongoing. Like other silencing therapeutics, the automaton (specifically its HHR core) can be rapidly designed *ab-initio* and screened against pathogen RNA once it is sequenced, which could enable a rapid response to future outbreaks.

The tracking behavior of the automaton, expanding and decaying depending on the concentration of its target, is a unique pharmacological advantage. It makes it possible to designate the automaton’s building blocks as an inactive prodrug, enabling their safe delivery in large amounts. Only in the presence of its “prey” (i.e. when the host cell is infected by the target virus), can the automaton assemble and begin replicating. The *in-vivo* stability of the automaton is programmable to an extent, using synthetic modifications in specific base stretches, such as LNA and MOE, as shown in this study. These levels of control and programmability are nearly unmatched in current mainstream therapeutics.

Like many viruses, which package and carry the protein machinery required for their replication in the destination cell, and like experimental CRISPR/Cas-based therapeutic formulations, the automaton is delivered along with its protein machinery. For purposes of this proof-of-principle study we used off-the-shelf enzymes. However, future designs could include human enzymes isolated or optimized specifically for the automaton, although this study has shown that delivery of the automaton packages is both efficient and non-toxic in insect cells and adult insects. It may be even possible to design automata that entirely utilize the endogenous cellular machinery, which would simplify its delivery and formulation.

Interestingly, several structural and functional features of the automaton could highlight plausible features of molecular life forms from the RNA world^75^. For example, the automaton has multiple parts that compose a complex body plan. This model has been previously suggested as underlying mechanism for structural and functional diversity in the RNA world, since it is assumed that ancient RNA polymerizing ribozymes had generally low processivity, thus likely capable of synthesizing relatively short species^76^ (it is important to note that recent studies described synthetic, single-chain ribozymes with impressive catalytic performance^23, 77, 78^). Second, it contains a metabolic ribozyme, which is capable of breaking down other molecules in order to utilize them as a source of building blocks. Although the terms “molecular ecology” and “molecular predators” have been previously suggested for synthetic systems and in bacterial small RNA networks,^79–81^ the concept is essentially descriptive of specific dynamics, and has not been constructed at a metabolic level. In this narrow sense, the present automaton could emulate realistic predatory behavior.

Third, some designs which we tested only briefly (and are described in **Supplementary note 5**) included a sensory part or “organ”, enabling the automaton to sense its target and respond to it. Sensing of and response to environmental cues such as temperature, light, food, prey, and predators, is a hallmark feature of living organisms, and the chemistry of nucleic acids provides primitive mechanisms for sensory capabilities. A simple example would be based on base pairing in toehold-mediated strand displacement, translating hybridization to actuation. Another mechanism could be the recognition of a 3D structure, similar to aptamers and their targets, with the coupled actuator similar in principle to the concept of structure-switching aptamer-based sensors^82^. An intriguing feature of this type of sensing mechanism, is that it induces structural remodeling of the automaton that exposes the HHR gene to be transcribed, which could be viewed as analogous to chromatin remodeling in higher organisms, in response to various cues. Sensing and response provide an additional layer of control over the automaton, contributing to a less leaky, more stringent system.

Gene silencing and RNA-based therapeutics are still an emerging field of therapeutics, although it can be traced back four decades ago^83, 84^. RNA-based therapeutics offer many advantages over other classes of pharmaceuticals and biologics, among them target flexibility, facile and precisely-defined chemical synthesis, cost effectiveness, shelf stability, very low immunogenicity, and nearly on-demand design. Traditional issues such as *in-vivo* stability and nuclease resistance have been simplified by modern methods for synthesis and base modification. Their rapid design and platforms for evolution have made it possible to tailor oligonucleotides for variable targets in cancer^85^. The automaton described here, and improved versions of it, could be utilized as engines to drive the intracellular synthesis of RNA-based therapeutics such that they track their gene target, as a unique mode of control over the activity of the drug.

Interestingly, nucleic acid-based therapeutics are unique in being very highly “distributable”: once an oligonucleotide-based drug has been designed and its efficacy and safety confirmed, it can be sent digitally as a sequence file and synthesized locally on commercially available synthesizers. We have recently shown using network models^86^ that even when the oligonucleotide therapeutic is much less effective than “chemical” drugs (e.g. a small molecule or antibody), its rapid distribution - compared to shipping via sea and air - consistently enabled us to save more people. Importantly, distributing “chemical” drugs by shipping creates profound inequalities, inevitably leading to some populations receiving more material than needed, while others remain completely deprived of it. In stark contrast, nucleic acid therapeutics could lead to a significantly more equal distribution. In this regard, the current COVID-19 crisis highlights the important of these, traditionally overlooked, advantages of nucleic acid therapeutics.

## Methods

### Oligonucleotides

DNA and RNA oligonucleotides, including any modification, were ordered from Integrated DNA Technologies (IDT), reconstituted to 100 μM with ultrapure, DNase/RNase free water (Biological Industries, Israel), and stored at −20°C. DNA/RNA sequences are listed in **Table S1**.

### Automaton design and assembly

HHR molecules for given target sequences were designed using Ribosoft^62^. Automaton structure was then designed using a custom code ensuring its fit to the cleavage sites and the HHR, as well as compliance with the rules defined by Seeman^1^ for immobile junctions. Automaton structure simulations were carried out in NUPACK^37^ (**Supplementary note 1**). To prevent elongation of part L, a 3’-Phosphate modification was added. Automata were assembled by mixing all components (L, S, R(L) and R(S)) at a 1:1:1:1 ratio in 1× TAE buffer with 5 mM MgCl_2_ at 37° C over 1 h in a Bio-Rad C1000 Touch Thermal Cycler. Final concentration of each part was 1 μM in 25 μL. Structure assembly was verified by gel electrophoresis and fluorescence using internal-FAM labeled part L and 3’-dark quencher (FQ)-labeled part S. All the described automata were assembled following this protocol.

### Automaton stability assay

Automata were assembled as described above (final concentration 1 μM) using LNA-modified L and S. Assembled samples were stored at −20 °C. Structure stability was performed by removing samples and storing them at room temperature over 29 days at different time points: 0, 1, 3, 8, 15, 22, 29 days. Sample integrity was analyzed by gel electrophoresis as described below.

### Gel electrophoresis

Samples were run on 4% agarose (Bio-Lab, Israel) in 1× TBE buffer (Biological industries, Israel) supplemented with 5 mM MgCl_2_ (Ambion), and stained with 0.3 µg/ml ethidium bromide (Sigma-Aldrich), in an ice-water bath (80–100 V, 2-4 h). Running buffer contained 1× TBE and 5 mM MgCl_2_. Trackit Ultra low range DNA ladder (Invitrogen), Low Molecular Weight DNA Ladder (New England Biolabs), ssRNA ladder and Low range ssRNA ladder standards (New England Biolabs) were used. In specific cases, samples were also run on Novex 10% Tris borate-EDTA (TBE) polyacrylamide gel (Invitrogen) and 10% mini-PROTEAN TBE-Urea Precast gels (Bio-Rad). Running buffer contained 1× TBE. Gels were run for 45 min at 200 V and then stained for 20 min in 0.5 µg/ml EtBr. Gels were imaged on a Bio-Rad Gel-Doc Imager and analyzed using ImageLab v5.2.1 and ImageJ v1.5 software. Gels containing fluorescent samples were scanned in a Nikon Eclipse Ti2 microscope using a Plan Fluor 4x PhL DL objective, in the Cy3 (Emission wavelength: 605.0, Excitation wavelength: 544.5) and EGFP (Emission wavelength: 525.0, Excitation wavelength: 470.0) channels.

### DNA origami folding

Previously-described DNA rectangles^8^ were re-designed using caDNAno^33^ to include 6 edge staples comprising the automaton S strand on its 3’ to enable rectangle-based assembly. M13mp18 ssDNA (7,249 base long) genome was ordered from New England BioLabs (N4040S) and used as scaffold strand. DNA staples were ordered from IDT. All DNA sequences are listed in Supplementary note 3. Folding was carried out at a 1:10 scaffold:staple ratio (scaffold concentration 10 nM) in 1× TAE buffer (Invitrogen), 12.5 mM MgCl_2_ (Ambion) in a BioRad C1000 Touch Thermal Cycler, using the following annealing ramp: 5 min at 85 °C, 85 °C to 60 °C at −1 °C/15min, 60 °C to 25°C at −1° C/3hr. Rectangles were purified by centrifugal gel filtration. Finally, automaton oligos (except for S) were added at a concentration of 500 nM.

### Atomic force microscopy

Samples were imaged in a JPK/Bruker NanoWizard Ultra AFM III instrument. The scans were performed at room temperature in FastScan-AC mode using ultra-short cantilevers with a force constant of 0.3 N/m (79427F7L1218, Nano World). For imaging, 20 μL (concentration of 1-2 nM) of sample was deposited on freshly cleaved mica (Grade V1, 10 mm, #50, Ted Pella, Inc.) for 5 min, followed by a gentle wash with 1× TAE buffer with 12.5 mM MgCl_2_. Finally, 1 mL of washing buffer was added for scanning in solution and not in dry conditions. Images were analyzed using JPK Data Processing v6.1.96.

### High-performance liquid chromatography (HPLC)

Samples were analyzed on an Agilent 1260 infinity II HPLC instrument, using a ProSEC 300S Size Exclusion 300×7.5 mm column (Agilent). The solvent was 1× Dulbecco’s Phosphate Buffered Saline (PBS, Biological Industries). Flow rate was set to 1 mL/min and samples were injected in 1× PBS. Detection was measured at 260 nm, at 30°c. Fractions from HPLC were collected according to their relevant retention time as set by standard oligonucleotides ordered (IDT) analyzed by HPLC under identical conditions. HPLC fractions were concentrated using Ultra 0.5mL centrifugal filters Ultracel® 10K (Merck Millipore LTD). Concentration and purity of fractions were measured using NanoDrop (Thermo Scientific) and gel electrophoresis.

### Fluorescence measurements

Fluorescence measurements were performed in a CFX96 Touch Real-Time PCR detection system (Bio-Rad), measuring fluorescence every 1 min at 27 and 37 °C.

### Cell culture

*Spodoptera frugiperda* (Sf9) cells (CRL-1711, ATCC) were maintained and propagated at 27°C in serum-free insect cell culture medium (Gibco™ SynerGy™ Sf-900™ II SFM, Cat # 10-902-161). Sf9 cells were grown either as monolayers in 6 well plates (Corning, Cat # 3516) or in shaker flasks (Corning, Cat # 430183) agitated at 140 rpm in a humidified environment.

### Virus system and cell infection

Bac-to-Bac® AcMNPV expression system was used to construct a donor plasmid (pFastBac™ Dual) that encodes for GFP under the very late polyhedrin promoter. Competent *E. coli* DH10BAC cells, containing bacmid (AcMNPV shuttle vector plasmid) and a helper plasmid, were used to generate recombinant bacmids according to the manufacturer’s protocol (Invitrogen). Insertion of the gene into the bacmid was verified by PCR. Sf9 cells were transfected with recombinant bacmid DNA using Escort™ IV Transfection Reagent (Sigma, Cat # L3287) in 6 well plates. The cells were incubated for 5 h at 27°C, rinsed, and incubated for another 72 h. Media were harvested, centrifuged, and the virus containing supernatant was used for 2-3 successive infections resulting in amplification of the virus titer. For cell infection, *Sf9* cells (3×10^6^ cells/ml) were infected with the recombinant viruses at various MOI (multiplicity of infection) values ranging from 0.1 to 10. Three days post infection cells were harvested by centrifugation at 500 g for 5 min.

### Delivery formulations and experiments

Automaton and enzyme payload were packaged in liposomes using a sterile liposome kit (L-4395-1VL, Sigma-Aldrich) according to the manufacturer’s instructions. Dried lipid powder was hydrated with serum-free, antibiotic free insect cell culture medium containing designated payload (100 pmol L, 100 pmol S, 1 pmol R, 17 pmol Bsu, 6.4 pmol T7 polymerase) in 1 mL PBS by vortexing for 30 sec. Liposome size distribution was measured using dynamic light scattering on a Malvern Nano-ZS ZEN3600 instrument. Alternatively, automaton and enzyme payload were delivered using Mirus (*Trans*IT-X2® Dynamic Delivery System for *CRISPR/Cas9 Ribonucleoprotein (RNP) + DNA Oligo (ssODN) Delivery*) according to the manufacturer’s instructions (only replacing Cas9 protein in the original protocol with Bsu and T7 polymerases).

### GP64 target preparation

The AcMNPV GP64 gene, cloned into pJ208-ampR vector under the T7 promoter, was ordered from Atum. Competent cells (HIT Competent Cells DH5α, RBS) were transformed according to the manufacturer’s instructions. Transformed cells were subsequently plated onto LB-agar plates supplemented with 100 μg/ml ampicillin (Hy-Labs, Israel). Single colonies were grown overnight at 37 °C in 3 ml of LB (Hy-Labs) supplemented with 100 μg/ml ampicillin (Sigma). Plasmid DNA was purified using QIAprep spin miniprep kit (Qiagen) according to the manufacturer’s instructions. The concentration and purity of the fractions was determined using NanoDrop (Thermo Scientific) and gel electrophoresis. GP64 RNA was synthesized from the prepared plasmid using MEGAscript T7 in vitro transcription kit (Thermo scientific) according to the manufacturer’s instructions, using 12 µg plasmid DNA as template. DNAse treatment with Turbo-DNAse (Thermo scientific) was used to clean the RNA. Transcribed RNA was purified with the MEGAclear kit (Thermo scientific) according to the manufacturer’s instructions, and re-suspended in ultrapure, nuclease-free water. Agarose gel electrophoresis and NanoDrop analysis were used to validate the integrity, quality, and concentration of the transcripts.

### GP64 quantification from cells

Total RNA was extracted from 2.3×10^6^ Sf9 cells in each sample using NucleoSpin® RNA kit (Macherey-Nagel), according to the manufacturer’s instructions. The concentration, purity, and integrity of total RNA was determined using NanoDrop and gel electrophoresis. cDNA synthesis was performed using iScript™ cDNA synthesis (Bio-Rad) according to the manufacturer’s instructions. Equal amounts of total RNA (1000 ng/20 µL) were reverse-transcribed in all samples. Reactions were incubated in a CFX96 Touch Real-Time PCR detection system (Bio-Rad) by the following program: 25 °C for 5 min, 46 °C for 20 min, 95 °C for 60 s. GP64 expression was determined by qPCR, with primers designed using NCBI primer-BLAST^87^. qPCR analysis was performed using the iTaq Universal SYBR Green Supermix (Bio-Rad) according to the manufacturer’s instructions, in a CFX96 system by the following program: 95 °C for 3 min, 39 cycles of 95 °C for 10 s and 55 °C for 30 s. Melt curves were generated for each sample by heating PCR amplicons from 65 °C to 95 °C with a gradual increase of 0.5 °C/0.5 s.

### Flow cytometry

Flow cytometry was performed on a Becton-Dickinson Accuri C6 Plus cytometer equipped with 488 nm solid-state laser and a 640 nm diode laser. Data was analyzed using Kaluza Analysis 2.1 software using a C6 import module. For intracellular flow cytometry (IFC), cells were fixed at 2% formaldehyde for 10 min on ice, permeabilized using methanol at −20 °C for 10 min on ice, and washed with cold IFC buffer (0.1 w/v nuclease-free bovine serum albumin, 0.01% w/v sodium azide in PBS, pH 7.6). Anti-phospho-p38 MAPK (Thr180/Tyr182), anti-phospho-p42/44 MAPK (ERK1/2) (Thr202/Tyr204), and anti-Akt (pan), having broad cross-reactivity ranging from human to *Drosophila*, and including dictyoptera^88^, were purchased from Cell Signaling (cat # 14594, 13148, 5186) and used at 1:50 dilution in 30 min incubation. For HHR measurements, cells were treated with DNase I for 15 min in PBS at room temperature, washed thoroughly with IFC buffer and incubated with a specific probe (/5BioTinTEG/CTGGAGTTTCGTCGCATTTCA/3Cy5Sp/). All cells were washed with IFC buffer prior to acquisition. For quantitative flow cytometry, calibration curves were produced immediately prior to each experiment using a Quantum-MESF calibration kit (Bangs Labs, USA) of the appropriate fluorochrome according to the manufacturer’s instructions.

### T7 RNA polymerase labeling

Labeling of T7 RNA polymerase (Takara bio) was performed using Alexa Fluor™ 647 Protein Labeling kit (Thermo scientific) according to the manufacturer’s instructions.

### Animal studies

Adult *B. discoidalis* of both sexes were purchased from two independent farms, Yaldei Ha-teva or Beit Haiut (Israel), and housed at room temperature and humidity in large plastic containers. Insects were supplied with dry pet food, fresh fruit, and water *ad libitum*. Egg cartons were used as light shelters. Maximum number of insects per cage was limited to 12. For injection, insects were anesthetized one by one, by placement at −20 °C for 7-8 min (depending on animal behavior) in a fresh glass beaker. Once an insect has been anesthetized, a 15 *μ*L containing 4×10^5^ Sf9 cells either non-treated, infected, or infected and delivered with automaton payload were injected into the hemocoel using a Hamilton syringe. Infection was done at MOI of 0.1. Injection was carried out through the soft membrane between the 2nd-3rd abdominal sternites, close to the lateral body margin. Following injection, insects were left to recuperate. For hemolymph extraction, insects were anesthetized as described above. Hemolymph was extracted by puncturing the arthrodial membrane at the base of either metathoracic leg with an ice-cold needle dipped in anticoagulation buffer (30 mM citric acid, 30 mM sodium citrate, 1 mM EDTA). 25-50 µL of hemolymph were pipetted into a similar volume of ice-cold buffer and immediately diluted into 300 µL of ice-cold buffer in a fresh tube on ice.

### Statistical analysis

Statistical tests were performed with GraphPad Prism (version 8.3.0). All experiments were repeated a minimum of 3 independent times. *P*-values in each experiment were determined by unpaired two-tailed Student’s *t* test with 95% confidence interval. In all flow cytometric experiments two biological replicates were acquired per group, and a minimum of 10,000 total events was collected. Mean and %CV of gated events were used to derive SD for determination of statistical significance.

## Acknowledgements

The authors wish to thank the entire team at Augmanity, particularly to Dr. R. Spokoini-Stern, for valuable discussions and technical assistance.

## Author contributions

All authors performed experiments, analyzed experiments, and wrote the manuscript. IB supervised research.

## Supplementary note 1

### Prototype design and simulation

The design of the target was based on incorporation of the cleavage site of the chosen HHR (from the self-limiting automata (3WJ) design) into a random sequence in the loop site of a loop and stem structure that after HHR-mediated cleavage form a stable stem with flanking arms **(Fig. S1)**.

The automata consist of two DNA oligonucleotides that have an eight base complementation to each other and eight base complementation to regions in the cleaved target molecule. The L strand consists of a T7 promoter sequence, a complementary sequence of the HHR gene and a 3’ phosphorylation modification to prevent elongation from the L strand.

The automaton can be assembled only in the presence of R (cleaved target), which consists of two subunits that form a stable stem with flanking arms which represent a cleaved target molecule **(Fig. S2).**

Moreover, we tested the stability and shelf life time of the LNA-modified automaton comprising 7 LNA bases in L and S strands at room temperature over 29 days. The LNA-modified version remained intact. In addition, the assembly continued to occur as indicated by the increased intensity of the bands **(Fig. S3)**.

To protect the assembled automaton from RNase H, several variants were tested in which parts L and S were modified with either locked nucleic acids (LNA) or 2-O-methoxyethyl RNA (2-MOE) at various lengths along the regions complementary to part R (3, 5, and 7 nt). Different combinations of modifications were found effective in protecting the assembled automaton from RNase H **(Fig. S4)**

**Figure S1.**
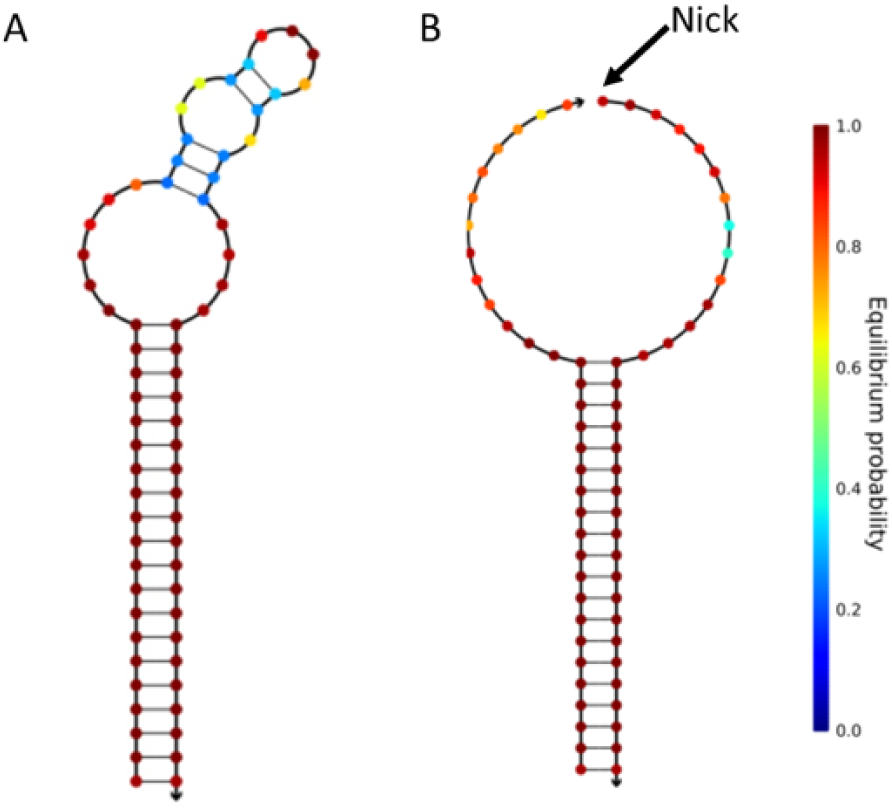
NUPACK simulation of the target structure. Before (**A**) and after (**B**) HHR cleavage at 37 °C.

**Figure S2.**
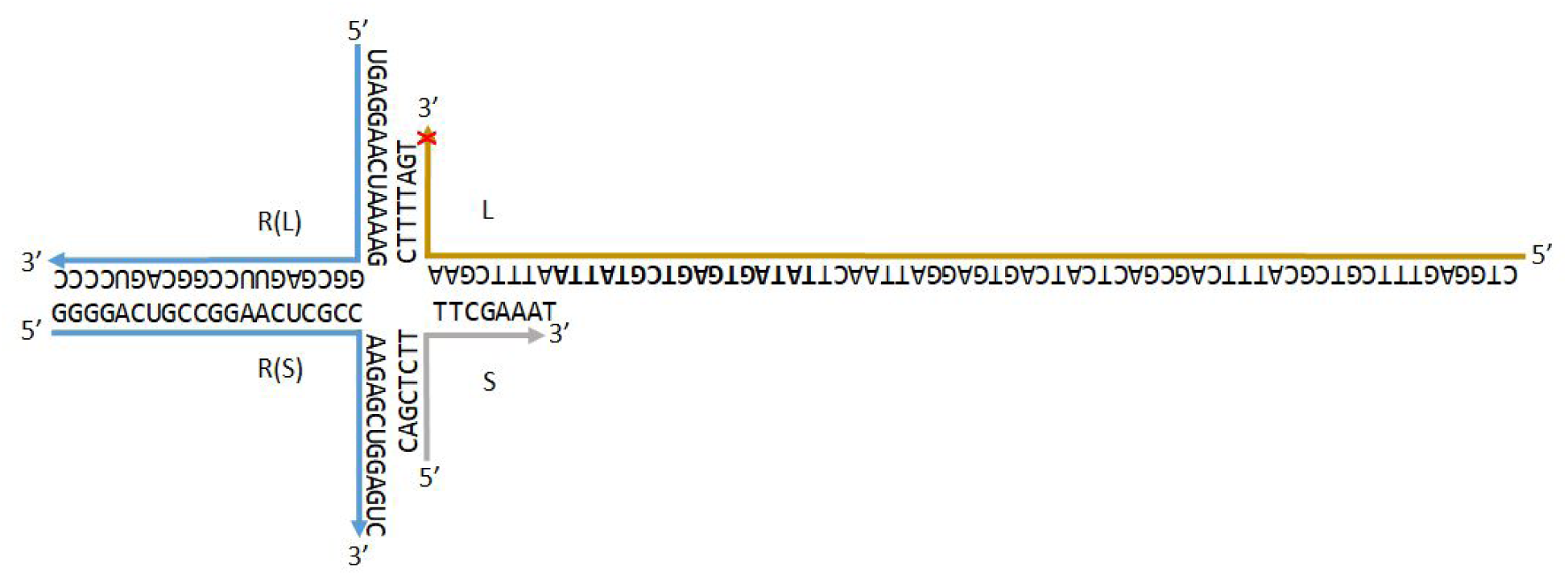
Schematic representation of the prototype automaton. RNA stem with two flanking arms with eight bases complementation to two DNA strands (L, S). L and S strands have eight bases complementing each other. The L strand consists of a T7 promoter sequence (bold), a complementary sequence of the HHR and a 3’ phosphorylation modification (red star).

**Figure S3.**
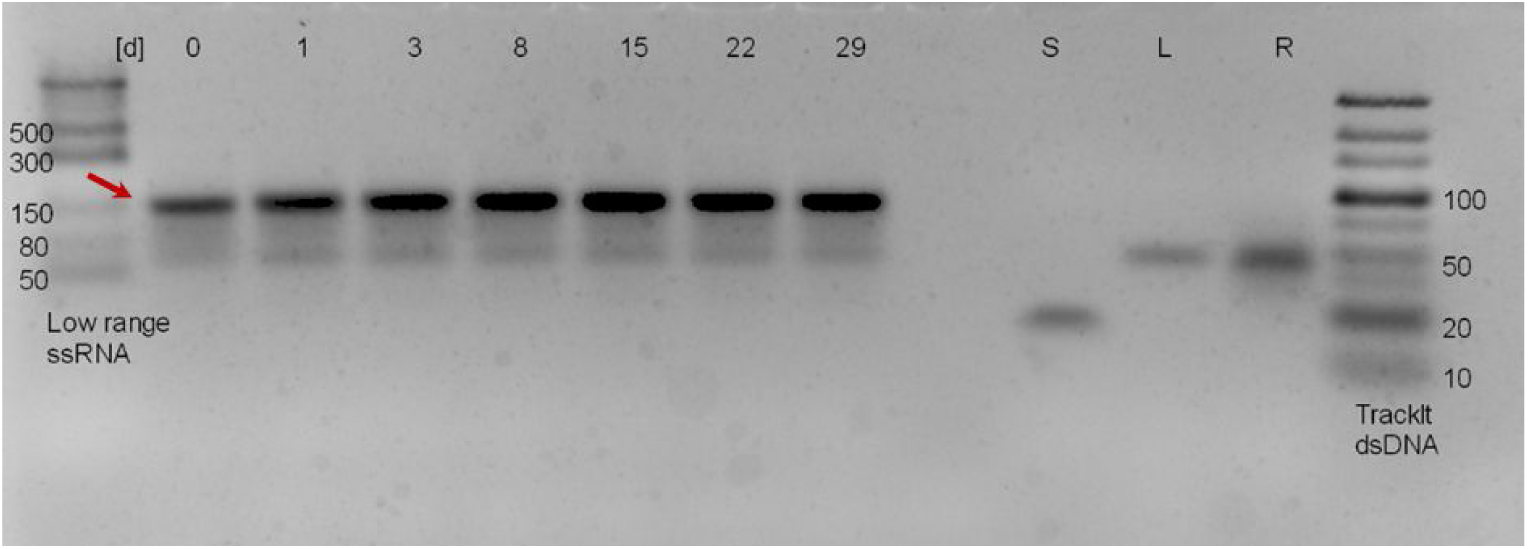
Long-term stability. Long-term stability of the LNA modified automaton (both L and S arms comprising 7 LNA bases) at room temperature over 29 days. The arrow indicates the band of fully assembled automaton into a 4-way junction.

**Figure S4.**
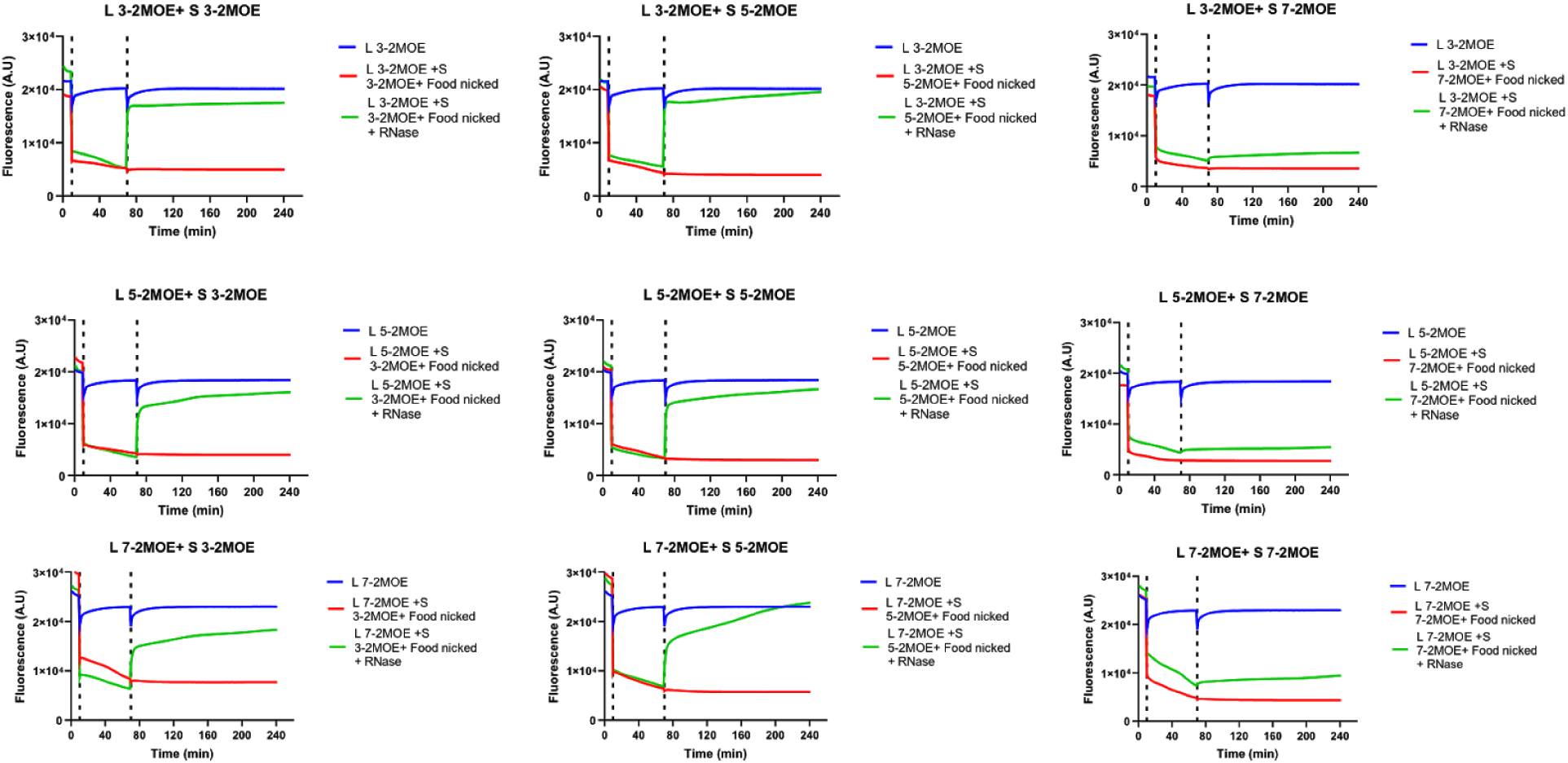
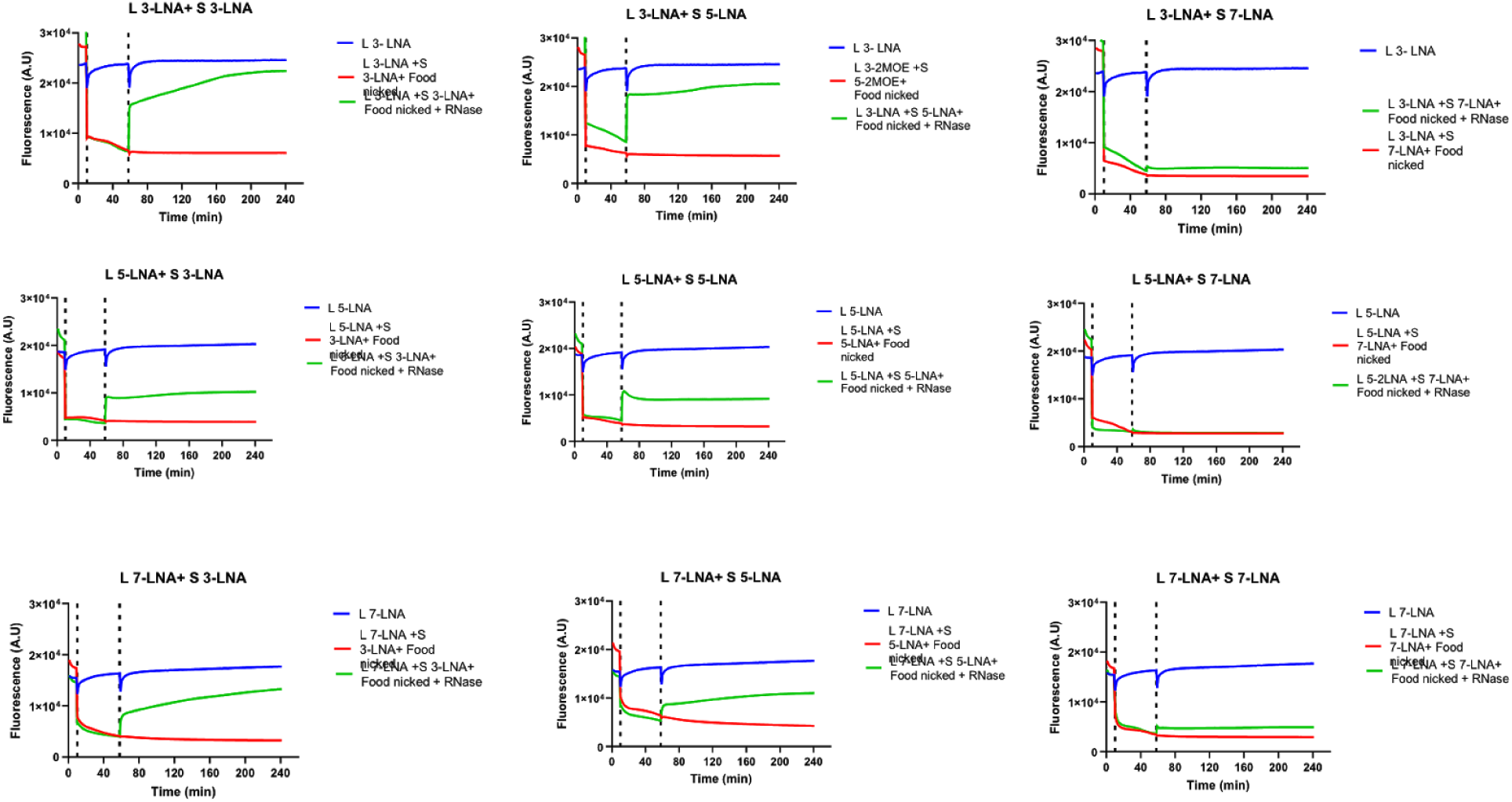
Modification in the automaton that protects from RNase H. Following automaton assembly and stability using modified FAM labeled L arm and modified dark FQ S arm in a RT-PCR machine. L and S were modified with either locked nucleic acids (LNA) or 2-O-methoxyethyl RNA (2-MOE) at various lengths along the regions complementary to part R (3, 5, and 7 nt). Initially the fluorescence of 1 µM of FAM labeled modified L was followed every 1 min using the FAM channel for 10 min, then the rest of the oligos (S, R(L), R(S)) were added to the reaction accordingly to final concentration of 1 µM. The fluorescence was measured every 1 min for an additional 60 min using the FAM channel. Then 50 units of RNase H (New England Biolabs) were added to the reaction and the fluorescence was measured every 1 min for an additional 120 min using the FAM channel.

## Supplementary note 2

### Automaton (prototype) sequence list

**Table S1.**
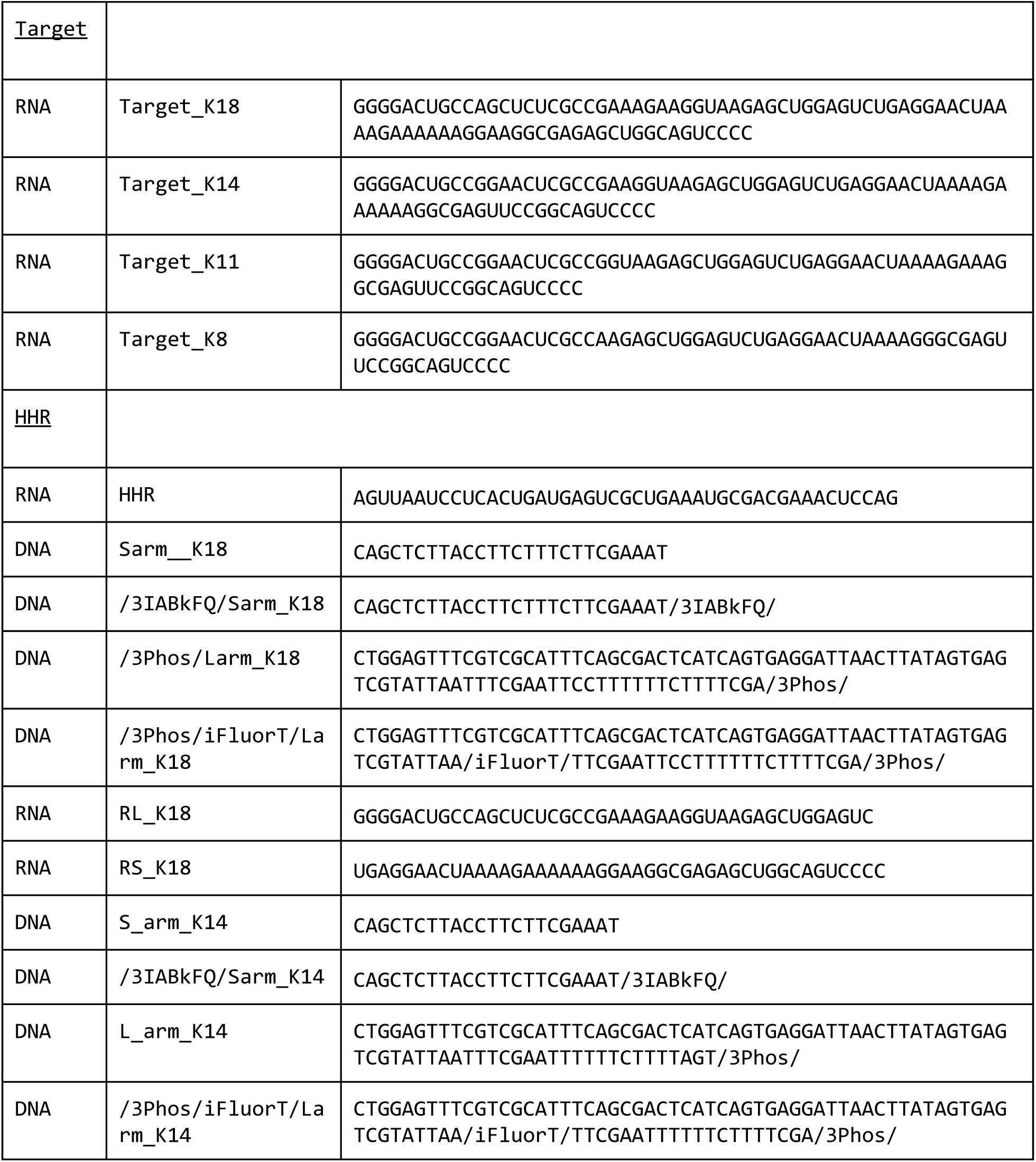
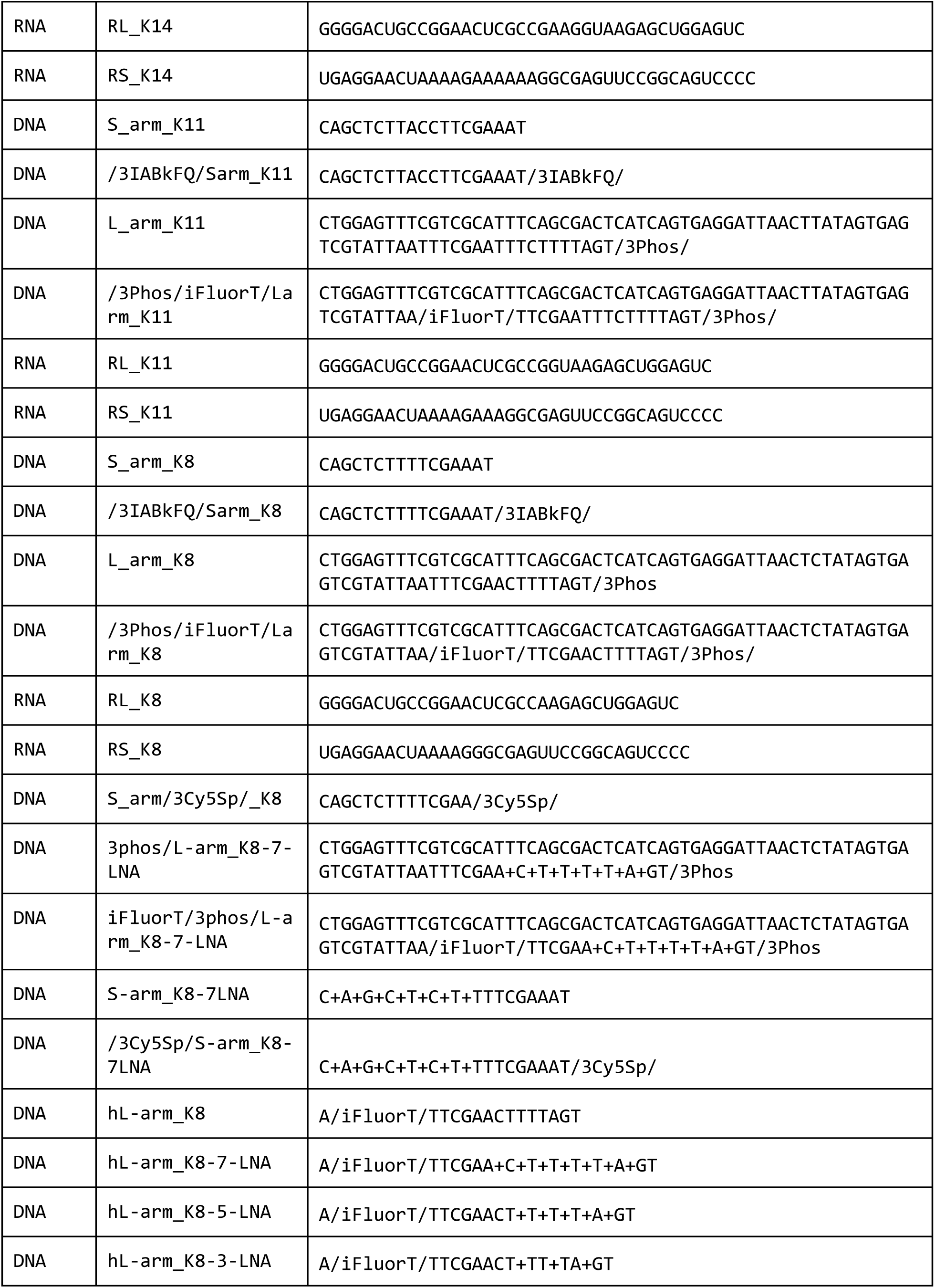
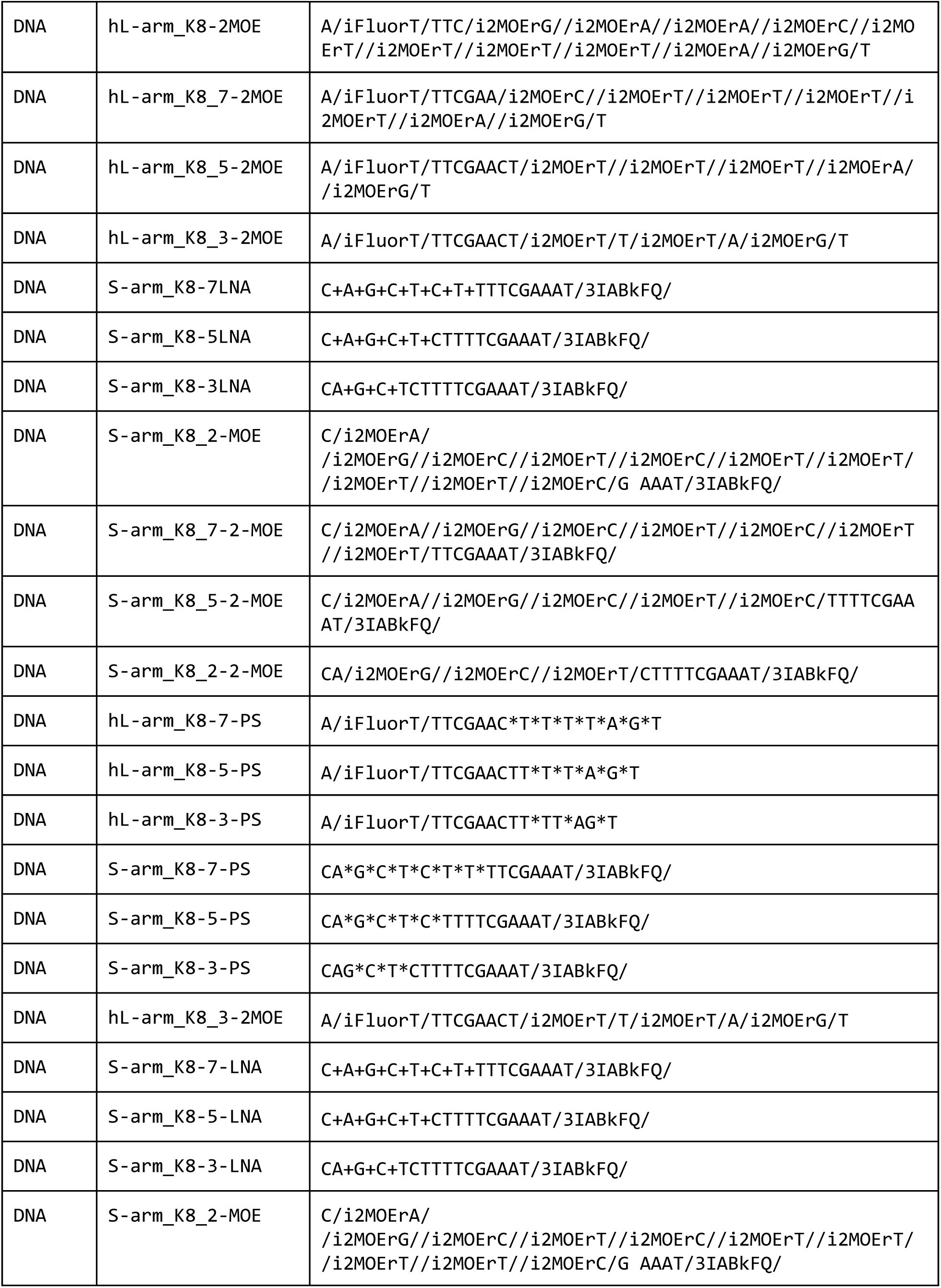
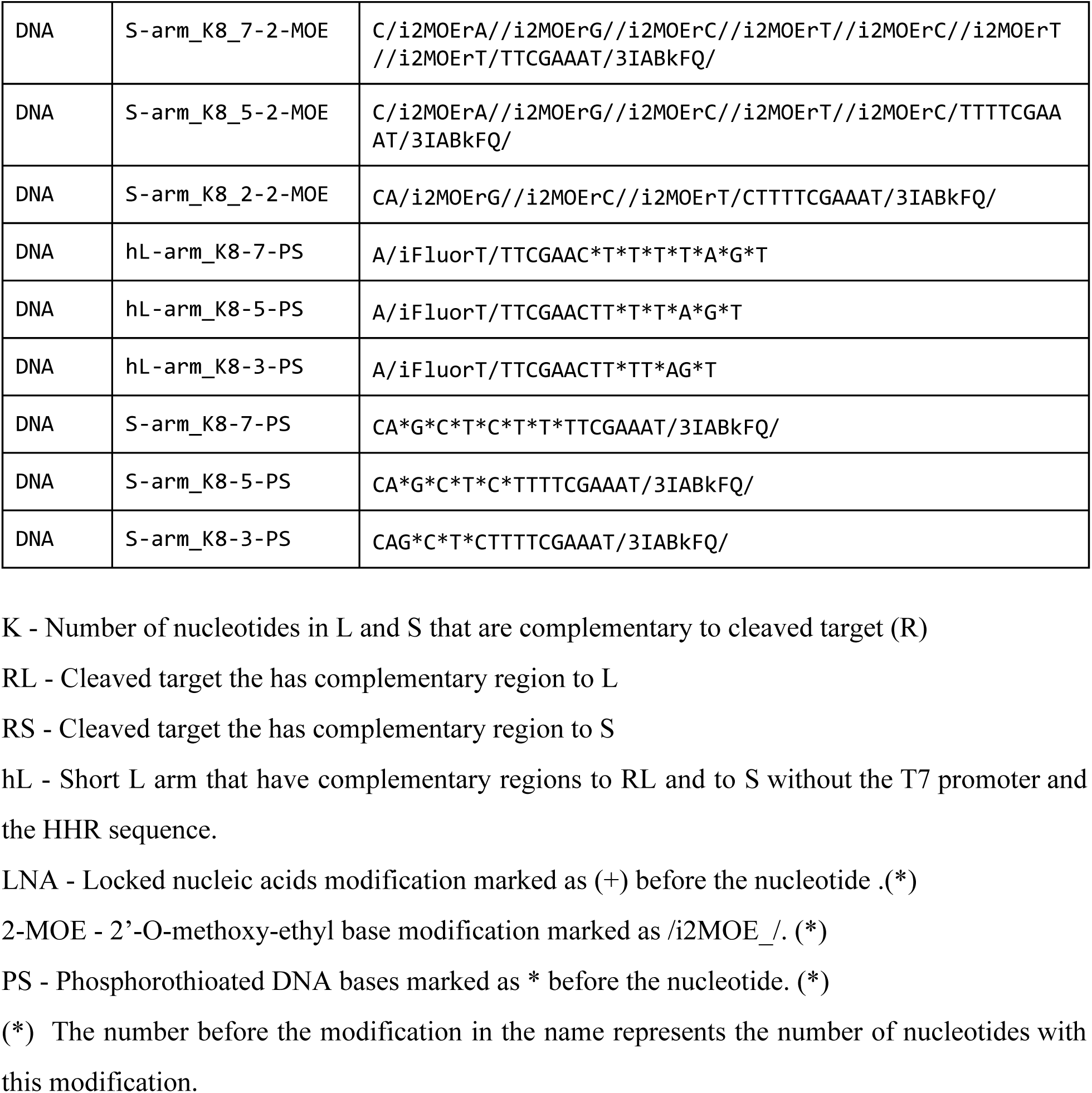
Automaton (prototype) sequence list.

## Supplementary note 3

### DNA origami rectangle design

Imaging molecular automaton itself was challenging since it is nearly AFM (atomic force microscopy) resolution limit (∼2nm) and built from short and partially single stranded regions [L-R(L) region is 2.75 nm, R(L)-R(S) region is 6.8 nm, and S-L is partially single stranded and 27 nm]. To facilitate the imaging of the molecular automaton, we attached it to a reference shape, which is bigger and more stable. For this purpose we adjusted Rothmund^8^ standard DNA rectangle design **(Fig. S5)** and elongated six edges staples at 3’ with – TTTTTTTT CAGCTCTTTTCGAAAT-3’ (8T followed by S strand) **(Fig. S6)**. This way we managed to scan and visualize fully assembled automata **(Fig. 1C, Fig. S7**). The folding of the DNA rectangles with the automaton and AFM scans described in methods and materials section.

**Figure S5.**
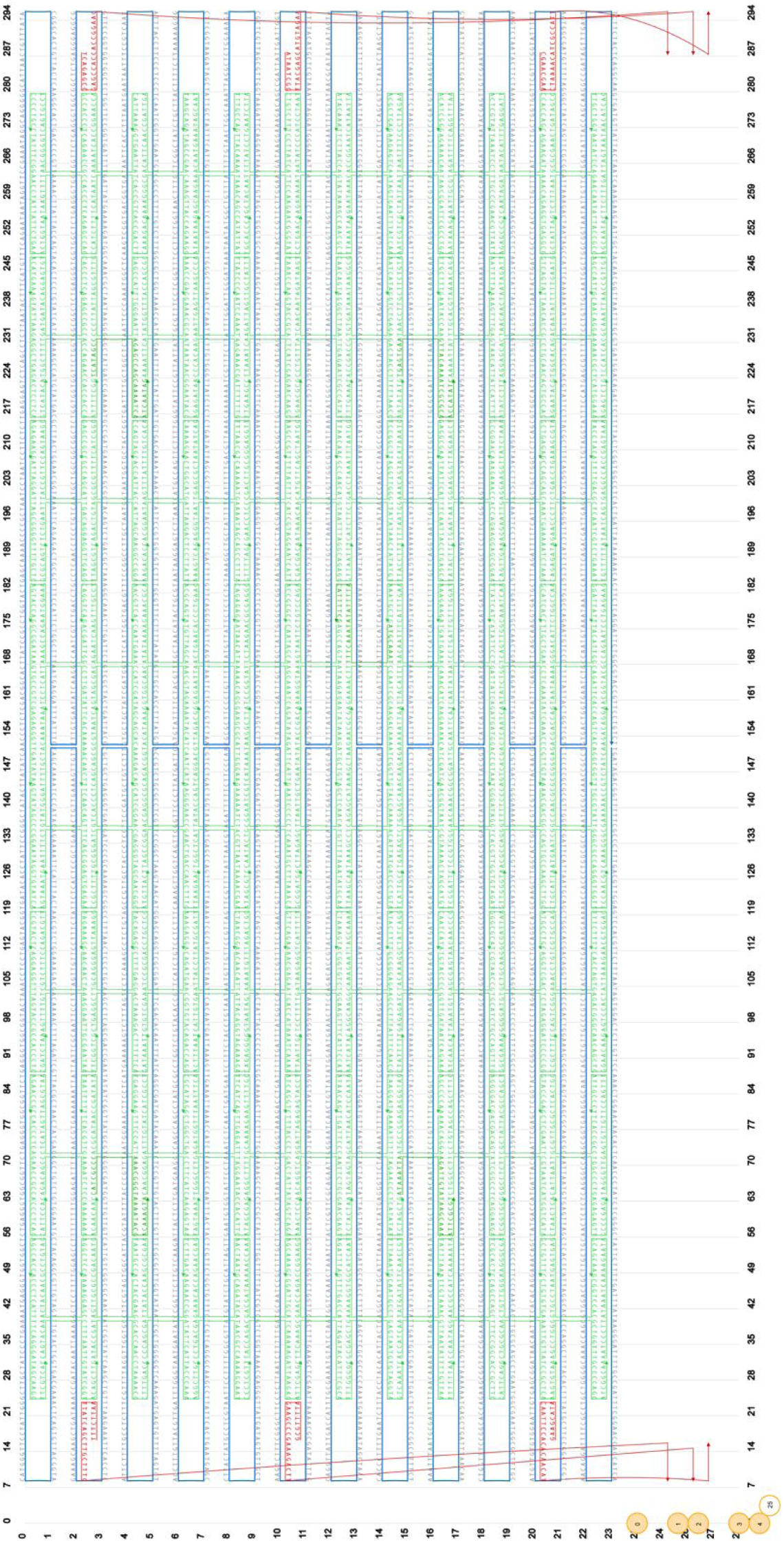
Modified DNA origami rectangles design using caDNAno. Red oligos represent the extended six edges staples. 8T and S sequences have been added manually to 3’ and do not appear in the caDNAno scheme.

**Figure S6.**
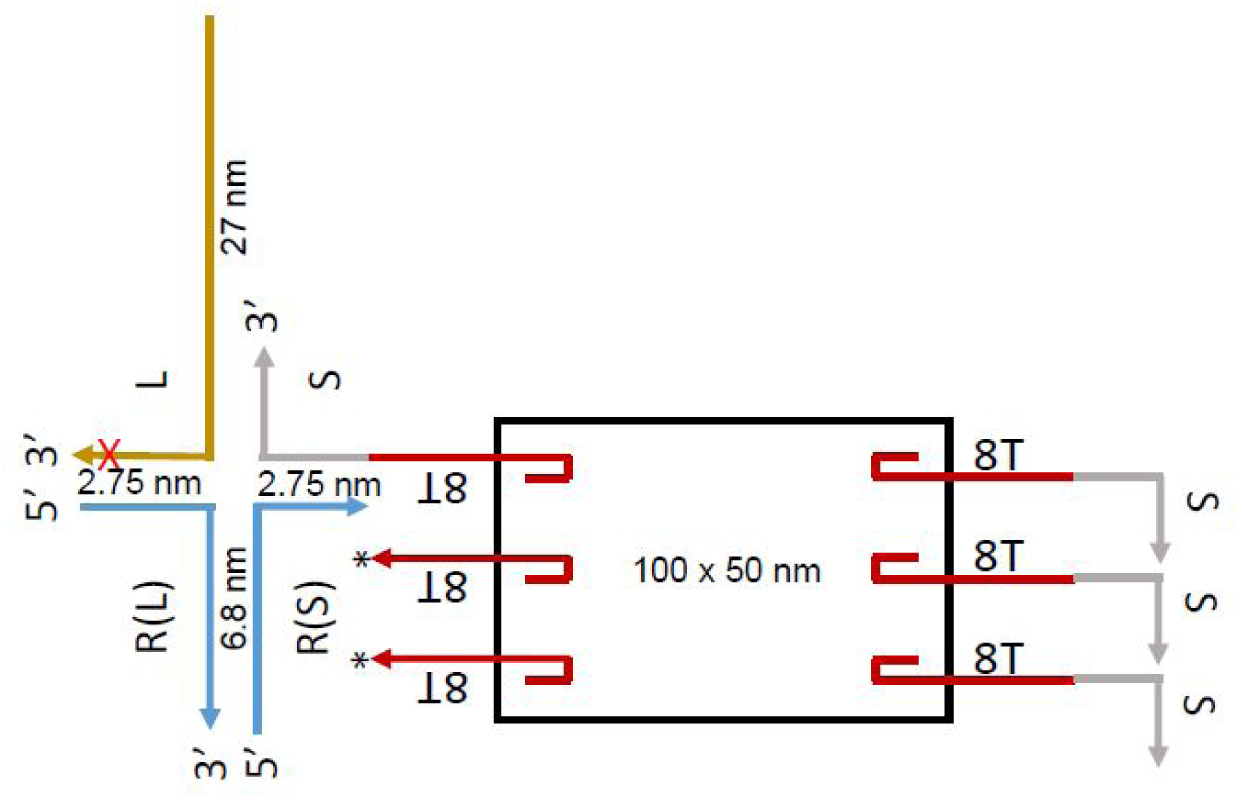
Schematic representation of a molecular automaton. attached to DNA origami rectangles via six edges staples containing S on 3’. The * at the left two staples represents the S strand attached to the staples. Approximated sizes are indicated.

**Figure S7.**
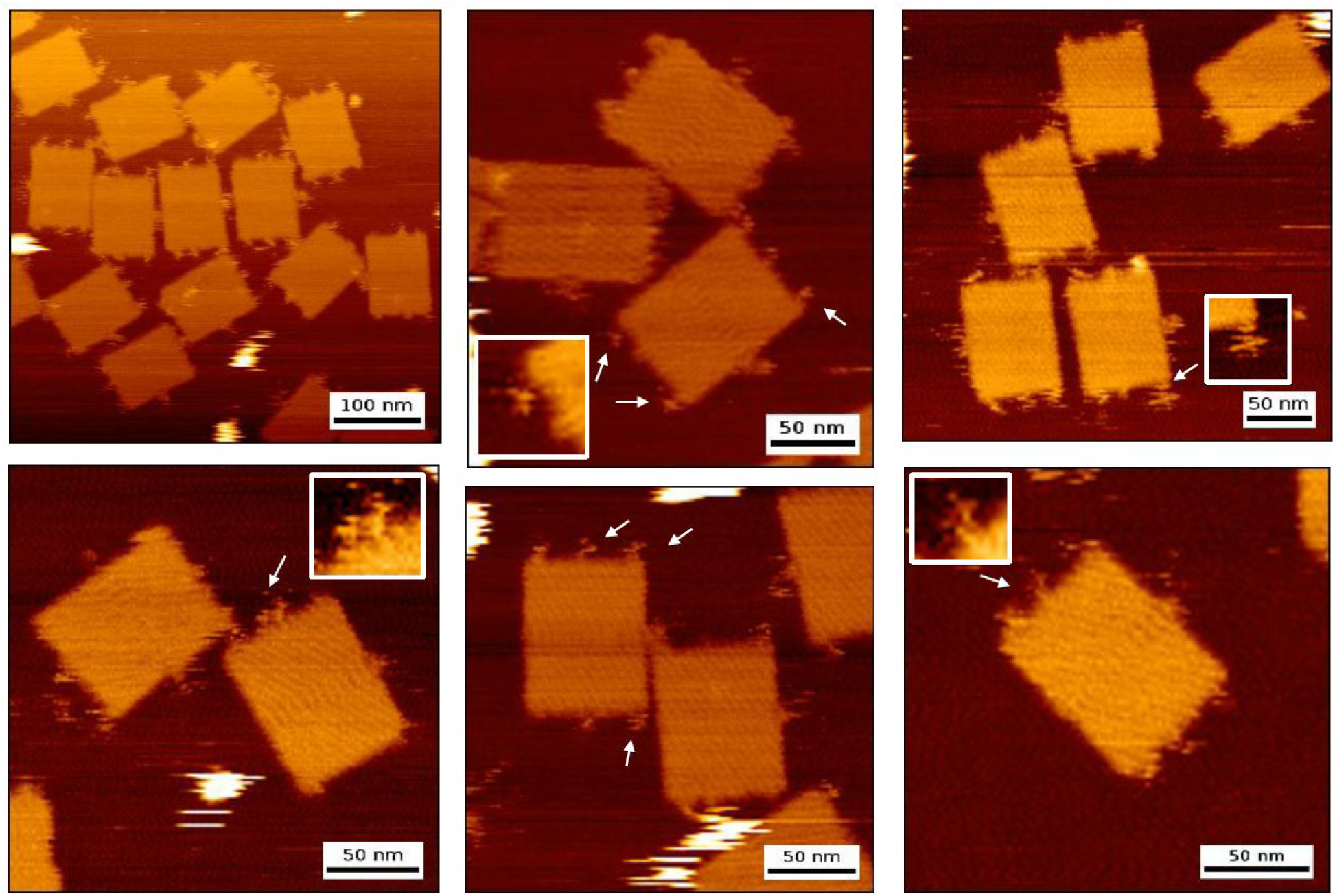
AFM scans. Images of the prototype automaton attached to DNA rectangles.

**<u>M13mp18 Scaffold</u>**

**<u>>M13mp18 [length=7249] [version=09-MAY-2008] [topology=circular] Cloning vector M13mp18, complete sequence.</u>**

AATGCTACTACTATTAGTAGAATTGATGCCACCTTTTCAGCTCGCGCCCCAAATGAAAATATAGCTAAACAGGTTATTGACCATT TGCGAAATGTATCTAATGGTCAAACTAAATCTACTCGTTCGCAGAATTGGGAATCAACTGTTATATGGAATGAAACTTCCAGACA CCGTACTTTAGTTGCATATTTAAAACATGTTGAGCTACAGCATTATATTCAGCAATTAAGCTCTAAGCCATCCGCAAAAATGACC TCTTATCAAAAGGAGCAATTAAAGGTACTCTCTAATCCTGACCTGTTGGAGTTTGCTTCCGGTCTGGTTCGCTTTGAAGCTCGAA TTAAAACGCGATATTTGAAGTCTTTCGGGCTTCCTCTTAATCTTTTTGATGCAATCCGCTTTGCTTCTGACTATAATAGTCAGGG TAAAGACCTGATTTTTGATTTATGGTCATTCTCGTTTTCTGAACTGTTTAAAGCATTTGAGGGGGATTCAATGAATATTTATGAC GATTCCGCAGTATTGGACGCTATCCAGTCTAAACATTTTACTATTACCCCCTCTGGCAAAACTTCTTTTGCAAAAGCCTCTCGCT ATTTTGGTTTTTATCGTCGTCTGGTAAACGAGGGTTATGATAGTGTTGCTCTTACTATGCCTCGTAATTCCTTTTGGCGTTATGT ATCTGCATTAGTTGAATGTGGTATTCCTAAATCTCAACTGATGAATCTTTCTACCTGTAATAATGTTGTTCCGTTAGTTCGTTTT ATTAACGTAGATTTTTCTTCCCAACGTCCTGACTGGTATAATGAGCCAGTTCTTAAAATCGCATAAGGTAATTCACAATGATTAA AGTTGAAATTAAACCATCTCAAGCCCAATTTACTACTCGTTCTGGTGTTTCTCGTCAGGGCAAGCCTTATTCACTGAATGAGCAG CTTTGTTACGTTGATTTGGGTAATGAATATCCGGTTCTTGTCAAGATTACTCTTGATGAAGGTCAGCCAGCCTATGCGCCTGGTC TGTACACCGTTCATCTGTCCTCTTTCAAAGTTGGTCAGTTCGGTTCCCTTATGATTGACCGTCTGCGCCTCGTTCCGGCTAAGTA ACATGGAGCAGGTCGCGGATTTCGACACAATTTATCAGGCGATGATACAAATCTCCGTTGTACTTTGTTTCGCGCTTGGTATAAT CGCTGGGGGTCAAAGATGAGTGTTTTAGTGTATTCTTTTGCCTCTTTCGTTTTAGGTTGGTGCCTTCGTAGTGGCATTACGTATT TTACCCGTTTAATGGAAACTTCCTCATGAAAAAGTCTTTAGTCCTCAAAGCCTCTGTAGCCGTTGCTACCCTCGTTCCGATGCTG TCTTTCGCTGCTGAGGGTGACGATCCCGCAAAAGCGGCCTTTAACTCCCTGCAAGCCTCAGCGACCGAATATATCGGTTATGCGT GGGCGATGGTTGTTGTCATTGTCGGCGCAACTATCGGTATCAAGCTGTTTAAGAAATTCACCTCGAAAGCAAGCTGATAAACCGA TACAATTAAAGGCTCCTTTTGGAGCCTTTTTTTTGGAGATTTTCAACGTGAAAAAATTATTATTCGCAATTCCTTTAGTTGTTCC TTTCTATTCTCACTCCGCTGAAACTGTTGAAAGTTGTTTAGCAAAATCCCATACAGAAAATTCATTTACTAACGTCTGGAAAGAC GACAAAACTTTAGATCGTTACGCTAACTATGAGGGCTGTCTGTGGAATGCTACAGGCGTTGTAGTTTGTACTGGTGACGAAACTC AGTGTTACGGTACATGGGTTCCTATTGGGCTTGCTATCCCTGAAAATGAGGGTGGTGGCTCTGAGGGTGGCGGTTCTGAGGGTGG CGGTTCTGAGGGTGGCGGTACTAAACCTCCTGAGTACGGTGATACACCTATTCCGGGCTATACTTATATCAACCCTCTCGACGGC ACTTATCCGCCTGGTACTGAGCAAAACCCCGCTAATCCTAATCCTTCTCTTGAGGAGTCTCAGCCTCTTAATACTTTCATGTTTC AGAATAATAGGTTCCGAAATAGGCAGGGGGCATTAACTGTTTATACGGGCACTGTTACTCAAGGCACTGACCCCGTTAAAACTTA TTACCAGTACACTCCTGTATCATCAAAAGCCATGTATGACGCTTACTGGAACGGTAAATTCAGAGACTGCGCTTTCCATTCTGGC TTTAATGAGGATTTATTTGTTTGTGAATATCAAGGCCAATCGTCTGACCTGCCTCAACCTCCTGTCAATGCTGGCGGCGGCTCTG GTGGTGGTTCTGGTGGCGGCTCTGAGGGTGGTGGCTCTGAGGGTGGCGGTTCTGAGGGTGGCGGCTCTGAGGGAGGCGGTTCCGG TGGTGGCTCTGGTTCCGGTGATTTTGATTATGAAAAGATGGCAAACGCTAATAAGGGGGCTATGACCGAAAATGCCGATGAAAAC GCGCTACAGTCTGACGCTAAAGGCAAACTTGATTCTGTCGCTACTGATTACGGTGCTGCTATCGATGGTTTCATTGGTGACGTTT CCGGCCTTGCTAATGGTAATGGTGCTACTGGTGATTTTGCTGGCTCTAATTCCCAAATGGCTCAAGTCGGTGACGGTGATAATTC ACCTTTAATGAATAATTTCCGTCAATATTTACCTTCCCTCCCTCAATCGGTTGAATGTCGCCCTTTTGTCTTTGGCGCTGGTAAA CCATATGAATTTTCTATTGATTGTGACAAAATAAACTTATTCCGTGGTGTCTTTGCGTTTCTTTTATATGTTGCCACCTTTATGT ATGTATTTTCTACGTTTGCTAACATACTGCGTAATAAGGAGTCTTAATCATGCCAGTTCTTTTGGGTATTCCGTTATTATTGCGT TTCCTCGGTTTCCTTCTGGTAACTTTGTTCGGCTATCTGCTTACTTTTCTTAAAAAGGGCTTCGGTAAGATAGCTATTGCTATTT CATTGTTTCTTGCTCTTATTATTGGGCTTAACTCAATTCTTGTGGGTTATCTCTCTGATATTAGCGCTCAATTACCCTCTGACTT TGTTCAGGGTGTTCAGTTAATTCTCCCGTCTAATGCGCTTCCCTGTTTTTATGTTATTCTCTCTGTAAAGGCTGCTATTTTCATT TTTGACGTTAAACAAAAAATCGTTTCTTATTTGGATTGGGATAAATAATATGGCTGTTTATTTTGTAACTGGCAAATTAGGCTCT GGAAAGACGCTCGTTAGCGTTGGTAAGATTCAGGATAAAATTGTAGCTGGGTGCAAAATAGCAACTAATCTTGATTTAAGGCTTC AAAACCTCCCGCAAGTCGGGAGGTTCGCTAAAACGCCTCGCGTTCTTAGAATACCGGATAAGCCTTCTATATCTGATTTGCTTGC TATTGGGCGCGGTAATGATTCCTACGATGAAAATAAAAACGGCTTGCTTGTTCTCGATGAGTGCGGTACTTGGTTTAATACCCGT TCTTGGAATGATAAGGAAAGACAGCCGATTATTGATTGGTTTCTACATGCTCGTAAATTAGGATGGGATATTATTTTTCTTGTTC AGGACTTATCTATTGTTGATAAACAGGCGCGTTCTGCATTAGCTGAACATGTTGTTTATTGTCGTCGTCTGGACAGAATTACTTT ACCTTTTGTCGGTACTTTATATTCTCTTATTACTGGCTCGAAAATGCCTCTGCCTAAATTACATGTTGGCGTTGTTAAATATGGC GATTCTCAATTAAGCCCTACTGTTGAGCGTTGGCTTTATACTGGTAAGAATTTGTATAACGCATATGATACTAAACAGGCTTTTT CTAGTAATTATGATTCCGGTGTTTATTCTTATTTAACGCCTTATTTATCACACGGTCGGTATTTCAAACCATTAAATTTAGGTCA GAAGATGAAATTAACTAAAATATATTTGAAAAAGTTTTCTCGCGTTCTTTGTCTTGCGATTGGATTTGCATCAGCATTTACATAT AGTTATATAACCCAACCTAAGCCGGAGGTTAAAAAGGTAGTCTCTCAGACCTATGATTTTGATAAATTCACTATTGACTCTTCTC AGCGTCTTAATCTAAGCTATCGCTATGTTTTCAAGGATTCTAAGGGAAAATTAATTAATAGCGACGATTTACAGAAGCAAGGTTA TTCACTCACATATATTGATTTATGTACTGTTTCCATTAAAAAAGGTAATTCAAATGAAATTGTTAAATGTAATTAATTTTGTTTT CTTGATGTTTGTTTCATCATCTTCTTTTGCTCAGGTAATTGAAATGAATAATTCGCCTCTGCGCGATTTTGTAACTTGGTATTCA AAGCAATCAGGCGAATCCGTTATTGTTTCTCCCGATGTAAAAGGTACTGTTACTGTATATTCATCTGACGTTAAACCTGAAAATC TACGCAATTTCTTTATTTCTGTTTTACGTGCAAATAATTTTGATATGGTAGGTTCTAACCCTTCCATTATTCAGAAGTATAATCC AAACAATCAGGATTATATTGATGAATTGCCATCATCTGATAATCAGGAATATGATGATAATTCCGCTCCTTCTGGTGGTTTCTTT GTTCCGCAAAATGATAATGTTACTCAAACTTTTAAAATTAATAACGTTCGGGCAAAGGATTTAATACGAGTTGTCGAATTGTTTG TAAAGTCTAATACTTCTAAATCCTCAAATGTATTATCTATTGACGGCTCTAATCTATTAGTTGTTAGTGCTCCTAAAGATATTTT AGATAACCTTCCTCAATTCCTTTCAACTGTTGATTTGCCAACTGACCAGATATTGATTGAGGGTTTGATATTTGAGGTTCAGCAA GGTGATGCTTTAGATTTTTCATTTGCTGCTGGCTCTCAGCGTGGCACTGTTGCAGGCGGTGTTAATACTGACCGCCTCACCTCTG TTTTATCTTCTGCTGGTGGTTCGTTCGGTATTTTTAATGGCGATGTTTTAGGGCTATCAGTTCGCGCATTAAAGACTAATAGCCA TTCAAAAATATTGTCTGTGCCACGTATTCTTACGCTTTCAGGTCAGAAGGGTTCTATCTCTGTTGGCCAGAATGTCCCTTTTATT ACTGGTCGTGTGACTGGTGAATCTGCCAATGTAAATAATCCATTTCAGACGATTGAGCGTCAAAATGTAGGTATTTCCATGAGCG TTTTTCCTGTTGCAATGGCTGGCGGTAATATTGTTCTGGATATTACCAGCAAGGCCGATAGTTTGAGTTCTTCTACTCAGGCAAG TGATGTTATTACTAATCAAAGAAGTATTGCTACAACGGTTAATTTGCGTGATGGACAGACTCTTTTACTCGGTGGCCTCACTGAT TATAAAAACACTTCTCAGGATTCTGGCGTACCGTTCCTGTCTAAAATCCCTTTAATCGGCCTCCTGTTTAGCTCCCGCTCTGATT CTAACGAGGAAAGCACGTTATACGTGCTCGTCAAAGCAACCATAGTACGCGCCCTGTAGCGGCGCATTAAGCGCGGCGGGTGTGG TGGTTACGCGCAGCGTGACCGCTACACTTGCCAGCGCCCTAGCGCCCGCTCCTTTCGCTTTCTTCCCTTCCTTTCTCGCCACGTT CGCCGGCTTTCCCCGTCAAGCTCTAAATCGGGGGCTCCCTTTAGGGTTCCGATTTAGTGCTTTACGGCACCTCGACCCCAAAAAA CTTGATTTGGGTGATGGTTCACGTAGTGGGCCATCGCCCTGATAGACGGTTTTTCGCCCTTTGACGTTGGAGTCCACGTTCTTTA ATAGTGGACTCTTGTTCCAAACTGGAACAACACTCAACCCTATCTCGGGCTATTCTTTTGATTTATAAGGGATTTTGCCGATTTC GGAACCACCATCAAACAGGATTTTCGCCTGCTGGGGCAAACCAGCGTGGACCGCTTGCTGCAACTCTCTCAGGGCCAGGCGGTGA AGGGCAATCAGCTGTTGCCCGTCTCACTGGTGAAAAGAAAAACCACCCTGGCGCCCAATACGCAAACCGCCTCTCCCCGCGCGTT GGCCGATTCATTAATGCAGCTGGCACGACAGGTTTCCCGACTGGAAAGCGGGCAGTGAGCGCAACGCAATTAATGTGAGTTAGCT CACTCATTAGGCACCCCAGGCTTTACACTTTATGCTTCCGGCTCGTATGTTGTGTGGAATTGTGAGCGGATAACAATTTCACACA GGAAACAGCTATGACCATGATTACGAATTCGAGCTCGGTACCCGGGGATCCTCTAGAGTCGACCTGCAGGCATGCAAGCTTGGCA CTGGCCGTCGTTTTACAACGTCGTGACTGGGAAAACCCTGGCGTTACCCAACTTAATCGCCTTGCAGCACATCCCCCTTTCGCCA GCTGGCGTAATAGCGAAGAGGCCCGCACCGATCGCCCTTCCCAACAGTTGCGCAGCCTGAATGGCGAATGGCGCTTTGCCTGGTT TCCGGCACCAGAAGCGGTGCCGGAAAGCTGGCTGGAGTGCGATCTTCCTGAGGCCGATACTGTCGTCGTCCCCTCAAACTGGCAG ATGCACGGTTACGATGCGCCCATCTACACCAACGTGACCTATCCCATTACGGTCAATCCGCCGTTTGTTCCCACGGAGAATCCGA CGGGTTGTTACTCGCTCACATTTAATGTTGATGAAAGCTGGCTACAGGAAGGCCAGACGCGAATTATTTTTGATGGCGTTCCTAT TGGTTAAAAAATGAGCTGATTTAACAAAAATTTAATGCGAATTTTAACAAAATATTAACGTTTACAATTTAAATATTTGCTTATA CAATCTTCCTGTTTTTGGGGCTTTTCTGATTATCAACCGGGGTACATATGATTGACATGCTAGTTTTACGATTACCGTTCATCGA TTCTCTTGTTTGCTCCAGACTCTCAGGCAATGACCTGATAGCCTTTGTAGATCTCTCAAAAATAGCTACCCTCTCCGGCATTAAT TTATCAGCTAGAACGGTTGAATATCATATTGATGGTGATTTGACTGTCTCCGGCCTTTCTCACCCTTTTGAATCTTTACCTACAC ATTACTCAGGCATTGCATTTAAAATATATGAGGGTTCTAAAAATTTTTATCCTTGCGTTGAAATAAAGGCTTCTCCCGCAAAAGT ATTACAGGGTCATAATGTTTTTGGTACAACCGATTTAGCTTTATGCTCTGAGGCTTTATTGCTTAATTTTGCTAATTCTTTGCCT TGCCTGTATGATTTATTGGATGTT

### Staple List

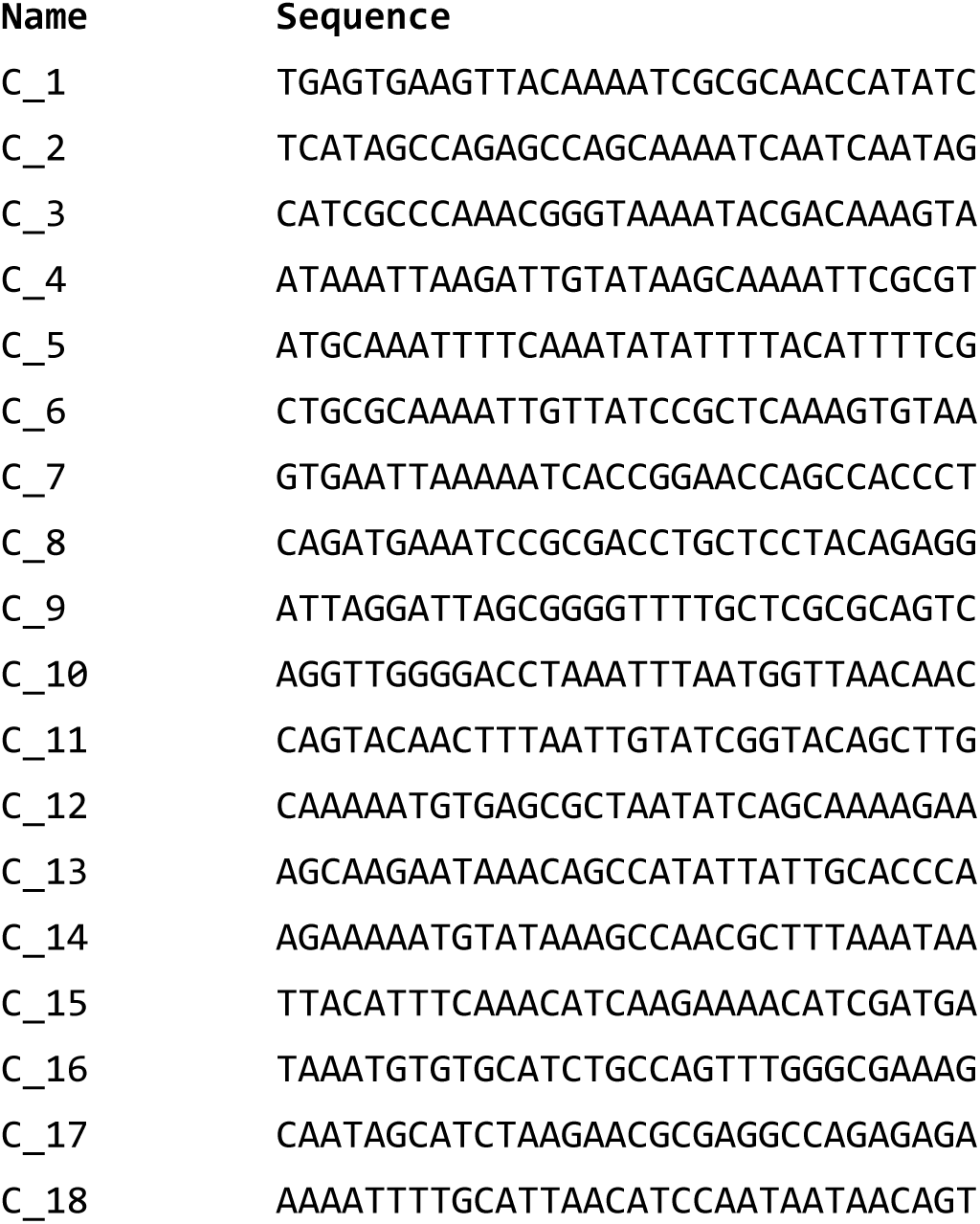
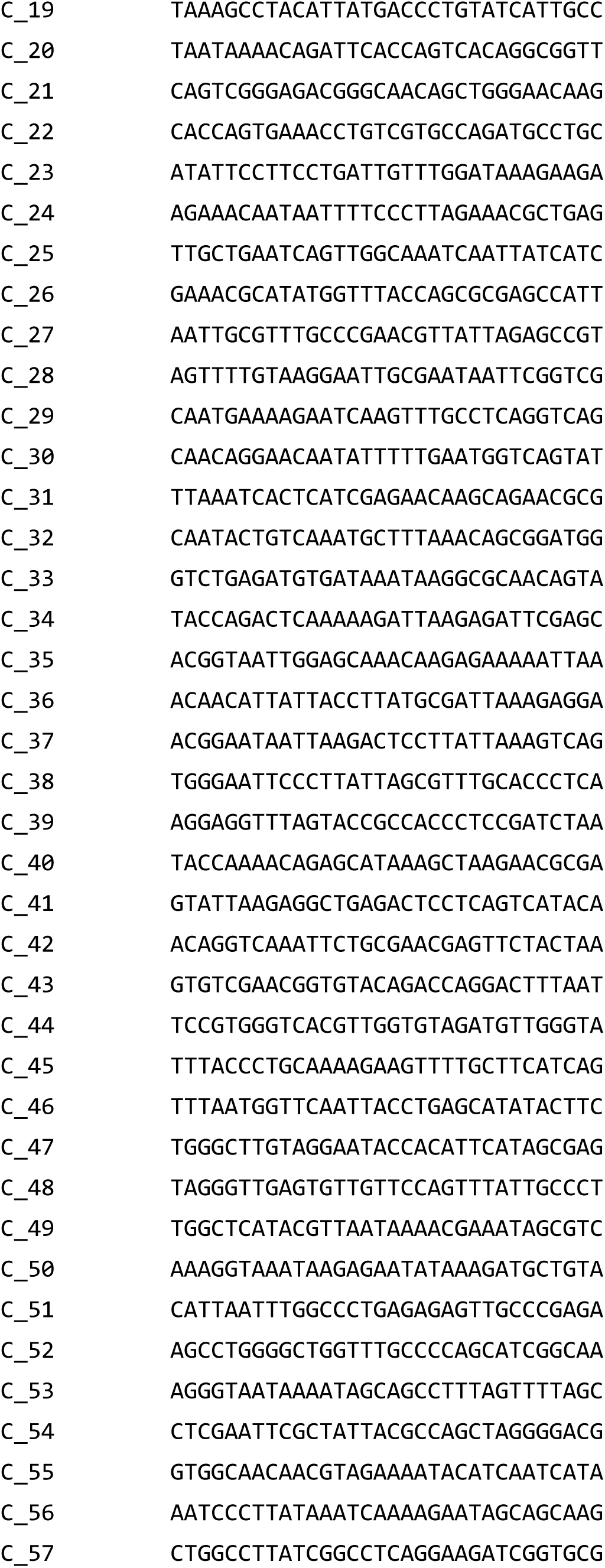
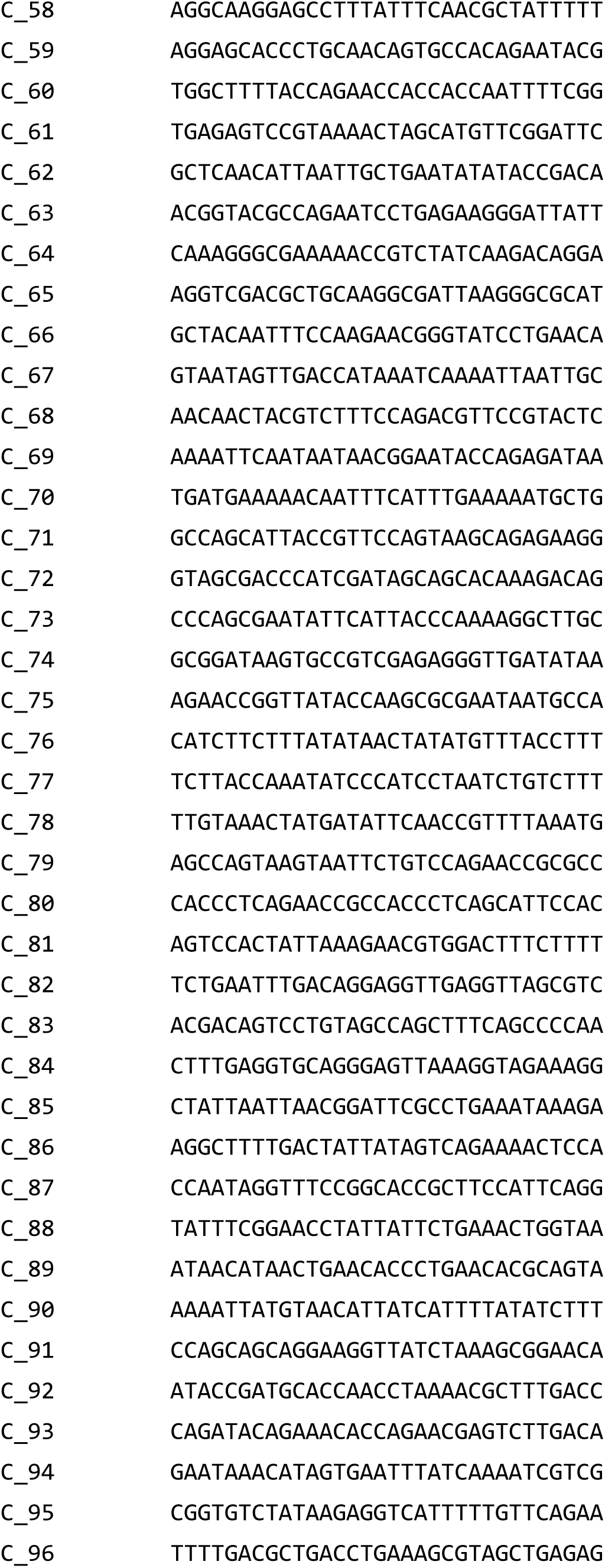
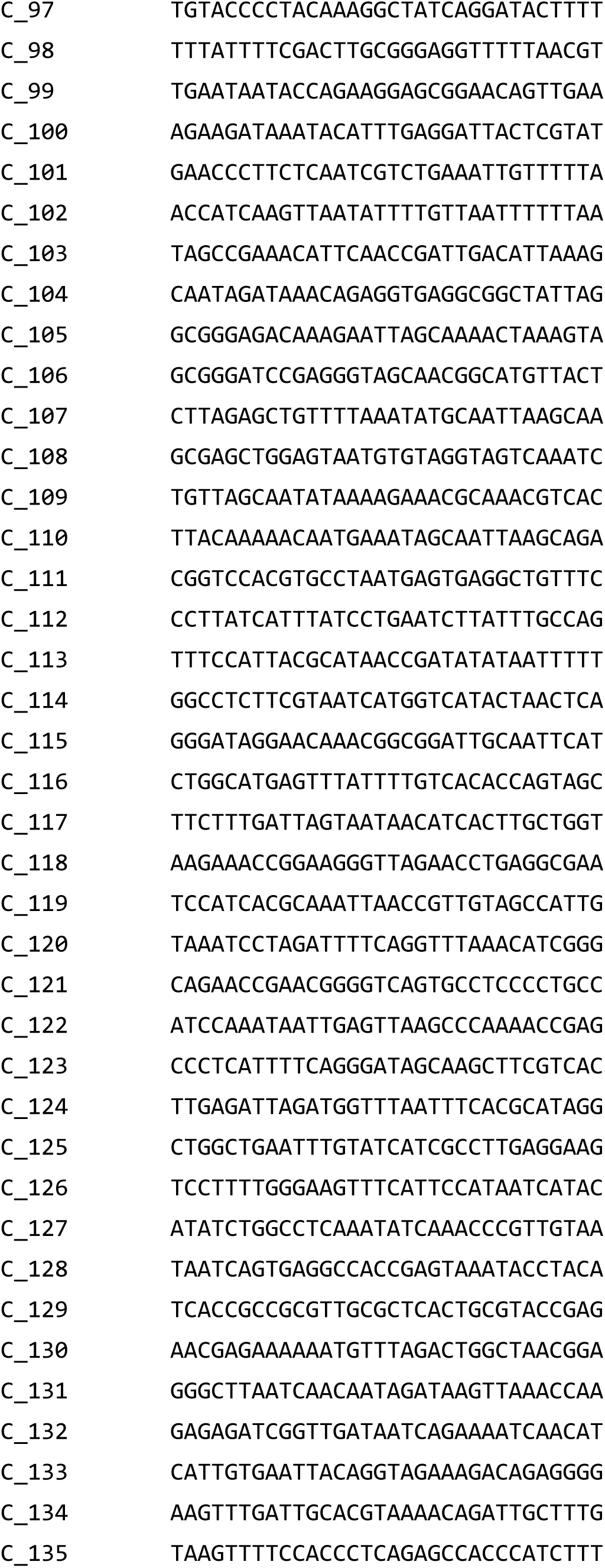
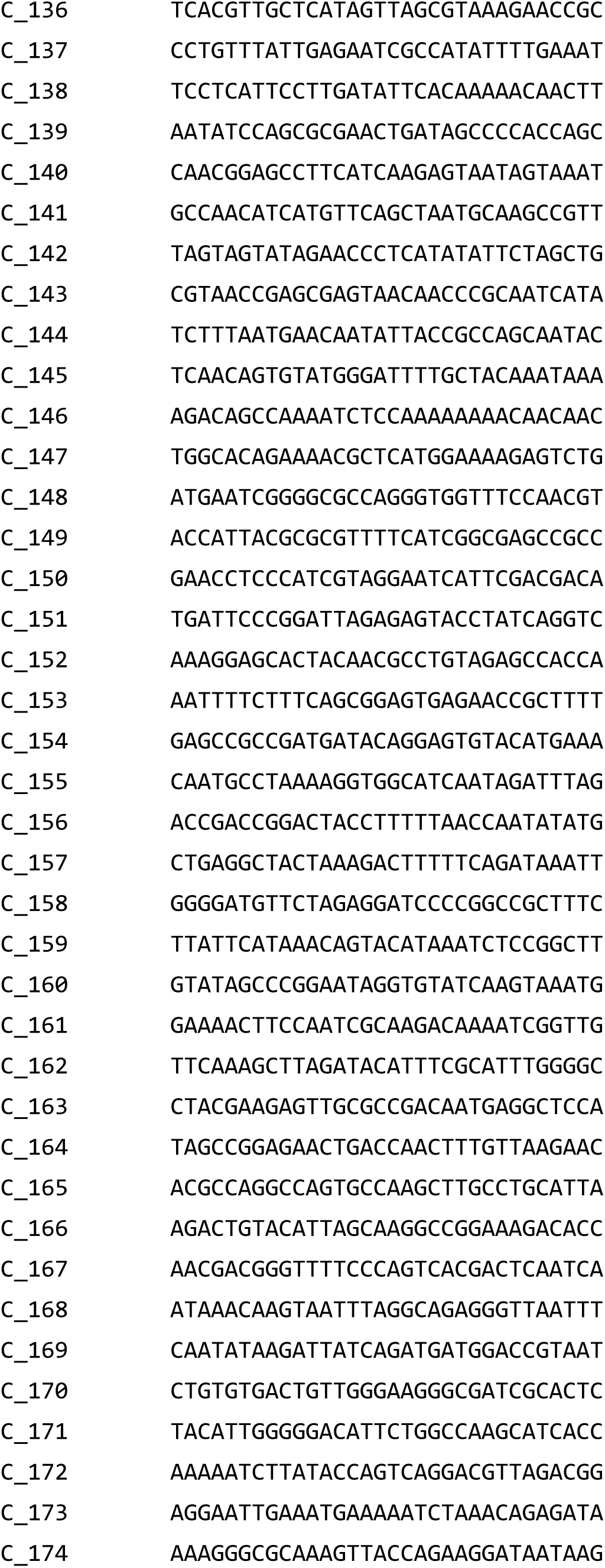
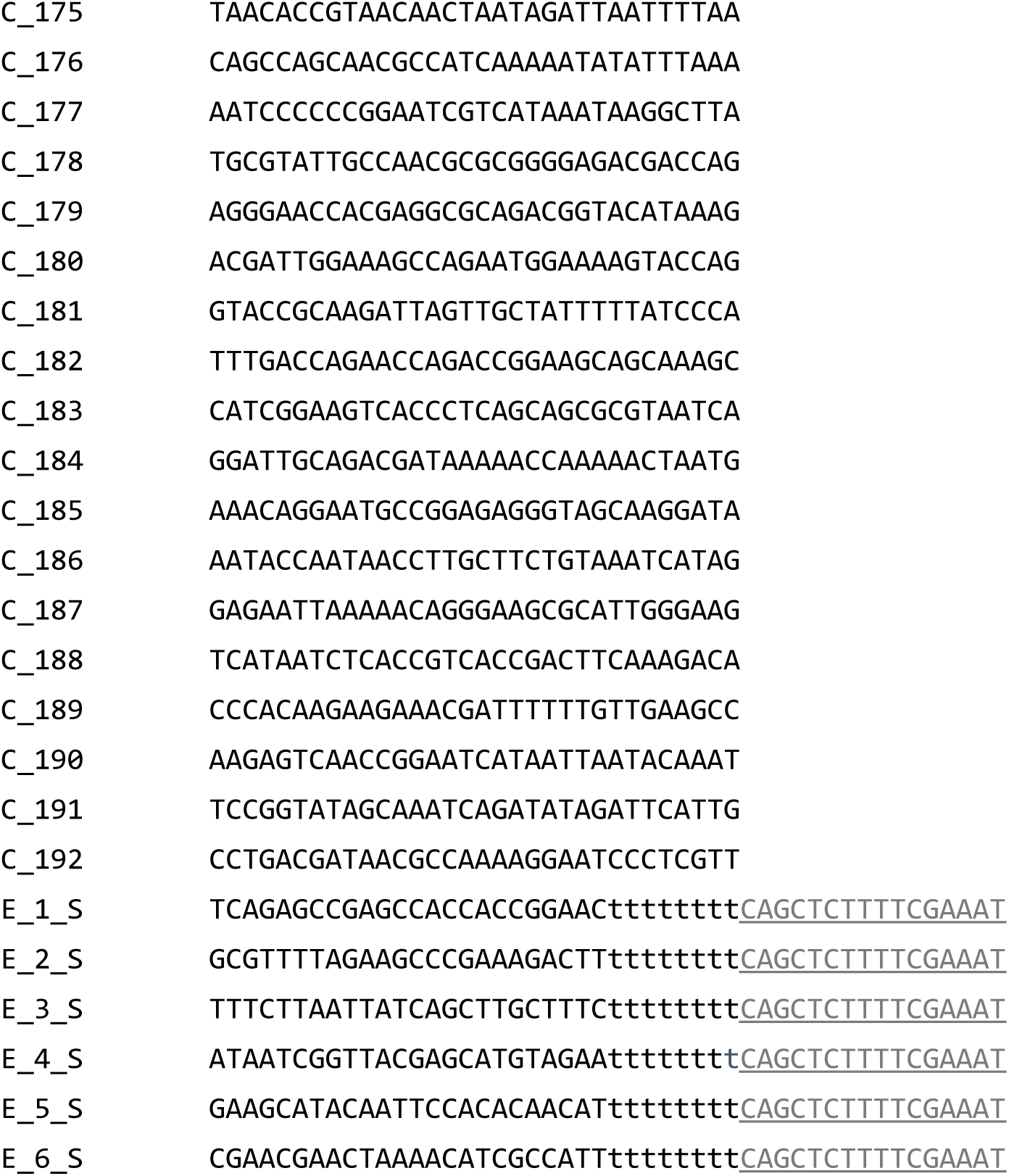

## Supplementary note 4

### HHR activity gels

In this study we used gel electrophoresis to measure various HHR reaction kinetics. Here several representative gels are shown.

**Figure S8.**
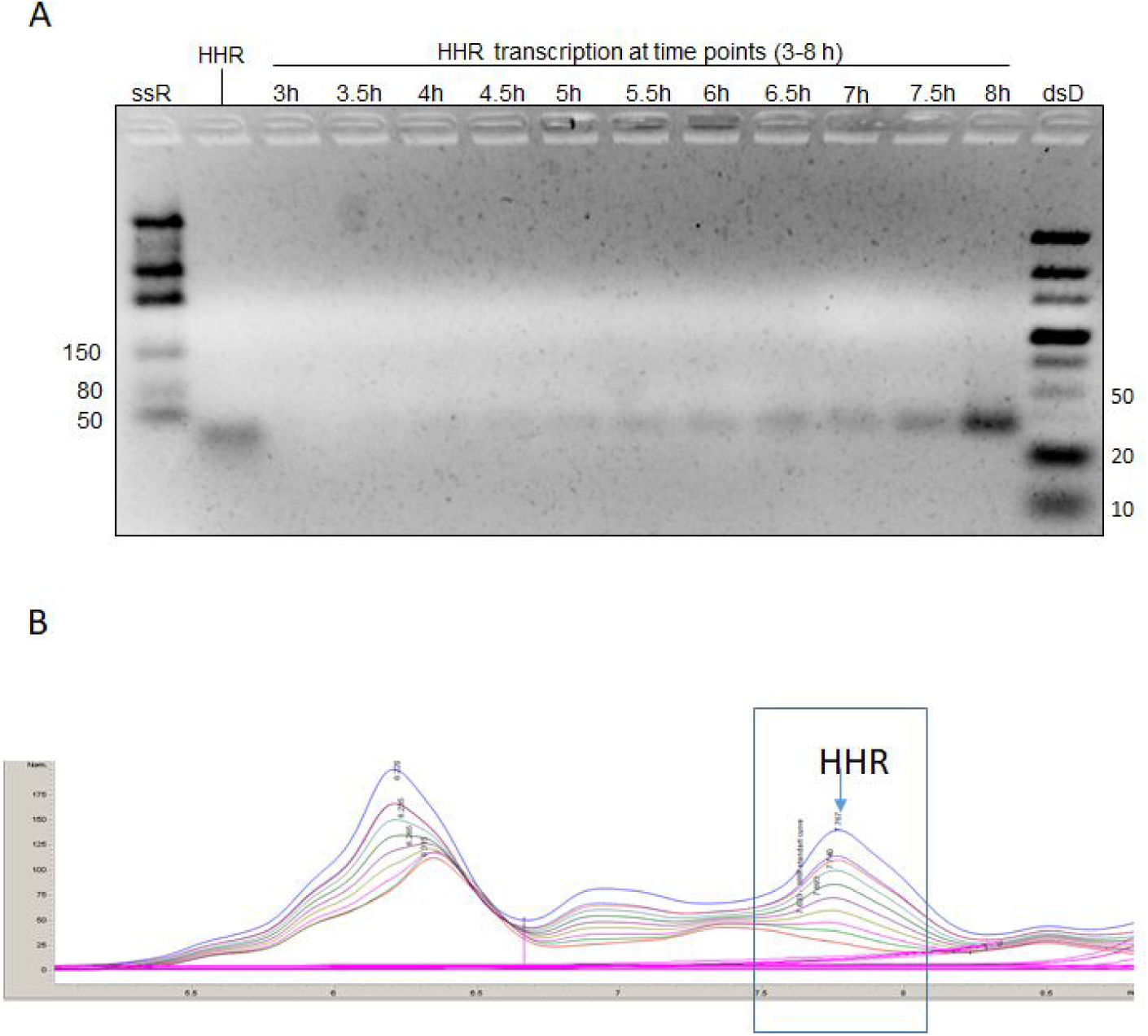
Transcription of HHR from assembled automata. Transcription reactions containing equal concentration of 3 µM of L, S, R(L), R(S), supplemented with dNTPs, NTPs, *Bsu* DNA polymerase and T7 RNA polymerase were incubated at 37°C. Following 3h of incubation, aliquots were drawn in 30 min intervals. **A,** HHR fractions were purified by HPLC and loaded on 4% TBE-agarose gel. **B,** HPLC analysis of the transcription aliquots post 3h (red), 3.5h (green), 4h (pink), 4.5h (beige), 5h (purple), 5.5h (dark green), 6h (turquoise), 6.5h (dashed blue), 7h (dashed red) and 7.5h (blue) post 37°C incubation.

**Figure S9.**
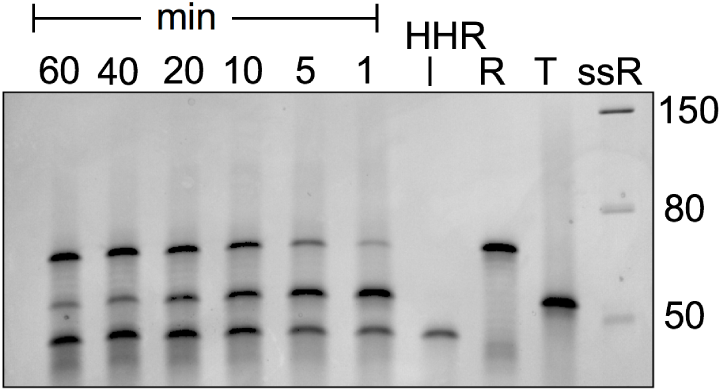
Kinetics of target digestion by HHR. Cleavage reaction was performed with equal concentration of 1.25 µM of target and HHR and incubated at 37°C, aliquots were drawn after 1,5,10,20,40,60 min and loaded on 10% TBE-urea PAGE.

**Figure S10.**
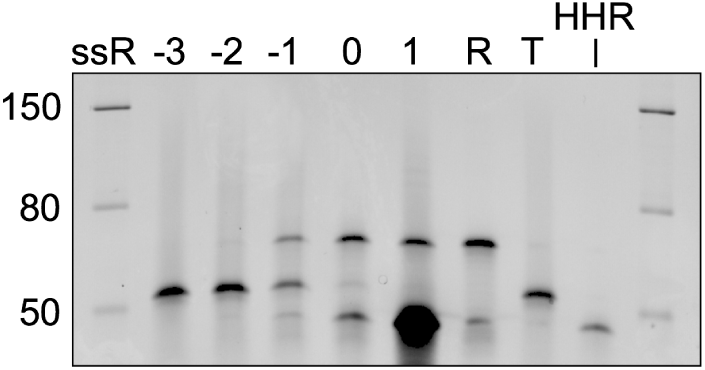
Dose response of target digestion by HHR. Cleavage reactions were performed with 1.25 µm of target and various concentrations of HHR with 10 fold dilutions starting from 1.25×10^1^ µm (labeled as 1) and incubated at 37°C. After 1 h of incubation, the samples were loaded on 10% TBE-urea PAGE.

## Supplementary note 5

### Alternative designs of the automaton

#### Self-limiting automata (3WJ)

These automata are composed of two ssDNA (L,S) that have short complementary sequences to each other and to the target and only then, they can form a three way junction (3WJ) **(Fig. S11, S12)**. The L strand contains a T7 promoter sequence and a complementary sequence of the HHR. Upon the 3WJ formation, DNA polymerase can elongate the S strand (**Fig. S13**) that produces active ds T7 promoter which in the presence of T7 RNA polymerase transcribes HHR **(Fig. S14)**. In turn, the transcribed HHR cleavages the target and reduces the target amount **(Fig S15)**. The reduction of the target diminishes the amount of the 3WJ and sequentially the HHR amount decreases.

**Table S2.**
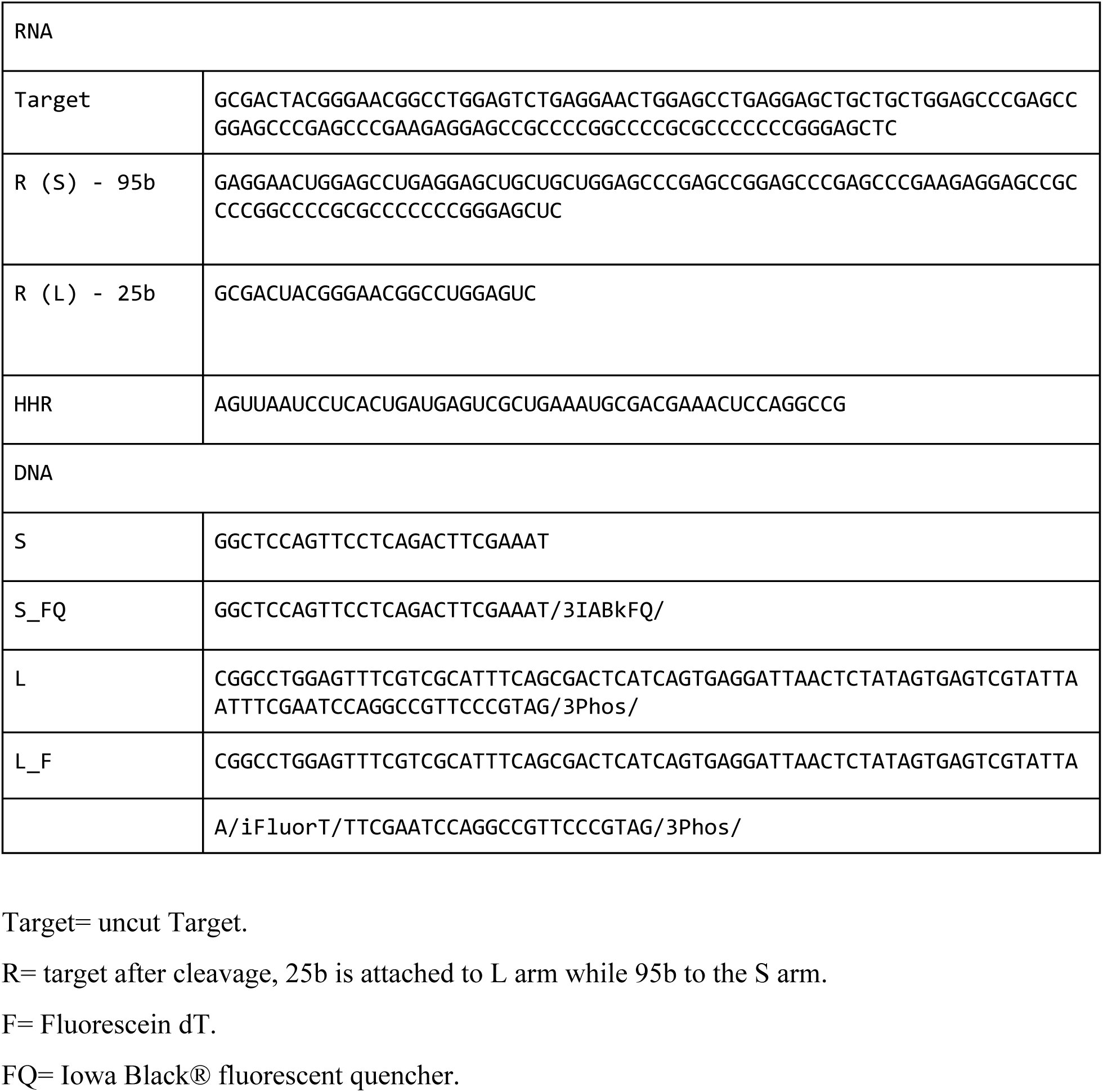
Self limiting automaton sequence list.

**Figure S11.**
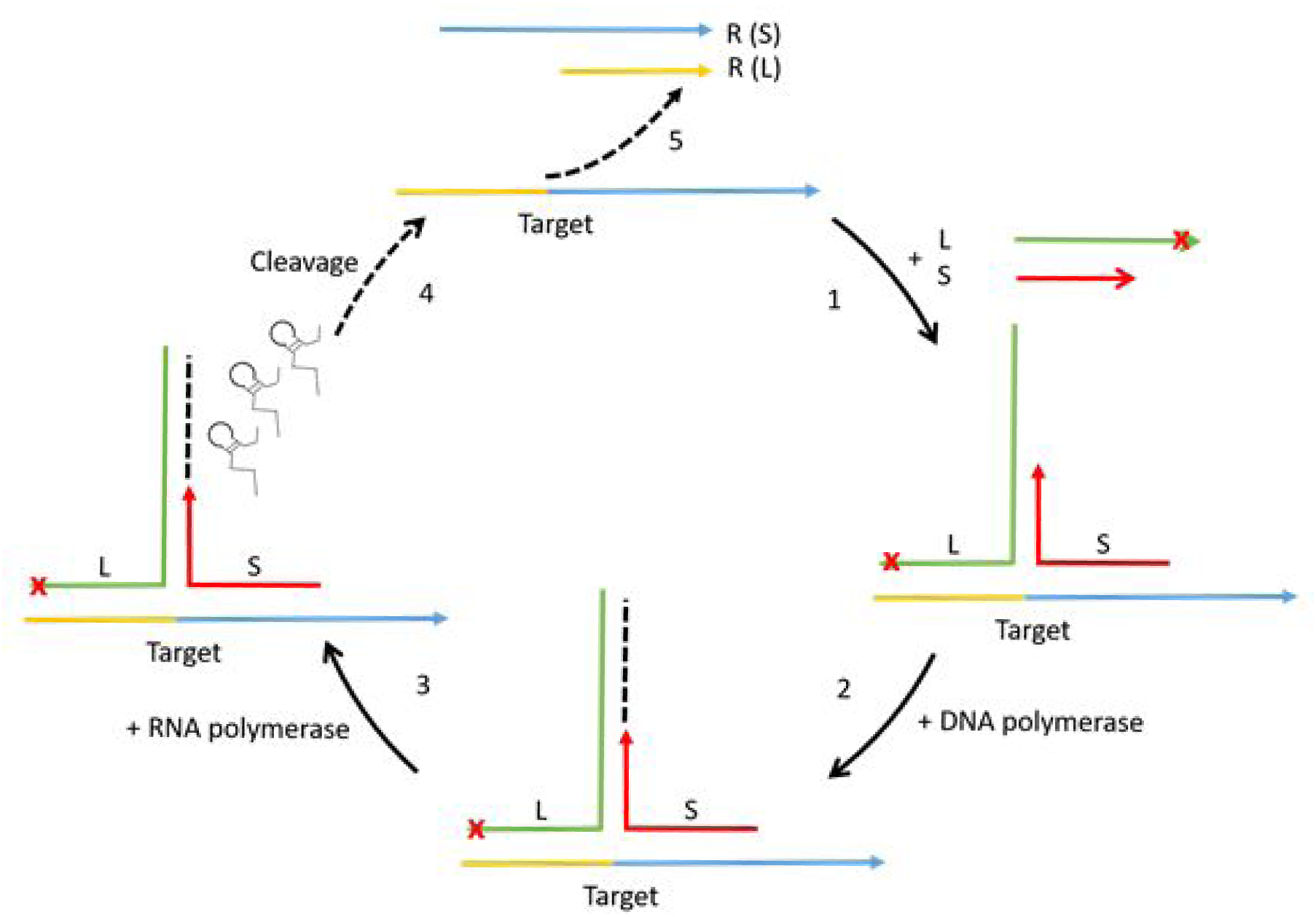
General scheme of the design, in five steps. (1) DNA oligonucleotides are added to the Target and form a 3WJ. (2) Addition of Bsu DNA polymerase to the reaction, leading to S elongation. (3) Addition of T7 RNA polymerase to the reaction, leading to HHR transcription. (4) HHR cleavage, follows by (5) components separation.

**Figure S12.**
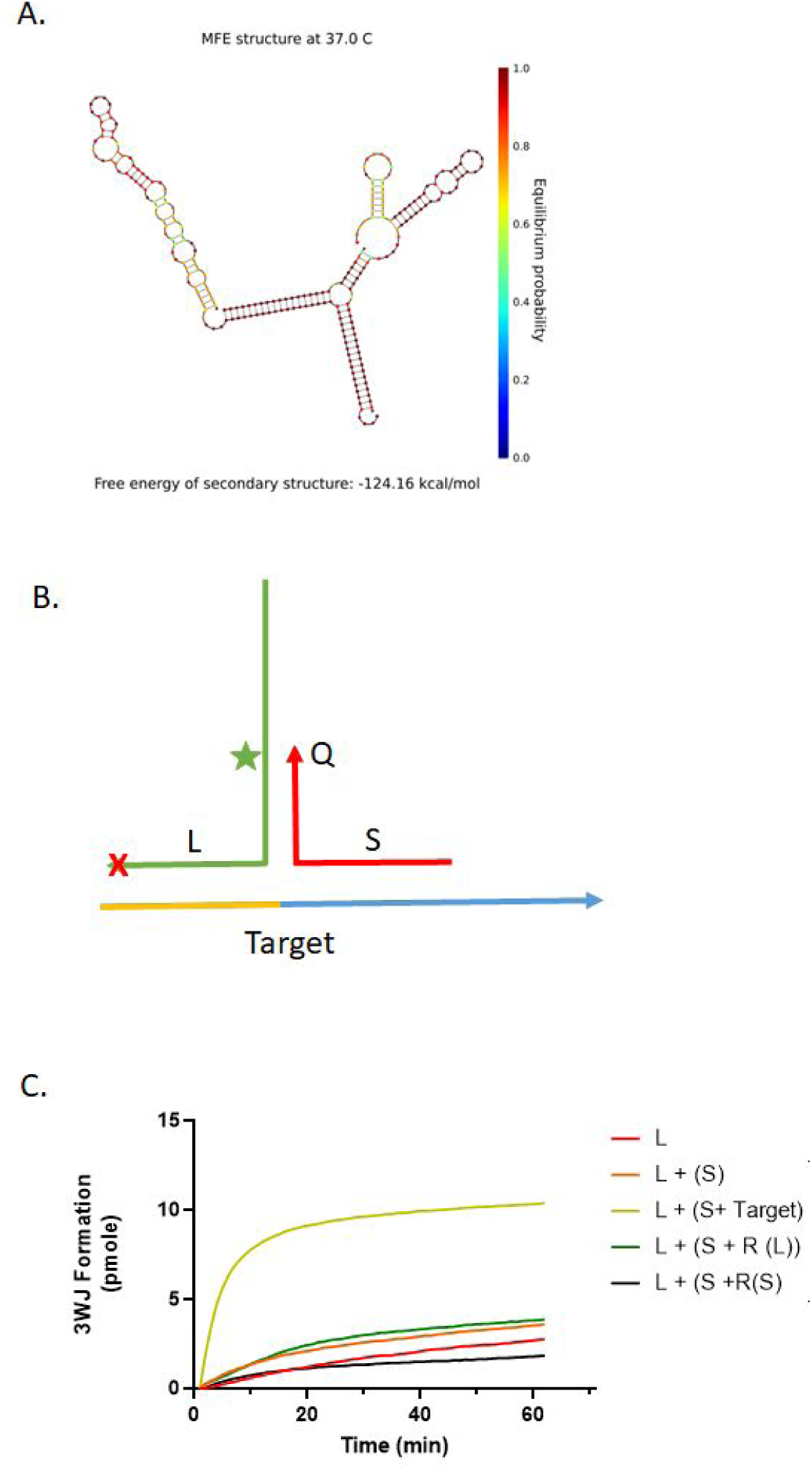
3WJ formation. **A,** Schematic representation of the 3WJ. **B,** Scheme of the design for testing 3WJ formation; internal fluorescent T was added to L, and a Iowa black fluorescent quencher was added in the 3’ of S. **C,** Following 3WJ formation using RT-PCR machine; initially the fluorescence of 1µM of L-F was followed every 1 min using the FAM channel for 10 min, then the rest of the oligos (S, Target, R(L), R(S)) were added to the reaction accordingly to final concentration of 1 µM. The fluorescence was measured every 1 min for an additional 60 min using the FAM channel. Data was normalized to baseline fluorescence and calculated as the number of assembled automaton molecules.

**Figure S13.**
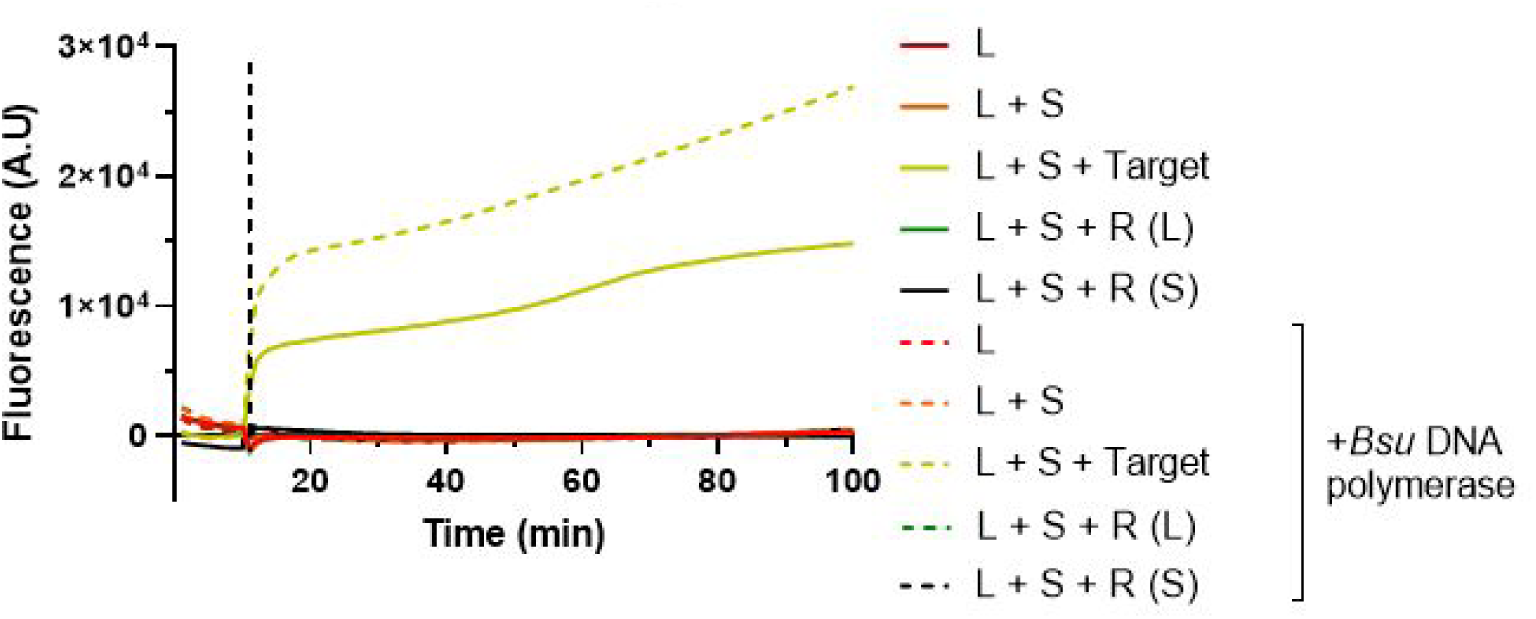
Complementary (S) strand synthesis. Following strand synthesis by addition of EvaGreen® using RT-PCR machine; fluorescence was followed using the SYBR channel, every 1 min. Samples were incubated with (dashed) and without (solid) Bsu DNA polymerase. Initially, all the oligos (S, Target, R(L), R(S)) with a final concentration of 1µM, EvaGreen® and dNTPS were added to the reaction and the fluorescence was followed every 1 min using the FAM channel for 10 min. Then Bsu was added to the reaction. The fluorescence was measured every 1 min for an additional 90 min using the FAM channel.

**Figure S14.**
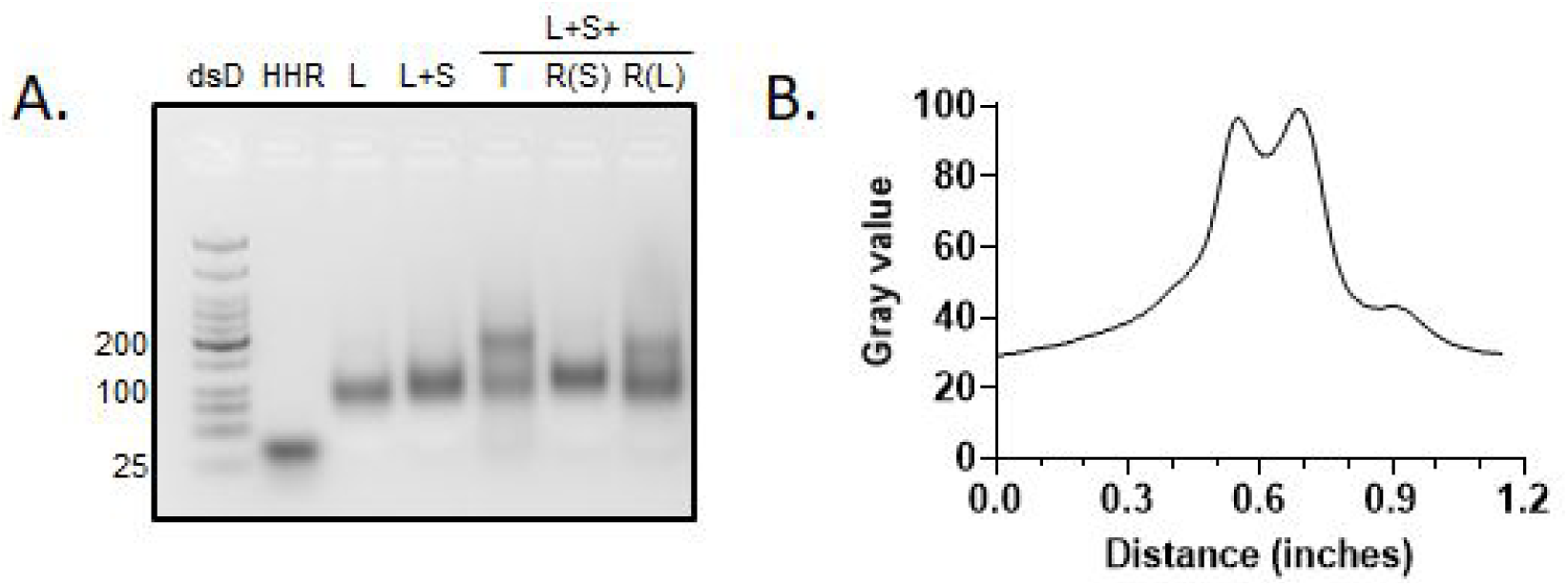
Transcription of HHR. **A,** Transcription reactions containing equal concentration of 3 µM of all the oligos, supplemented with dNTPs, NTPs, *Bsu* DNA polymerase and T7 RNA polymerase were incubated at 37°C for 3 h and then run in 3% TAE-agarose gel. **B,** Analysis of bend intensity profile of the transcription sample (lane 5, L +S +Target) by ImageJ. Shoulder at ∼0.9 in is *de-novo* transcribed HHR.

**Figure S15.**
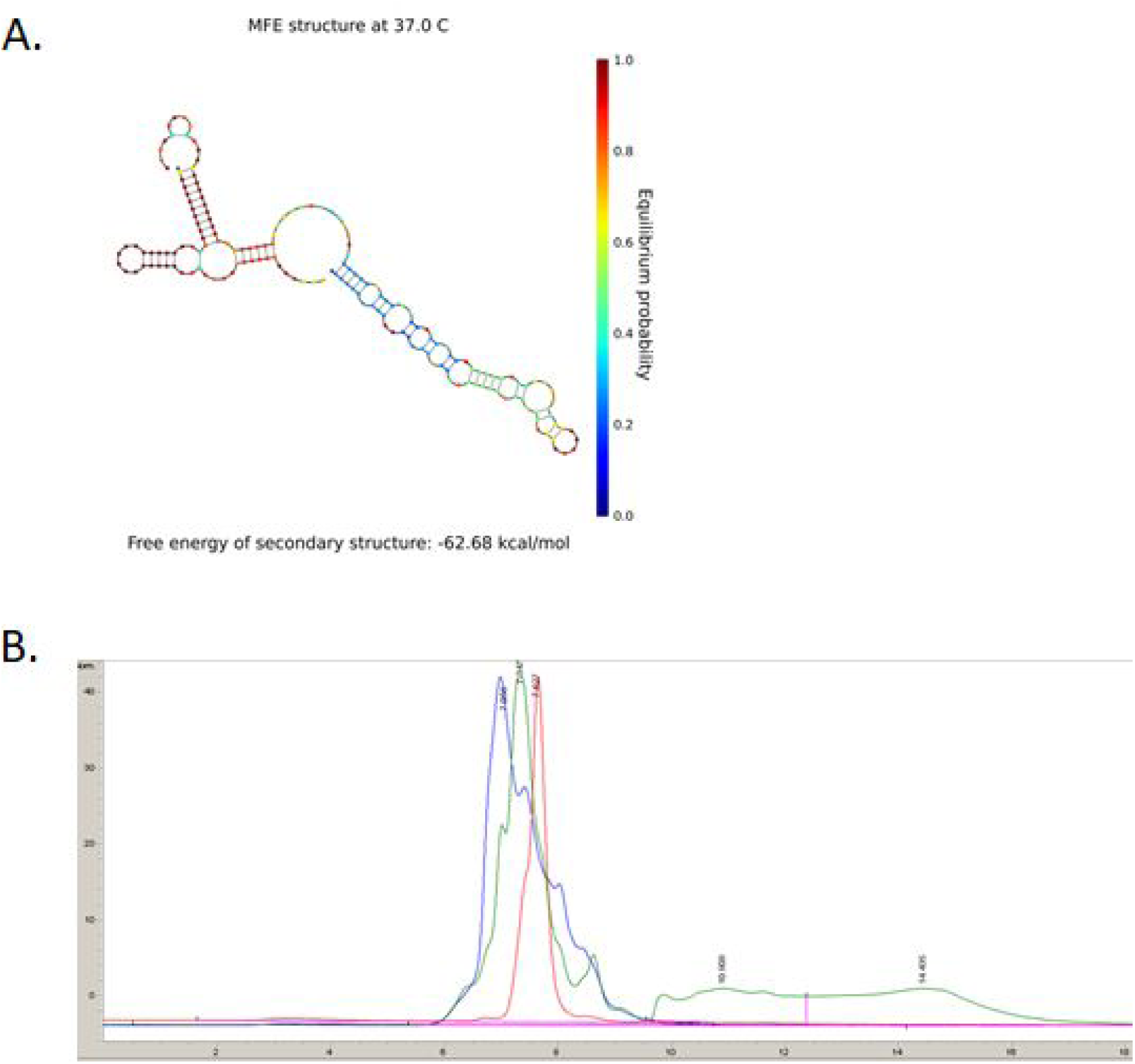
Cleavage of target by HHR. **A,** Schematic representation of the HHR binding to the target. **B,** Testing cleavage of target by transcribed HHR using HPLC; Cleavage reaction was performed with equal concentration of 1µM of Target and HHR and incubated at 37°C for 1 h. Target before cleavage (blue), HHR (red) and cleavage reaction (green).

#### Design of a sensing automaton

The idea is to build a molecular automaton, with complex recognition and sensing capabilities. First, the automaton assembles only in the presence of its specific target RNA (R) just like the prototype automaton. Upon assembly the sensor automaton is inactive as long as it doesn’t encounter the activating molecule.

The sensor automaton design is based on the prototype automaton **(Supplementary note 1, Table S3)**, and works via toehold mediated strand displacement mechanism. Two changes were done: first, L strand was extended at 5’ with <u>thoehold-T7</u> (10 and 17 bases respectively). This addition leads to conformational changes in L strand, which result in a formation of a hairpin loop, making the T7* and the HHR* sequences less accessible. The new L is denoted by sL. Second, <u>T7*-toehold*</u> sequence was added to 5’ of R strand **(Fig. S16A, B)**. The new L and R are denoted by sL and sR, respectively. The assembly of the sensor automaton was analyzed by NUPACK^37^ simulations **(Fig. S16C, D)** and verified by *in-vitro* assembly assay following gel electrophoresis as described in the materials and method section. **(Fig. S17).**

Another sensing mechanism can be an aptamer-target and ligand-protein interactions, where such interactions lead to unwinding of the L strand, and activation of the sensor automaton.

**Figure S16.**
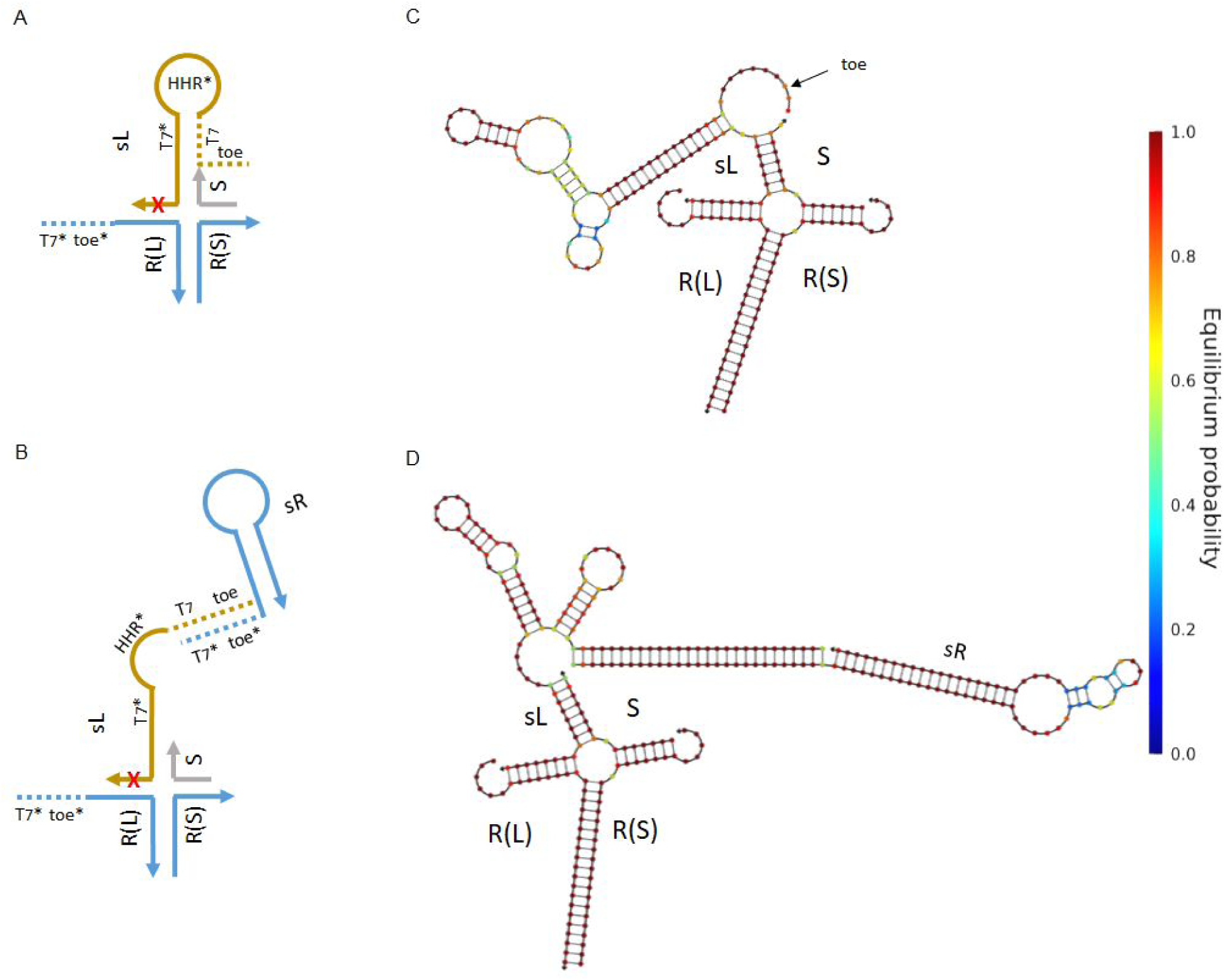
Design and assembly simulation of the sensing automaton. **A-B,** Schematic illustration of an inactive and active sensor automaton, respectively. **C-D,** Nupack simulation of an inactive and active sensor automaton, respectively.

**Figure S17.**
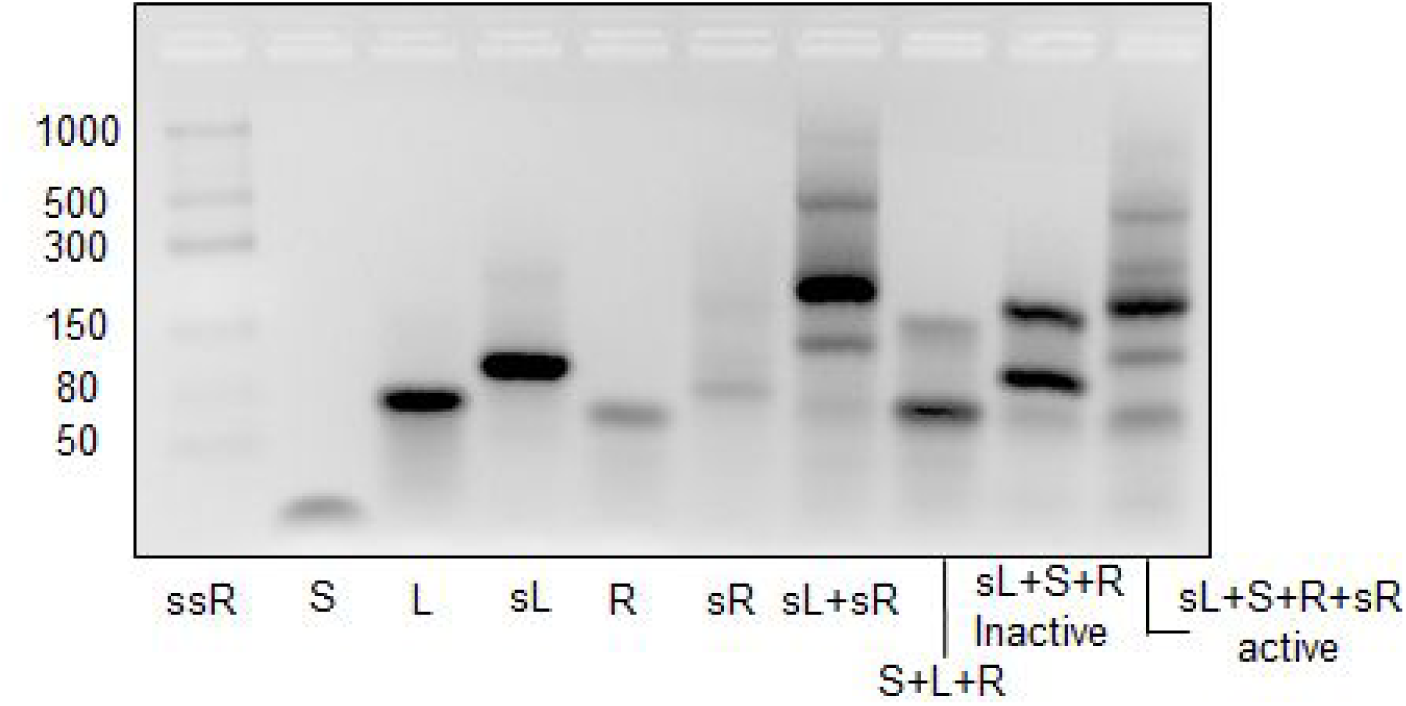
*In-vitro* assembly of the sensing automaton. Gel electrophoresis of fully assembled prototype and sensor automata. The sensor automaton checked in two possible states: inactive and active.

**Table S3.**
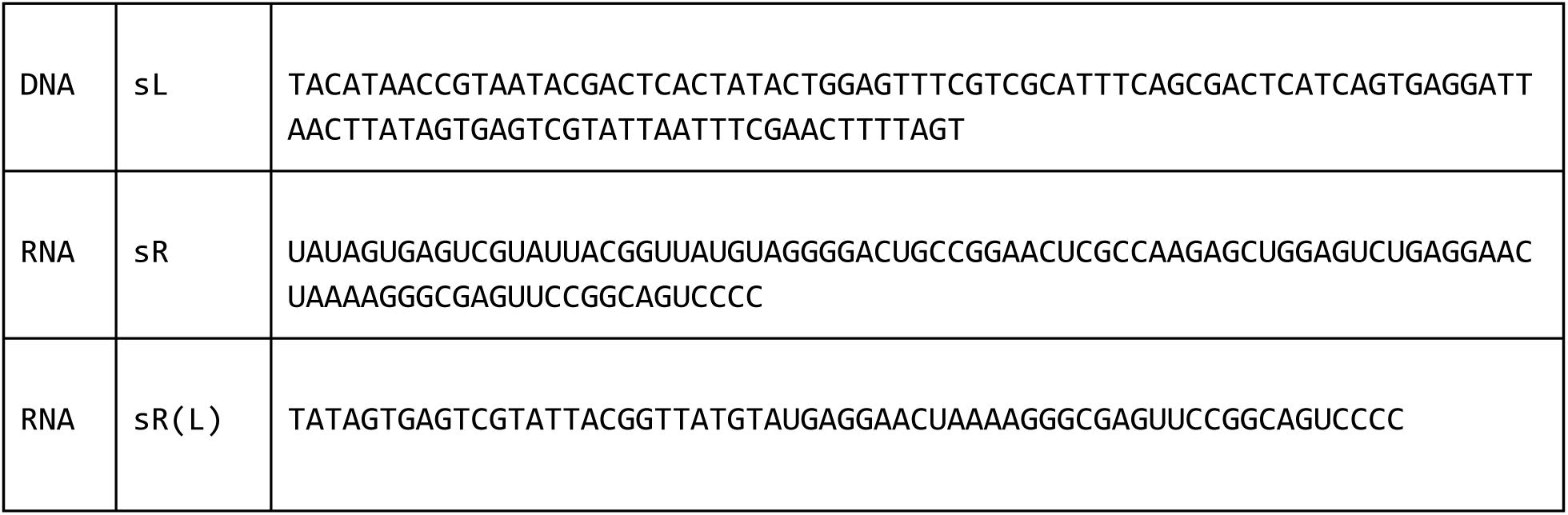
Sensing automaton sequence list. S and R(S) remain as in the prototype automaton design.

## Supplementary note 6

### Sf9 culture system, delivery, and infection

We made use of *Autographa californica* multicapsid nucleopolyhedrovirus (AcMNPV) and a commercially-available cell line, Sf9, obtained from *S. frugiperda*, which are susceptible to infection by this virus. Sf9 cells can be grown as adherent cells, as well as suspended, enabling us to mimic *in-vivo* infection of an insect naturally possessing cells as part of a tissue or single ones located at the insect’s hemocoel.

In order to follow infection rate, GFP was cloned under the very late strong promoter of polyhedrin (PH). The virus was amplified by 2 rounds of amplification and the virus stock titer was determined to be 2*10^8^ pfu/ml, kept protected from light at 4°C (titer was determined by End Point Dilution technique).

For the purpose of calibrating our model system in terms of adherent vs. suspended cells, as well as infection progress, we infected adherent Sf9 cells (seeded in regular 6 well plate, 1*10^6^ cells/well) or suspended (seeded in low-binding 6 well plate, 1*10^6^ cells/well). We observed similar behavior for adherent and suspended cells at various MOI’s and a distinct infection rate for each MOI as can be observed for 72 h post infection (**Fig. S18**).

Time dependent kinetics were determined for each MOI by harvesting infected wells at different time points post infection, a MOI dependent kinetics were observed (**Fig. S19**).

Since our automaton targets RNA coding for GP64, which is being expressed starting at the early stages of infection, as opposed to our reporter gene GFP which is being expressed under the very late promoter of polyhedrin, we needed to assess the actual definite amount of GP64 RNA for a specific time point post infection and a specific MOI (**Fig. S20**). GP64 RNA copy number served us for assessing the therapeutic dose of automaton we delivered as a viral infection treatment.

Automaton and enzyme payload delivery was tested by a non-lipophilic agent (Mirus-*Trans*IT-X2® Dynamic Delivery System for *CRISPR/Cas9 Ribonucleoprotein (RNP) + DNA Oligo (ssODN) Delivery*) and by liposome (liposome kit, L-4395-1VL, Sigma-Aldrich).

In order to assess the size distribution of our liposomes, we made use of DLS Malven Nano-ZS, ZEN3600 DLS instrument. Samples of liposomes were prepared according to manufacturer’s recommendations. The vast majority of the liposomes are ∼1 µm diameter (**Fig. S21**).

Efficiency of enzyme delivery by liposome was done by fluorescently labeling a T7 RNA polymerase and test treated vs. non-treated cells by FACS (**Fig. S22**).

**Figure S18.**
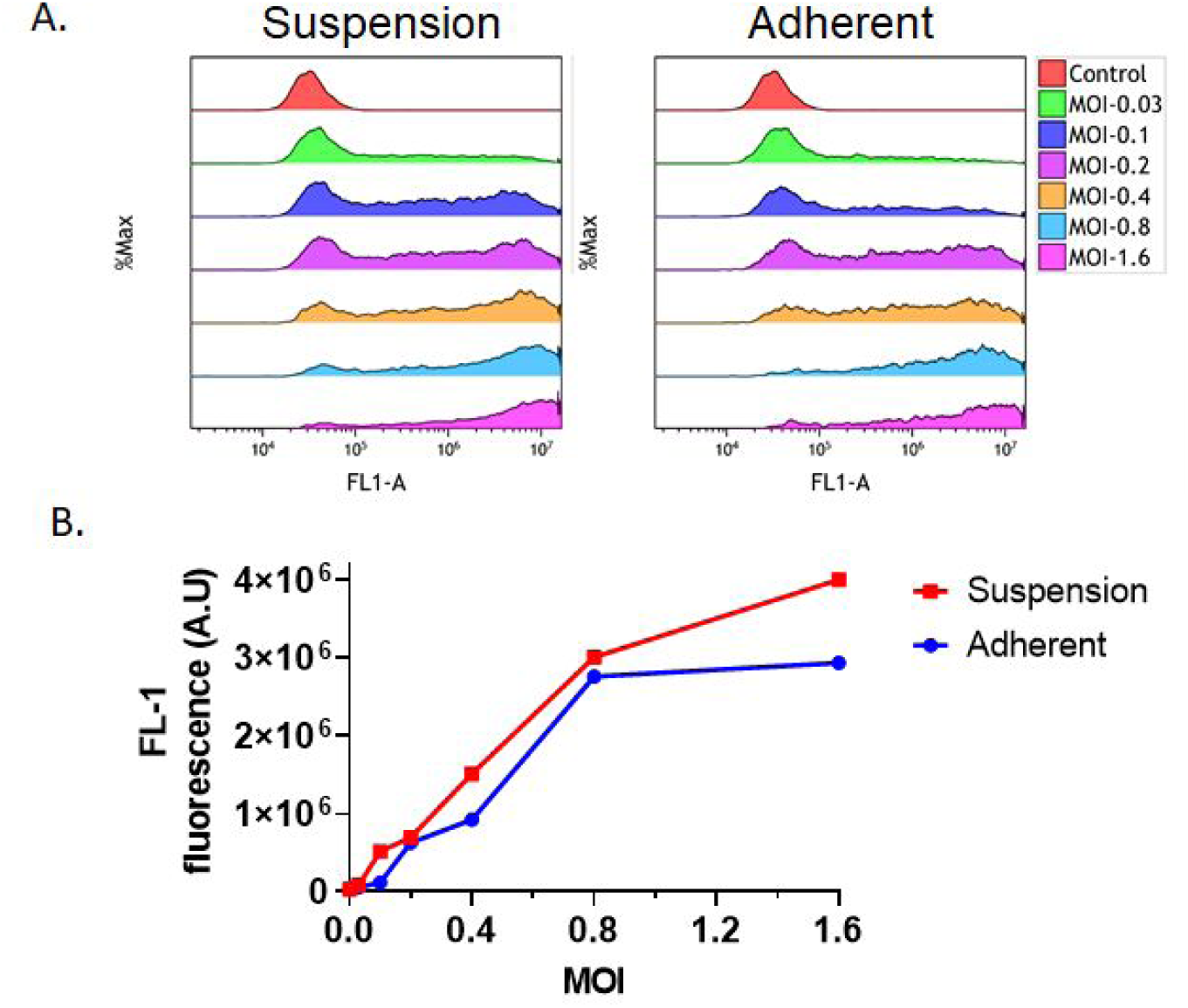
Calibration of AcMNPV infection of suspended and adherent cell culture of Sf9 cells. Adherent and suspended cultures of 1×10^6^ Sf9 cells were infected with AcMNPV in various MOI (0.03, 0.1, 0.2, 0.4, 0.8 and 1.6). After 72 h post infection, cells were harvested and GFP signal was monitored using flow cytometry. **A,** GFP signal distribution in the population of suspended (left panel) and adherent (right panel) cells infected with increasing MOI. **B,** Median GFP signal in suspended (red line) and adherent (blue line) cells culture infected with increasing MOI.

**Figure S19.**
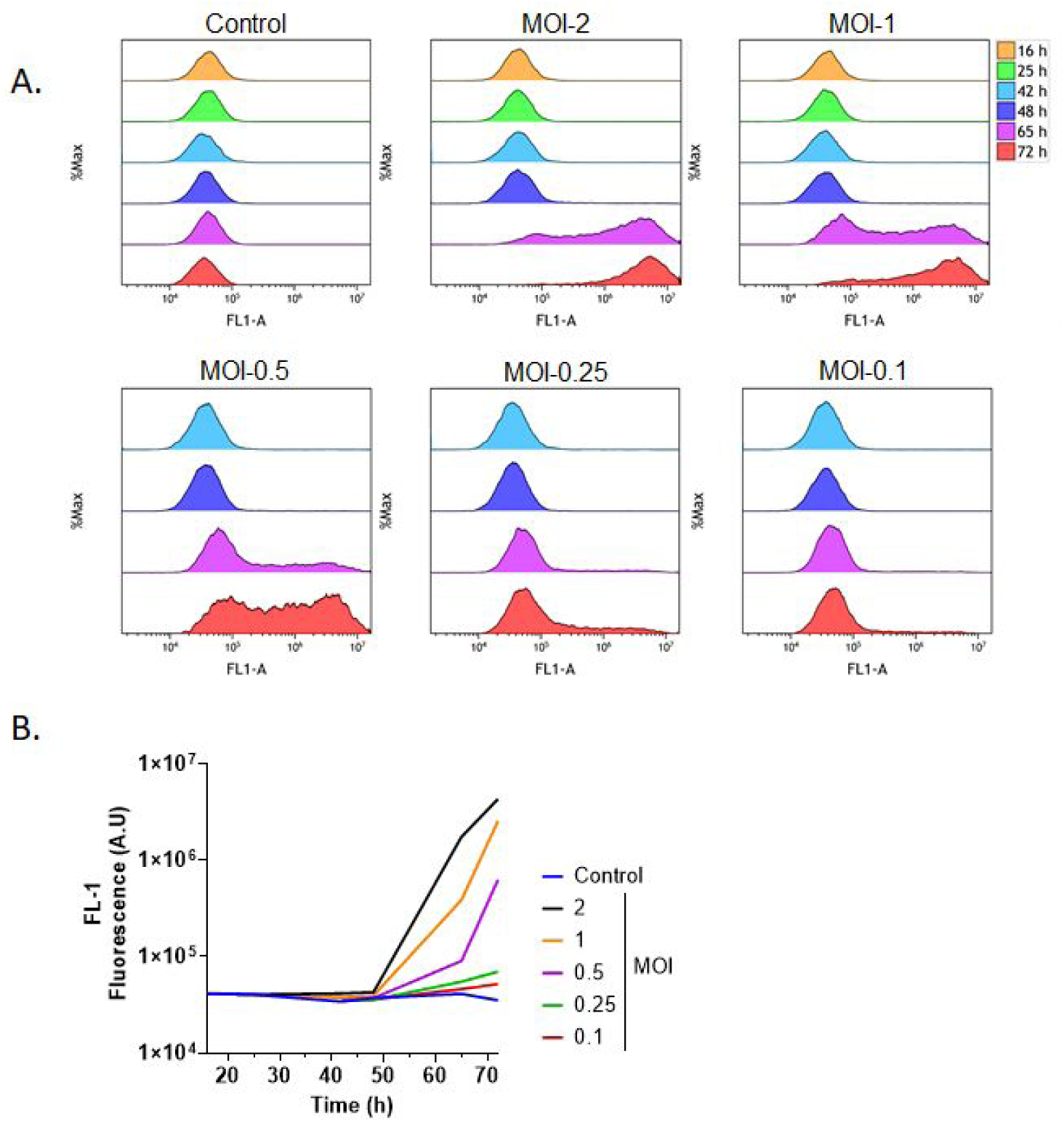
AcMNPV infection dynamics of adherent Sf9 cell culture. Adherent culture of 1×10^6^ Sf9 cells was infected with AcMNPV in various MOI (0.1, 0.25, 0.5, 1 and 2). After 16, 25, 42, 48, 65 and 72 h post infection, cells were harvested and GFP signal was monitored using flow cytometry. **A,** GFP signal distribution in the population of adherent Sf9 cell culture that were infected with increasing MOI over time. **B,** Median GFP signal in adherent Sf9 cells culture infected with increasing MOI over time.

**Figure S20.**
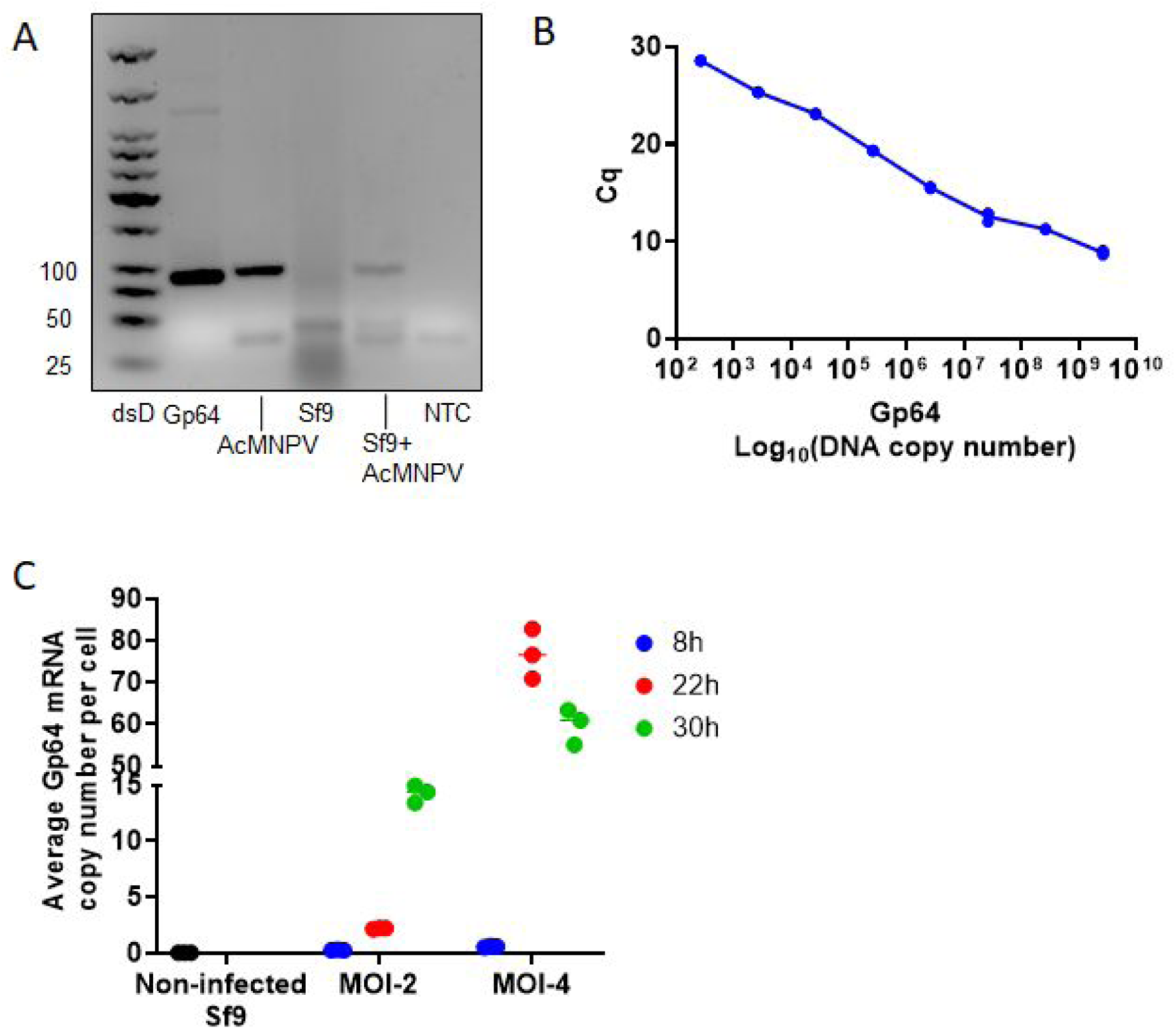
GP64 RNA quantification during infection. **A,** To test primer specificity to GP64, a PCR was conducted using specific primers that amplify 92 bp region in the gene. Amplification can be detected only in AcMNPV lysate, Sf9 cells infected with AcMNPV and in the control plasmid cloned with the AcMNPV gene GP64. **B,** Correlation between the Cq to infinite number of GP64 molecules a qPCR was performed using known amounts of GP64 molecules with 10 fold dilutions starting from 2.5×10^9^ molecules to 250 molecules. **C,** Adherent culture of Sf9 cells was infected with AcMNPV in MOI 2 or 4. After 8, 22 and 30 h post infection, cells were harvested, total RNA was extracted and reverse-transcribed. GP64 expression was determined by qPCR relative to a control with known amounts of GP64 molecules.

**Figure S21.**
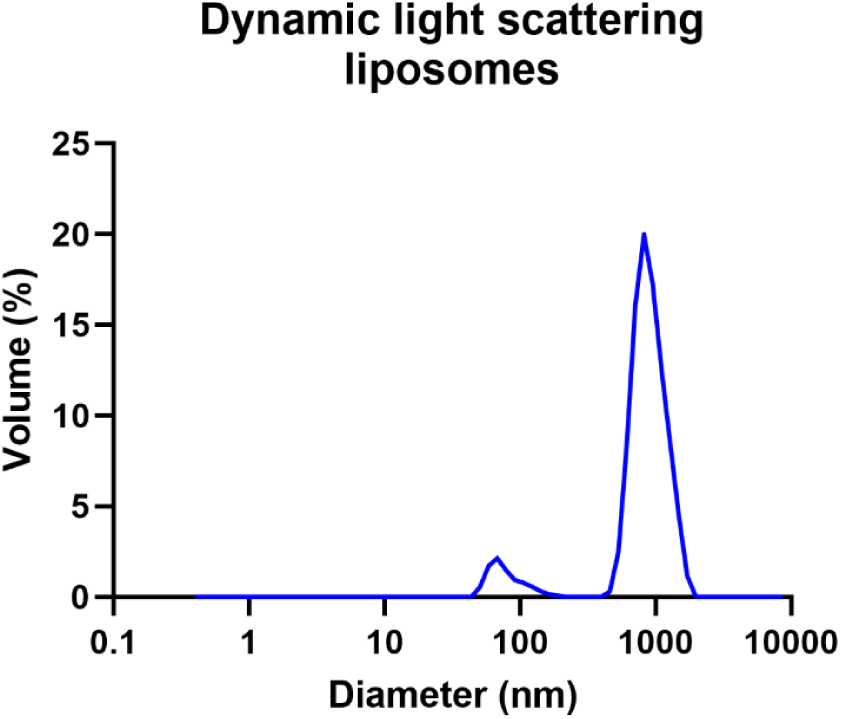
Liposomes size distribution. Liposome size distribution was measured using a dynamic light scattering instrument. The average diameter of the liposomes is ∼1 µm.

**Figure S22.**
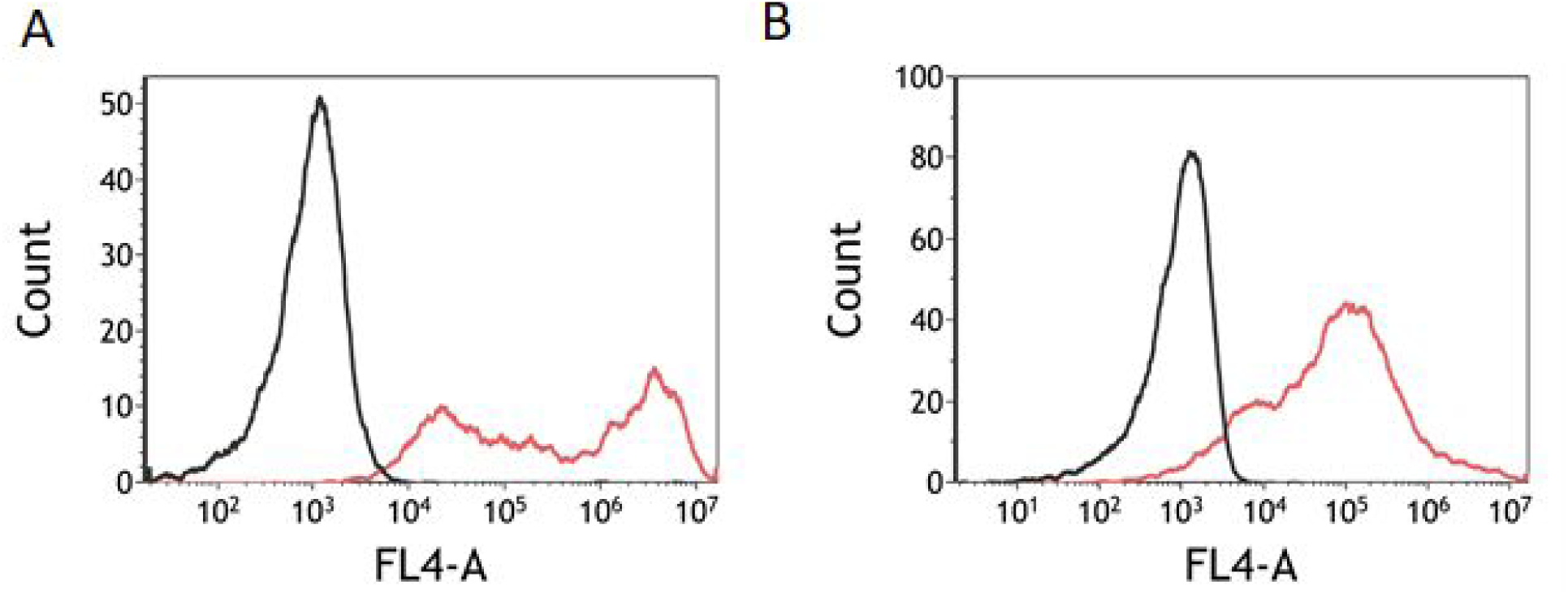
Transfection of labeled T7 polymerase and DNA. T7 polymerase and the S-arm of the predator were labeled with Alexa Fluor 647 and Cy5, respectively, packed into liposome and transfected adherent Sf9 cells. After 1h post transfection the cells were harvested and fluorescence was measured using the FL4 channel via flow cytometry. Un-transfected cells (black) had a median fluorescence of 842 A.U, while the transfected cells with labeled T7 polymerase (**A,** red line) and S-arm (**B,** red line) had a median fluorescence of 381,281 A.U and 72,357 A.U, respectively (*P* < 10^-4^).

## Supplementary note 7

### Design and simulation of GP64 automaton

To test our concept in cells (Sf9) and *in-vivo* (Adult *Blaberus discoidalis*) we designed a molecular automaton that targets the GP64 gene in AcMNPV. GP64 CDS sequence was downloaded from NCBI (NC_001623.1) and used Ribosoft^62^ to identify possible HHRs and cleavage sites. Four HHRs that were ranked first were chosen for further cleavage assays **(Fig. 3D).** The cleavage assays were performed on a GP64 transcripts obtained by in vitro-transcription reaction on a plasmid comprising a GP64 gene **(Fig. 3D,Fig. S23**). Based on the results and GP64 secondary structure as predicted by mFold^89^, we proceeded with HHR that targets GP64 at position 262-264 **(Fig. S24A).** A full molecular automaton comprising L, S and the cleaved GP64 as R was designed manually **(Table S4)**. L and S complement the RNA sequence **(Fig. S24B).** As in the prototype automaton, the L strand consists of a T7 promoter sequence, a complementary sequence of the HHR gene and a 3’ phosphorylation modification to prevent elongation from the L strand.

Assembly of the automaton was verified by NUPACK^37^ simulations. L and S undergo assembly to form a stable 4 way junction preferably when the target RNA is cleaved by the HHR, making the bases that participate in the hybridization more accessible **(Fig. S25)**. We verified that such a 4-way junction can be assembled in vitro, by using FAM labeled L arm and dark-FQ quencher S arm, and following the decrease in the fluorescence by RT-PCR. As R strand we used the R from the prototype design with different arms complementary to GP64 **(Fig. S26).** To increase the assembly and the stability of the automaton *in-vivo*, we elongated the L and S arms by 4 bases - ‘GATA’ was added to 3’ of L arm and ‘GACG’ was added to 5’ of S arm. In addition, we modified 7 bases in the complementary parts to the target with LNA bases, so the hybridization to the target RNA will be stronger.

**Figure S23.**
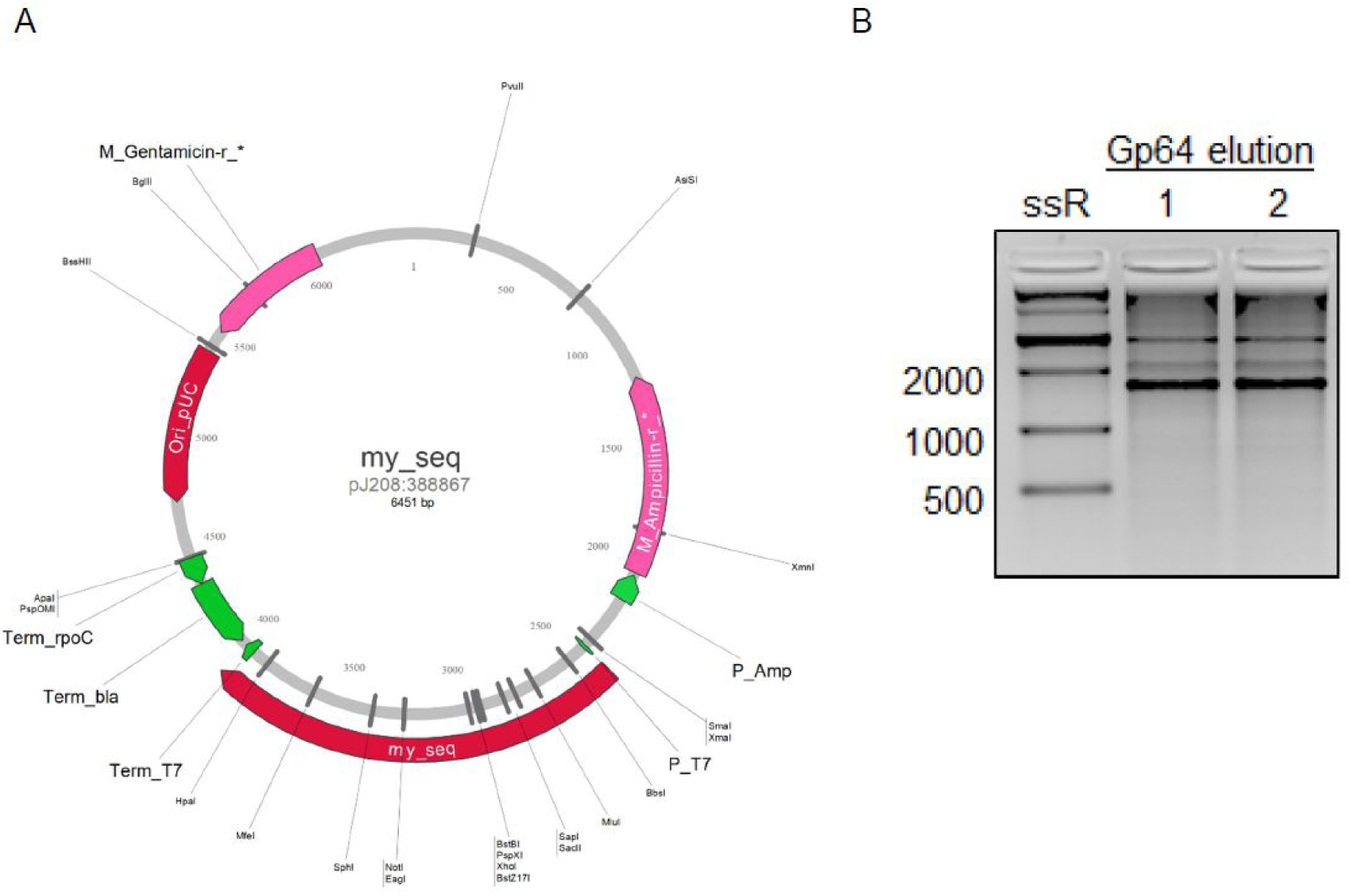
*In-vitro* transcription of GP64 RNA. **A** Plasmid map of AcMNPV GP64 cloned into a pJ209-ampR (ATUM) vector unter a T7 promoter. **B,** *In-vitro* transcription of GP64 RNA. GP64 RNA was transcribed from pJ208-ampR vector cloned with the AcMNPV GP64 gene under the T7 promoter. Transcribed GP64 were purified in two elution steps and loaded on 2% TBE-agarose gel.

**Figure S24.**
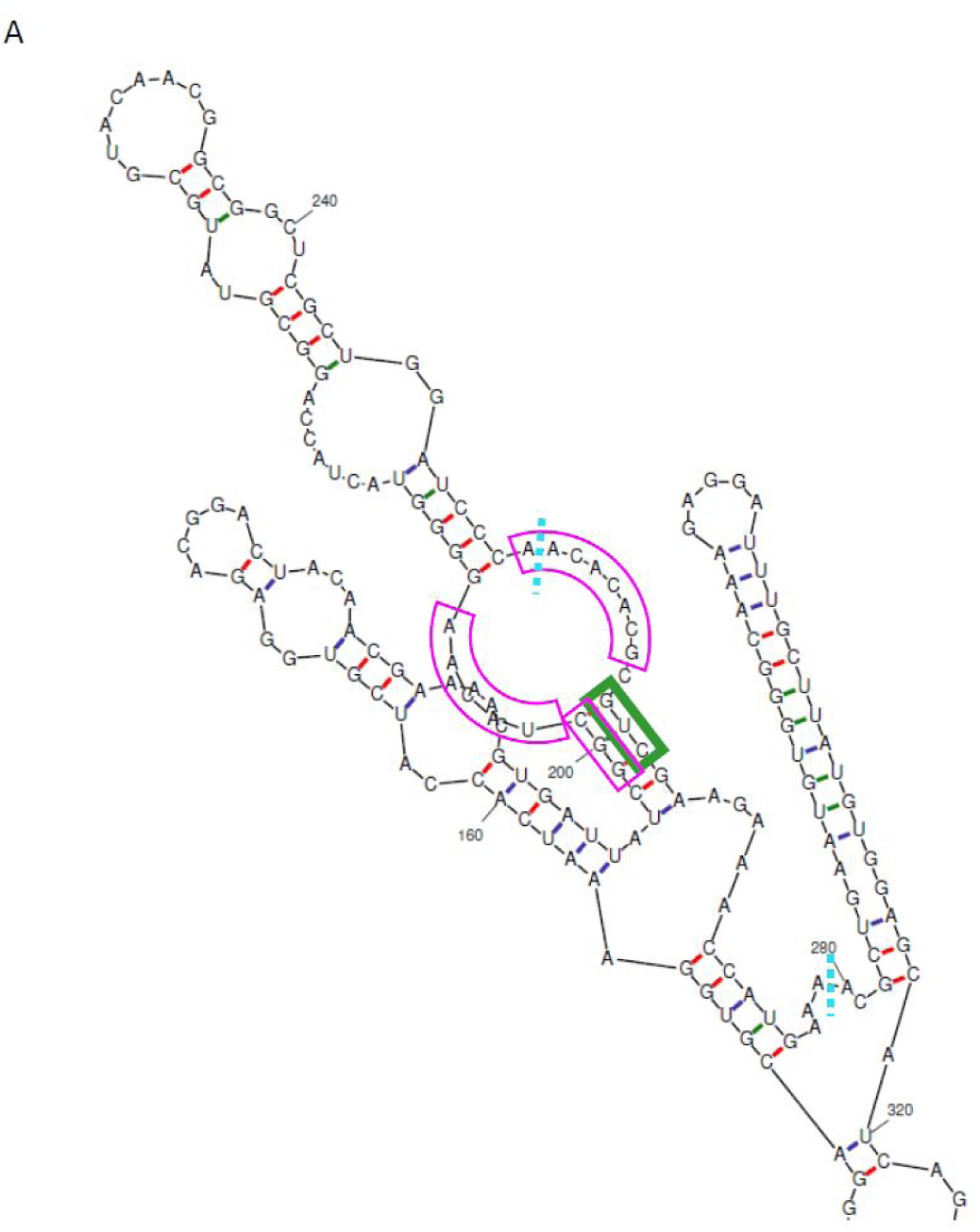
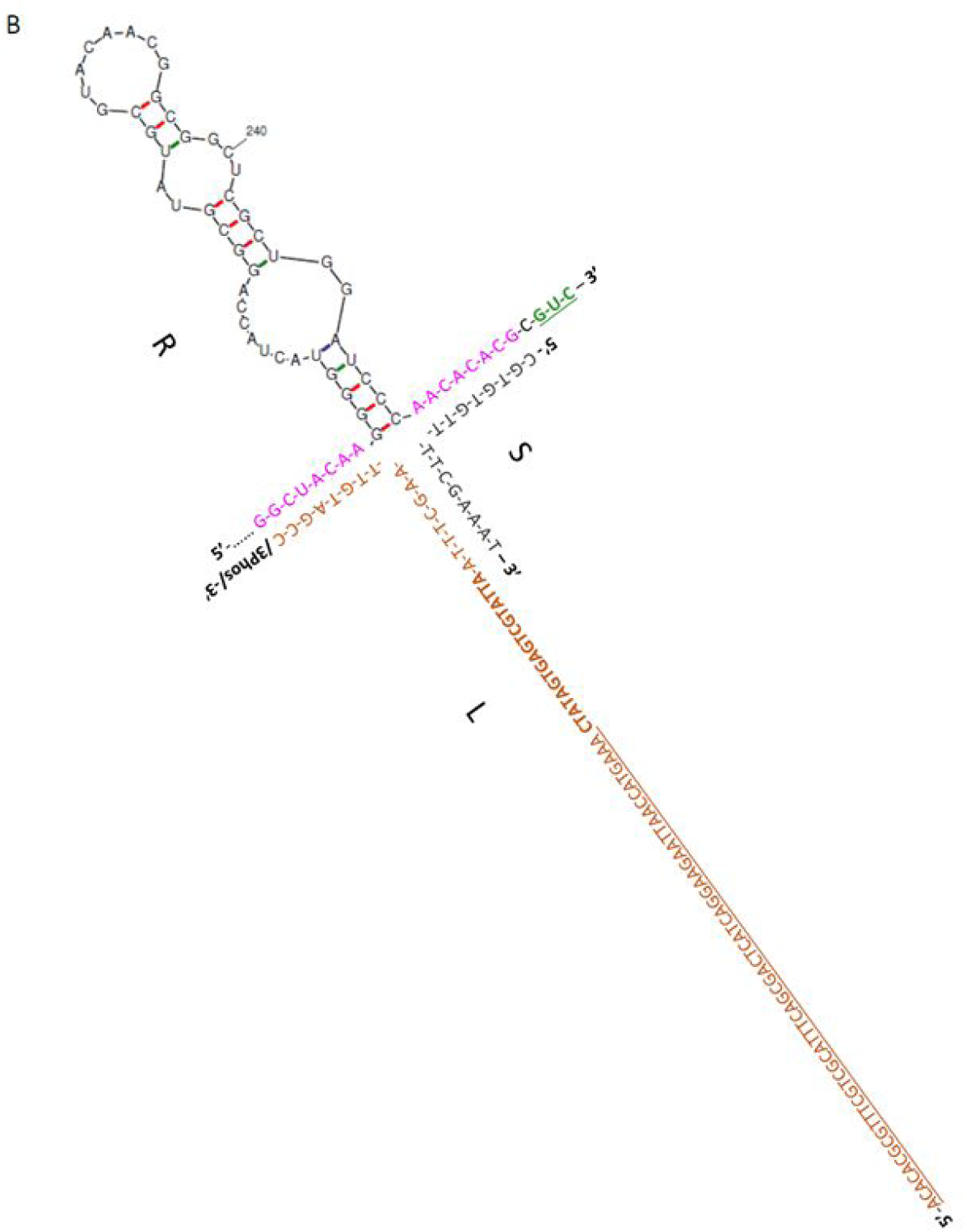
Design of GP64 molecular automaton. R strand as represented is part of the GP64 RNA (positions 199-264b). Cleavage sequence ‘GUC’ is colored in green. **A,** Schematic illustration before cleavage. HHR bending region located between the turquoise lines ACACACGCGUCGAAGAAACCAUGAAA (positions 254-279). The fuchsia marked parts are R(S) and R(L). **B,** Assembled GP64 automaton on cleaved target. S, L and R are represented in gray, brown and black and fuchsia respectively. Fuchsia parts represent R(S) and R(L) regions. HHR and T7 complementary sequences are underlined and bolded accordingly in L strand.

**Figure S25.**
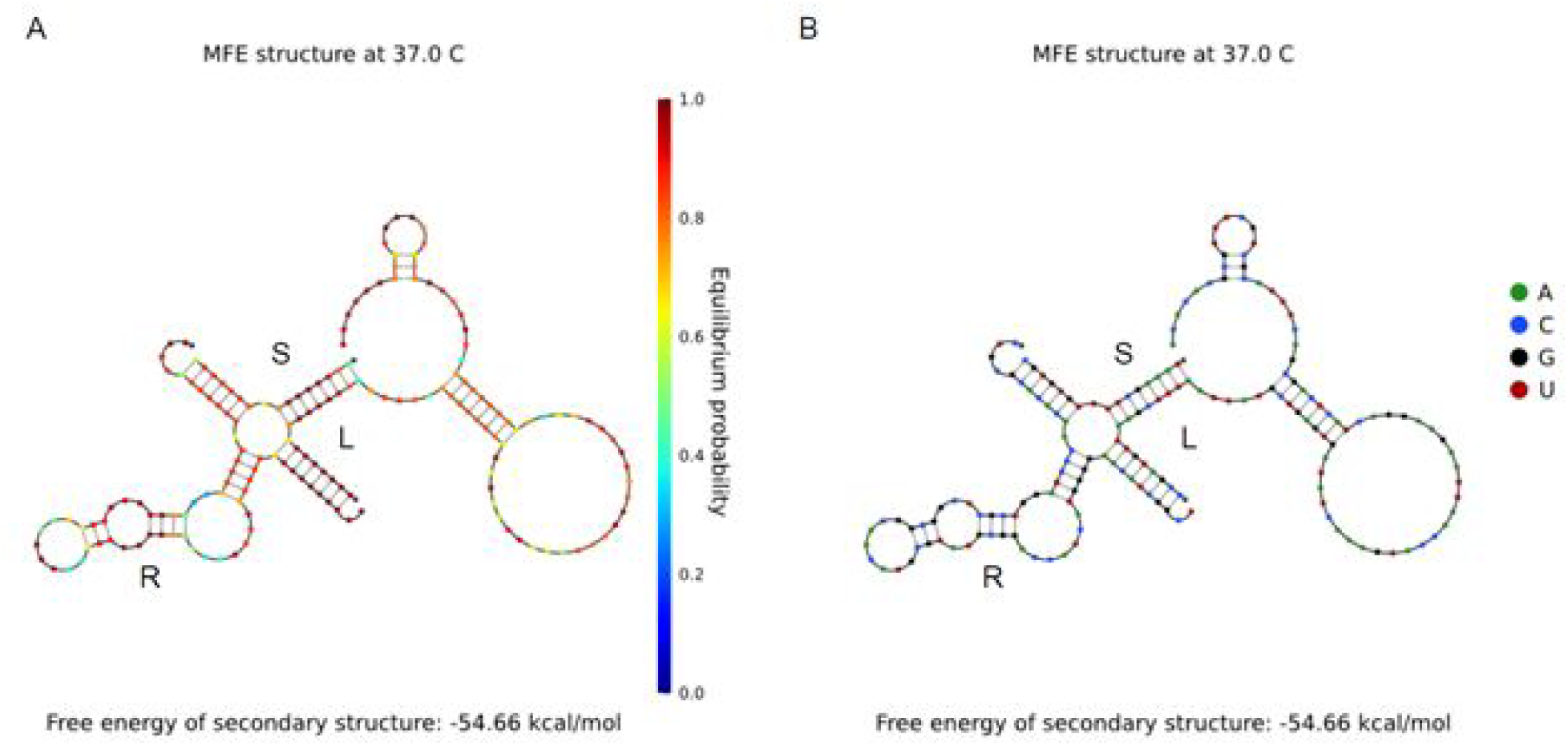
NUPACK assembly simulation of GP64 automaton. R represents bases located in 197-264 positions (UCGGCUACAAGGGGUACUACCAGGCGUAUGCGUACAACGGCGGCUCGCUGGAUCCCAACACACGCGUC), which is part of GP64 RNA that participates in the cleavage and automaton assembly. L-R-S predicted secondary structure at 37° C. L, S and R represent the automaton parts. **A,** Equilibrium probability **B,** Base identity shading.

**Figure S26.**
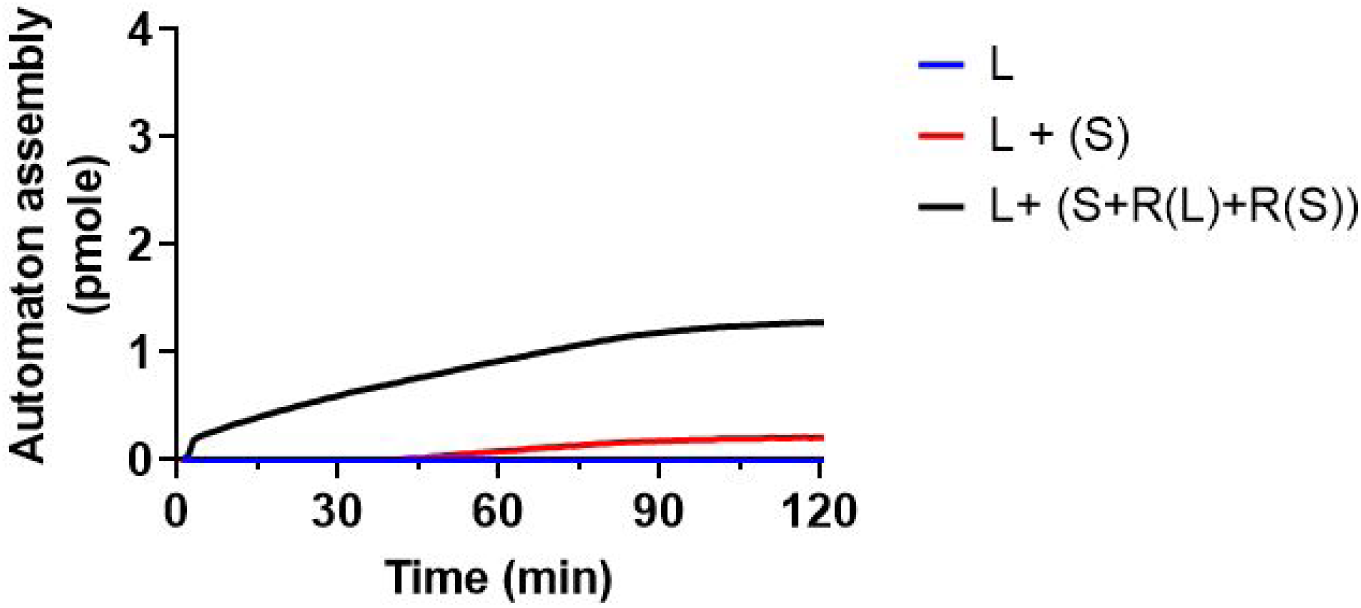
*In-vitro* assembly of GP64 molecular automaton. L strand was labeled with FAM while S with dark FQ. Automaton assembly resulted in fluorescence decrease and converted to moles. The target - R, is required to form a stable hybridization between L and S.

**Table S4.**
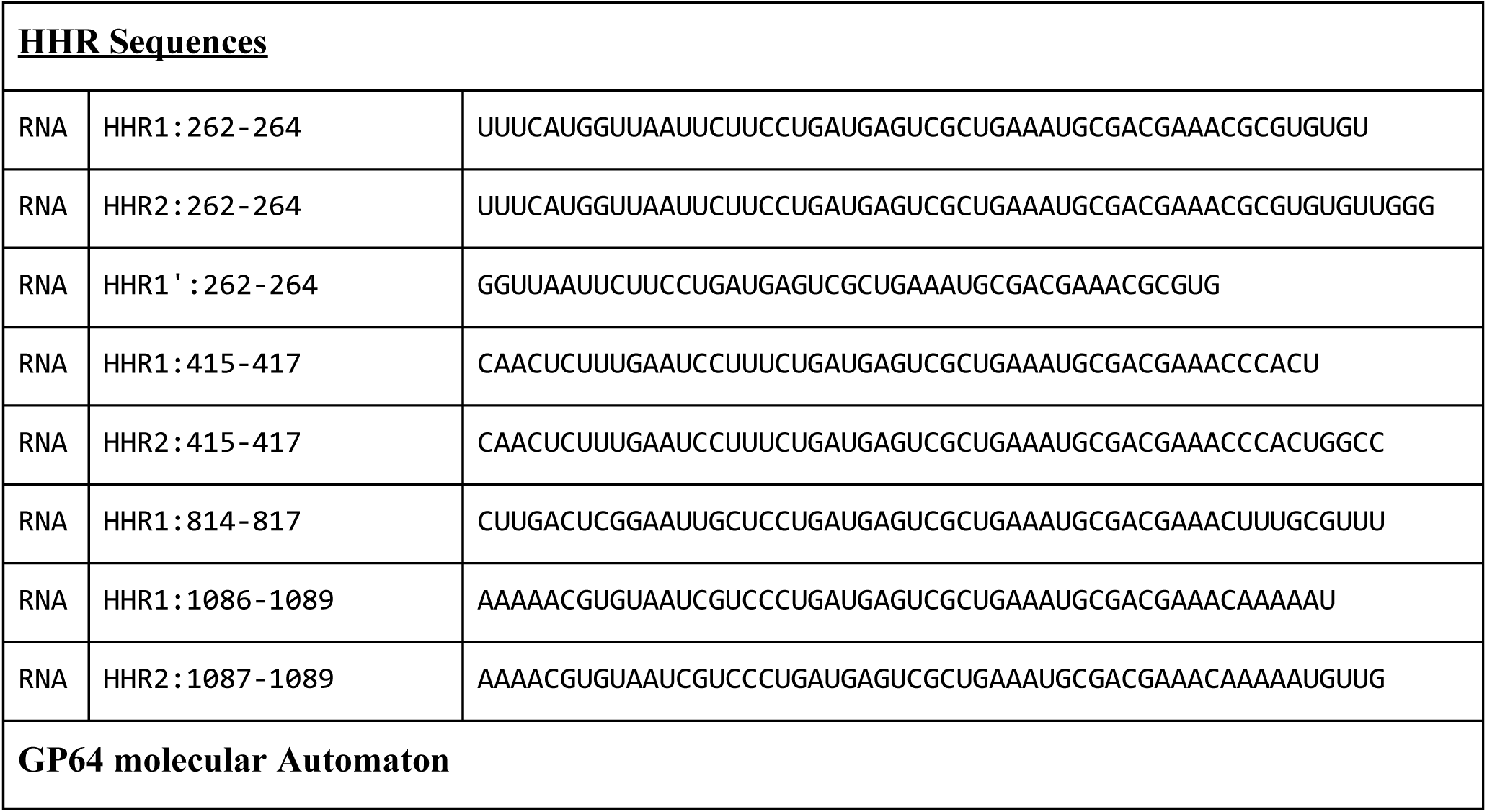
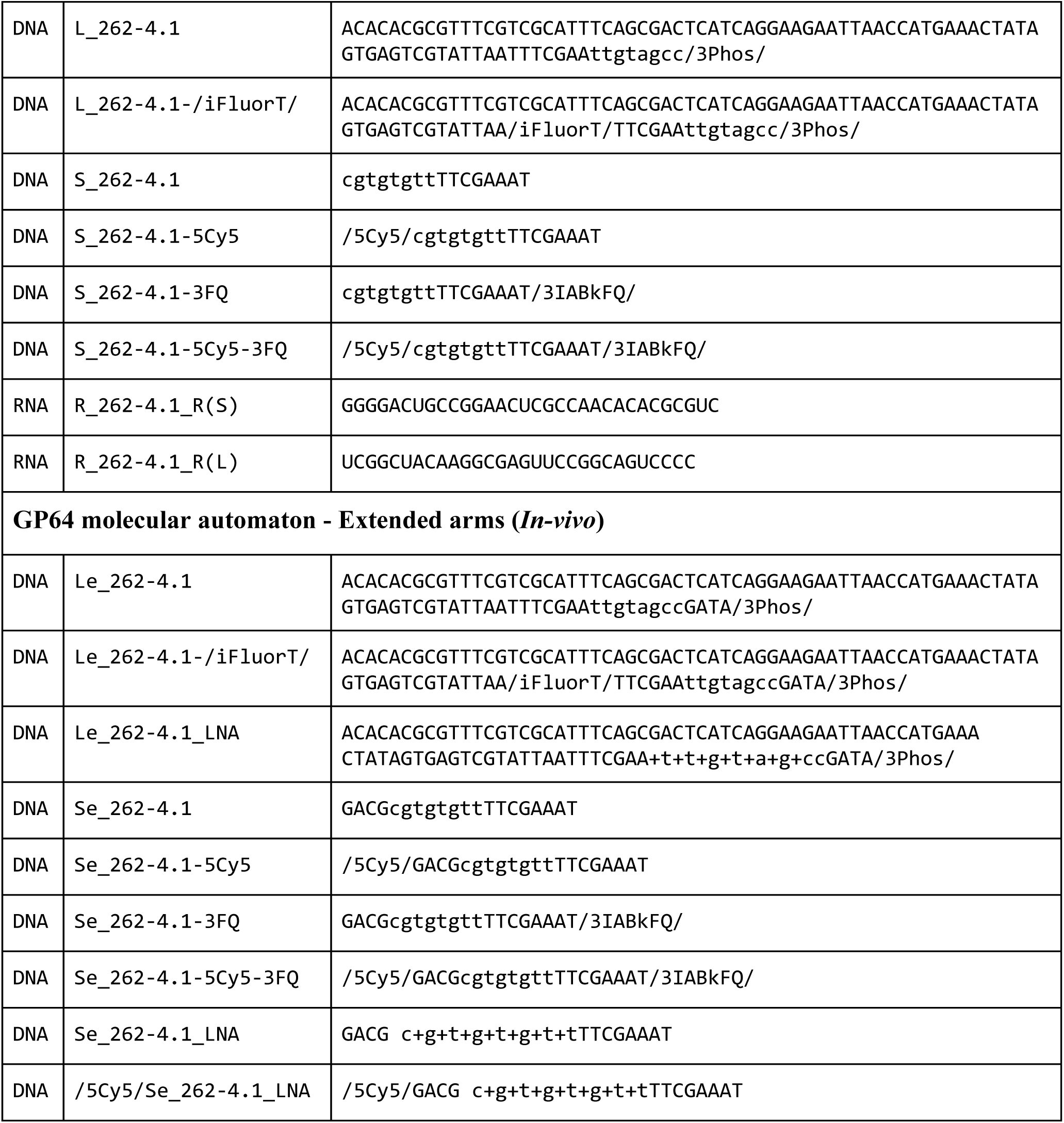
GP64 automaton sequence list

**GP64 CDS (NC_001623.1):**

>lcl|NC_001623.1_cds_NP_054158.1_129 [gene=Ac-GP64] [locus_tag=ACNVgp129] [db_xref=GeneID:1403961] [protein=major budded virus envelope glycoprotein] [protein_id=NP_054158.1] [location=complement(108179..109717)] [gbkey=CDS]

ATGGTAAGCGCTATTGTTTTATATGTGCTTTTGGCGGCGGCGGCGCATTCTGCCTTTGCGGCGGAGCACTGCAACGCGCAAATGA AGACGGGTCCGTACAAGATTAAAAACTTGGACATTACCCCGCCCAAGGAAACGCTGCAAAAGGACGTGGAAATCACCATCGTGGA GACGGACTACAACGAAAACGTGATTATCGGCTACAAGGGGTACTACCAGGCGTATGCGTACAACGGCGGCTCGCTGGATCCCAAC ACACGCGTCGAAGAAACCATGAAAACGCTGAATGTGGGCAAAGAGGATTTGCTTATGTGGAGCATCAGGCAGCAGTGCGAGGTGG GCGAAGAGCTGATCGACCGTTGGGGCAGTGACAGCGACGACTGTTTTCGCGACAACGAGGGCCGCGGCCAGTGGGTCAAAGGCAA AGAGTTGGTGAAGCGGCAGAATAACAATCACTTTGCGCACCACACGTGCAACAAATCGTGGCGATGCGGCATTTCCACTTCGAAA ATGTACAGCAGGCTCGAGTGCCAGGACGACACGGACGAGTGCCAGGTATACATTTTGGACGCTGAGGGCAACCCCATCAACGTGA CCGTGGACACTGTGCTTCATCGAGACGGCGTGAGTATGATTCTCAAACAAAAGTCTACGTTCACCACGCGCCAAATAAAAGCTGC GTGTCTGCTCATTAAAGATGACAAAAATAACCCCGAGTCGGTGACACGCGAACACTGTTTGATTGACAATGATATATATGATCTT TCTAAAAACACGTGGAACTGCAAGTTTAACAGATGCATTAAACGCAAAGTCGAGCACCGAGTCAAGAAGCGGCCGCCCACTTGGC GCCACAACGTTAGAGCCAAGTACACAGAGGGAGACACTGCCACCAAAGGCGACCTGATGCATATTCAAGAGGAGCTGATGTACGA AAACGATTTGCTGAAAATGAACATTGAGCTGATGCATGCGCACATCAACAAGCTAAACAATATGCTGCACGACCTGATAGTCTCC GTGGCCAAGGTGGACGAGCGTTTGATTGGCAATCTCATGAACAACTCTGTTTCTTCAACATTTTTGTCGGACGACACGTTTTTGC TGATGCCGTGCACCAATCCGCCGGCACACACCAGTAATTGCTACAACAACAGCATCTACAAAGAAGGGCGTTGGGTGGCCAACAC GGACTCGTCGCAATGCATAGATTTTAGCAACTACAAGGAACTAGCAATTGACGACGACGTCGAGTTTTGGATCCCGACCATCGGC AACACGACCTATCACGACAGTTGGAAAGATGCCAGCGGCTGGTCGTTTATTGCCCAACAAAAAAGCAACCTCATAACCACCATGG AGAACACCAAGTTTGGCGGCGTCGGCACCAGTCTGAGCGACATCACTTCCATGGCTGAAGGCGAATTGGCCGCTAAATTGACTTC GTTCATGTTTGGTCATGTAGTTAACTTTGTAATTATATTAATTGTGATTTTATTTTTGTACTGTATGATTAGAAACCGTAATAGA CAATATTAA

## References

1. Seeman, N. C. Nucleic acid junctions and lattices. J. Theor. Biol. 99, 237–247 (1982).

2. Wang, T. et al. A DNA Crystal Designed to Contain Two Molecules per Asymmetric Unit Cell. (2011) doi:10.2210/pdb3nao/pdb.

3. Hao, Y. et al. A device that operates within a self-assembled 3D DNA crystal. Nat. Chem. 9, 824–827 (2017).

4. Zheng, J. et al. From molecular to macroscopic via the rational design of a self-assembled 3D DNA crystal. Nature 461, 74–77 (2009).

5. Mao, C., LaBean, T. H., Relf, J. H. & Seeman, N. C. Logical computation using algorithmic self-assembly of DNA triple-crossover molecules. Nature 407, 493–496 (2000).

6. Zhang, F., Simmons, C. R., Gates, J., Liu, Y. & Yan, H. Self-Assembly of a 3D DNA Crystal Structure with Rationally Designed Six-Fold Symmetry. Angew. Chem. Int. Ed Engl. 57, 12504–12507 (2018).

7. He, Y. et al. Hierarchical self-assembly of DNA into symmetric supramolecular polyhedra. Nature 452, 198–201 (2008).

8. Rothemund, P. W. K. Folding DNA to create nanoscale shapes and patterns. Nature 440, 297–302 (2006).

9. Douglas, S. M. et al. Self-assembly of DNA into nanoscale three-dimensional shapes. Nature 459, 414–418 (2009).

10. Wagenbauer, K. F., Sigl, C. & Dietz, H. Gigadalton-scale shape-programmable DNA assemblies. Nature 552, 78–83 (2017).

11. Gerling, T., Wagenbauer, K. F., Neuner, A. M. & Dietz, H. Dynamic DNA devices and assemblies formed by shape-complementary, non-base pairing 3D components. Science 347, 1446–1452 (2015).

12. Han, D., et al. Single-stranded DNA and RNA origami. Science 358, (2017).

13. Ong, L. L. et al. Programmable self-assembly of three-dimensional nanostructures from 10,000 unique components. Nature 552, 72–77 (2017).

14. Schueder, F. et al. Multiplexed 3D super-resolution imaging of whole cells using spinning disk confocal microscopy and DNA-PAINT. Nat. Commun. 8, 2090 (2017).

15. Hübner, K. et al. Directing Single-Molecule Emission with DNA Origami-Assembled Optical Antennas. Nano Lett. 19, 6629–6634 (2019).

16. Nickels, P. C. et al. Molecular force spectroscopy with a DNA origami-based nanoscopic force clamp. Science 354, 305–307 (2016).

17. Bastings, M. M. C. et al. Modulation of the Cellular Uptake of DNA Origami through Control over Mass and Shape. Nano Lett. 18, 3557–3564 (2018).

18. Winfree, E., Liu, F., Wenzler, L. A. & Seeman, N. C. Design and self-assembly of two-dimensional DNA crystals. Nature 394, 539–544 (1998).

19. Adleman, L. M. Molecular computation of solutions to combinatorial problems. Science 266, 1021–1024 (1994).

20. Benenson, Y., Gil, B., Ben-Dor, U., Adar, R. & Shapiro, E. An autonomous molecular computer for logical control of gene expression. Nature 429, 423–429 (2004).

21. Benenson, Y. et al. Programmable and autonomous computing machine made of biomolecules. Nature 414, 430–434 (2001).

22. Samanta, B. & Joyce, G. F. A reverse transcriptase ribozyme. Elife 6, (2017).

23. Horning, D. P. & Joyce, G. F. Amplification of RNA by an RNA polymerase ribozyme. Proc. Natl. Acad. Sci. U. S. A. 113, 9786–9791 (2016).

24. Sczepanski, J. T. & Joyce, G. F. A cross-chiral RNA polymerase ribozyme. Nature 515, 440–442 (2014).

25. Zhang, B. & Cech, T. R. Peptide bond formation by in vitro selected ribozymes. Nature 390, 96–100 (1997).

26. Cuenoud, B. & Szostak, J. W. A DNA metalloenzyme with DNA ligase activity. Nature 375, 611–614 (1995).

27. Ellington, A. D. & Szostak, J. W. In vitro selection of RNA molecules that bind specific ligands. Nature vol. 346 818–822 (1990).

28. Tuerk, C. & Gold, L. Systematic evolution of ligands by exponential enrichment: RNA ligands to bacteriophage T4 DNA polymerase. Science 249, 505–510 (1990).

29. Seelig, G., Soloveichik, D., Zhang, D. Y. & Winfree, E. Enzyme-Free Nucleic Acid Logic Circuits. Science vol. 314 1585–1588 (2006).

30. Qian, L., Winfree, E. & Bruck, J. Neural network computation with DNA strand displacement cascades. Nature 475, 368–372 (2011).

31. Qian, L. & Winfree, E. Scaling up digital circuit computation with DNA strand displacement cascades. Science 332, 1196–1201 (2011).

32. Zhang, D. Y. & Seelig, G. Dynamic DNA nanotechnology using strand-displacement reactions. Nat. Chem. 3, 103–113 (2011).

33. Douglas, S. M. et al. Rapid prototyping of 3D DNA-origami shapes with caDNAno. Nucleic Acids Res. 37, 5001–5006 (2009).

34. Kim, D.-N., Kilchherr, F., Dietz, H. & Bathe, M. Quantitative prediction of 3D solution shape and flexibility of nucleic acid nanostructures. Nucleic Acids Res. 40, 2862–2868 (2012).

35. Zhu, J., Wei, B., Yuan, Y. & Mi, Y. UNIQUIMER 3D, a software system for structural DNA nanotechnology design, analysis and evaluation. Nucleic Acids Research vol. 37 2164–2175 (2009).

36. Lakin, M. R., Youssef, S., Polo, F., Emmott, S. & Phillips, A. Visual DSD: a design and analysis tool for DNA strand displacement systems. Bioinformatics 27, 3211–3213 (2011).

37. Zadeh, J. N. et al. NUPACK: Analysis and design of nucleic acid systems. J. Comput. Chem. 32, 170–173 (2011).

38. Feynman, R. There’s Plenty of Room at the Bottom. Feynman and Computation 63–76 doi:10.1201/9780429500459-7.

39. Eric Drexler, K. Engines of Creation. (Anchor Books, 1986).

40. Asimov, I. Fantastic Voyage: A Novel. (Bantam, 2011).

41. Schwartz, J. T., von Neumann, J. & Burks, A. W. Theory of Self-Reproducing Automata. Mathematics of Computation vol. 21 745 (1967).

42. Douglas, S. M., Bachelet, I. & Church, G. M. A logic-gated nanorobot for targeted transport of molecular payloads. Science 335, 831–834 (2012).

43. Kaminka, G. A., Spokoini-Stern, R., Amir, Y., Agmon, N. & Bachelet, I. Molecular Robots Obeying Asimov’s Three Laws of Robotics. Artif. Life 23, 343–350 (2017).

44. Amir, Y. et al. Universal computing by DNA origami robots in a living animal. Nat. Nanotechnol. 9, 353–357 (2014).

45. Amir, Y., Abu-Horowitz, A., Werfel, J. & Bachelet, I. Nanoscale Robots Exhibiting Quorum Sensing. Artif. Life 25, 227–231 (2019).

46. Gu, H., Chao, J., Xiao, S.-J. & Seeman, N. C. A proximity-based programmable DNA nanoscale assembly line. Nature 465, 202–205 (2010).

47. Lund, K. et al. Molecular robots guided by prescriptive landscapes. Nature 465, 206–210 (2010).

48. Li, S. et al. A DNA nanorobot functions as a cancer therapeutic in response to a molecular trigger in vivo. Nat. Biotechnol. 36, 258–264 (2018).

49. Andersen, E. S. et al. Self-assembly of a nanoscale DNA box with a controllable lid. Nature 459, 73–76 (2009).

50. Thubagere, A. J., et al. A cargo-sorting DNA robot. Science 357, (2017).

51. Freitas, R. A. & Merkle, R. C. Kinematic Self-replicating Machines. (Landes Bioscience, 2004).

52. Penrose, L. S. & Penrose, R. A Self-reproducing Analogue. Nature 179, 1183–1183 (1957).

53. Wang, T. et al. Self-replication of information-bearing nanoscale patterns. Nature 478, 225–228 (2011).

54. Kim, J., Lee, J., Hamada, S., Murata, S. & Ha Park, S. Self-replication of DNA rings. Nat. Nanotechnol. 10, 528–533 (2015).

55. Zhuo, R. et al. Litters of self-replicating origami cross-tiles. Proc. Natl. Acad. Sci. U. S. A. 116, 1952–1957 (2019).

56. Codd, E. F. A Self-Reproducing Universal Computer-Constructor. Cellular Automata 81–105 (1968) doi:10.1016/b978-1-4832-0014-9.50010-8.

57. Penrose, L. S. Self-Reproducing Machines. Scientific American vol. 200 105–114 (1959).

58. Osmanov, Z. On the interstellar Von Neumann micro self-reproducing probes. International Journal of Astrobiology 1–4 (2019) doi:10.1017/s1473550419000259.

59. Liu, B. L. et al. ICP34.5 deleted herpes simplex virus with enhanced oncolytic, immune stimulating, and anti-tumour properties. Gene Therapy vol. 10 292–303 (2003).

60. Perri, S. et al. An alphavirus replicon particle chimera derived from venezuelan equine encephalitis and sindbis viruses is a potent gene-based vaccine delivery vector. J. Virol. 77, 10394–10403 (2003).

61. [No title]. http://symposium.cshlp.org/content/11/local/back-matter.pdf.

62. Kharma, N. et al. Automated design of hammerhead ribozymes and validation by targeting the PABPN1 gene transcript. Nucleic Acids Res. 44, e39 (2016).

63. O’Rourke, S. M. & Scott, W. G. Structural Simplicity and Mechanistic Complexity in the Hammerhead Ribozyme. Prog. Mol. Biol. Transl. Sci. 159, 177–202 (2018).

64. Elbaz, J., Yin, P. & Voigt, C. A. Genetic encoding of DNA nanostructures and their self-assembly in living bacteria. Nat. Commun. 7, 11179 (2016).

65. Delebecque, C. J., Lindner, A. B., Silver, P. A. & Aldaye, F. A. Organization of intracellular reactions with rationally designed RNA assemblies. Science 333, 470–474 (2011).

66. Zhao, S. et al. Efficient Intracellular Delivery of RNase A Using DNA Origami Carriers. ACS Appl. Mater. Interfaces 11, 11112–11118 (2019).

67. Schaffert, D. H. et al. Intracellular Delivery of a Planar DNA Origami Structure by the Transferrin-Receptor Internalization Pathway. Small 12, 2634–2640 (2016).

68. Palazzolo, S. et al. An Effective Multi-Stage Liposomal DNA Origami Nanosystem for In Vivo Cancer Therapy. Cancers 11, (2019).

69. Perrault, S. D. & Shih, W. M. Virus-inspired membrane encapsulation of DNA nanostructures to achieve in vivo stability. ACS Nano 8, 5132–5140 (2014).

70. Blissard, G. W. & Theilmann, D. A. Baculovirus Entry and Egress from Insect Cells. Annu Rev Virol 5, 113–139 (2018).

71. Xu, J. et al. ERK signaling couples nutrient status to antiviral defense in the insect gut. Proc. Natl. Acad. Sci. U. S. A. 110, 15025–15030 (2013).

72. Zhou, P. et al. A pneumonia outbreak associated with a new coronavirus of probable bat origin. Nature 579, 270–273 (2020).

73. Asian Development Bank. Developing Asia Growth to Fall in 2020 on COVID-19 Impact. Asian Development Bank https://www.adb.org/news/developing-asia-growth-fall-2020-covid-19-impact (2020).

74. Thanh Le, T. et al. The COVID-19 vaccine development landscape. Nat. Rev. Drug Discov. (2020) doi:10.1038/d41573-020-00073-5.

75. Gilbert, W. Origin of life: The RNA world. Nature vol. 319 618–618 (1986).

76. Striggles, J. C., Martin, M. B. & Schmidt, F. J. Frequency of RNA-RNA interaction in a model of the RNA World. RNA 12, 353–359 (2006).

77. Attwater, J., Raguram, A., Morgunov, A. S., Gianni, E. & Holliger, P. Ribozyme-catalysed RNA synthesis using triplet building blocks. eLife vol. 7 (2018).

78. Wachowius, F. & Holliger, P. Non-Enzymatic Assembly of a Minimized RNA Polymerase Ribozyme. ChemSystemsChem vol. 1 12–15 (2019).

79. McCaskill, J. S. Spatially resolved in vitro molecular ecology. Biophys. Chem. 66, 145–158 (1997).

80. Wlotzka, B. & McCaskill, J. S. A molecular predator and its prey: coupled isothermal amplification of nucleic acids. Chem. Biol. 4, 25–33 (1997).

81. Figueroa-Bossi, N. & Bossi, L. Sponges and Predators in the Small RNA World. Microbiol Spectr 6, (2018).

82. Nutiu, R. & Li, Y. Structure-switching signaling aptamers. J. Am. Chem. Soc. 125, 4771–4778 (2003).

83. Lieberman, J. Tapping the RNA world for therapeutics. Nat. Struct. Mol. Biol. 25, 357–364 (2018).

84. Kole, R., Krainer, A. R. & Altman, S. RNA therapeutics: beyond RNA interference and antisense oligonucleotides. Nat. Rev. Drug Discov. 11, 125–140 (2012).

85. Mamet, N. et al. Discovery of tumoricidal DNA oligonucleotides by response-directed in vitro evolution. Commun Biol 3, 29 (2020).

86. Hacohen, A., Cohen, R., Efroni, S., Barzel, B. & Bachelet, I. Digitizable therapeutics for decentralized mitigation of global pandemics. Sci. Rep. 9, 14345 (2019).

87. Primer designing tool. https://www.ncbi.nlm.nih.gov/tools/primer-blast/.

88. Bulmer, M. S., Bachelet, I., Raman, R., Rosengaus, R. B. & Sasisekharan, R. Targeting an antimicrobial effector function in insect immunity as a pest control strategy. Proc. Natl. Acad. Sci. U. S. A. 106, 12652–12657 (2009).

89. Zuker, M. Mfold web server for nucleic acid folding and hybridization prediction. Nucleic Acids Research vol. 31 3406–3415 (2003).

